# Resource Allocation in Annual Plants

**DOI:** 10.1101/2021.04.19.440512

**Authors:** David McMorris, Glenn Ledder

## Abstract

The fitness of an annual plant can be thought of as how much fruit is produced by the end of its growing season. Under the assumption that annual plants grow to maximize fitness, we can use techniques from optimal control theory to understand this process. We introduce a model for resource allocation in annual plants which extends classical work by Iwasa and Roughgarden to a case where both carbohydrates and mineral nutrients are allocated to shoots, roots, and fruits in annual plants. We use optimal control theory to determine the optimal resource allocation strategy for the plant throughout its growing season as well as develop a numerical scheme to implement the model in MATLAB. Our results suggest that what is optimal for an individual plant is highly dependent on initial conditions, and optimal growth has the effect of driving a wide range of initial conditions toward common configurations of biomass by the end of a growing season.

## 1 Introduction

Plant life history theory is generally concerned with the strategies plants employ for survival and reproduction, as well as how these processes influence population dynamics. The question of how plants allocate resources, e.g. C, N, P, and in some models, biomass, and what drives these allocation rules, is central to this pursuit. There are several schools of thought which seek to provide a framework for understanding these allocation patterns. One school of thought suggests that allocation rules should ultimately be the result of natural selection, and so resource allocation should optimize fitness in some sense (see Iwasa and Roughgarden 1984; Velten and Richter 1995). Another framework views biomass allocation as following certain allometric scaling relationships (see McCarthy and Enquist 2007; Enquist and Niklas 2002), and yet another views allocation not through the lens of an individual organism or specific genome, but rather from a game-theoretical perspective in which allocation rules are driven by competition, and follow an evolutionary stable strategy (see Dybzinski et al. 2011). It has also been suggested that allocation may be an emergent property of the many processes involved in plant growth rather than the result of a central allocation process (Feller et al. 2015). For a more complete review of allocation theory, we refer the reader to Ledder et al. (2020) or Poorter et al. (2012).

In this paper, we take the viewpoint that resource allocation, in annual plants specifically, should serve to optimize overall fitness. It’s important to note that, while patterns of growth consistent with optimal allocation have been observed to some extent (see e.g. McCarthy and Enquist 2007), even if this is not universally true it is still important to have a theory of optimal growth for comparison with observed behavior. Whereas previous work has focused primarily on optimal allocation of a single resource, be it carbon or biomass, and ignore the role of mineral nutrients, our work seeks to develop a theory that acknowledges the importance of both carbon and mineral nutrients. This is in line with the functional equilibrium hypothesis, which states that optimal growth occurs when resources are allocated in such a way that no single resource is any more limiting than any other (see Bloom et al. 1985; Weiner 2004; Poorter and Nagel 2000).

This work is primarily an extension of classical work by Iwasa and Roughgarden (1984), who considered a model for photosynthate (C) allocation in annual plants. They assumed that photosynthate production depends on both shoots and roots, and that allocation is split between shoots, roots, and fruits so as to maximize fruit yield. They used optimal control theory to determine that fruit yield is maximized by a three-phase growth pattern, characterized by an initial phase of shoot-only or root-only growth, a period of ‘balanced growth’ during which shoots and roots grow simultaneously, and ultimately a period of fruit-only growth at the end of the growing season.

We present a model which extends the work of Iwasa and Roughgarden to a case which directly incorporates the role of mineral nutrients. In particular, we model the allocation of carbohydrates and mineral nutrients in an annual plant with the objective of optimizing fruit yield, and use optimal control theory to determine the allocation strategies. In this paper we will consider a case where shoots and roots require both resources, and, for simplicity, fruits only require carbohydrates, as in Iwasa and Roughgarden (1984). This simplification allows us to obtain mathematical results which provide an important ‘proof of concept’ for the viability of this type of model, and which we intend to use in the future to guide more general two-resource models.

We find, among other things, that this addition results in four phases of growth, rather than three as seen in Iwasa and Roughgarden (1984). This additional phase consists of a period of mixed vegetative/reproductive growth, during which the fruits and roots grow simultaneously. Furthermore, our results indicate that what is optimal for one plant may not be optimal for another, and optimal growth is largely dependent on initial conditions. We also conclude that this range of optimal strategies may have the effect of reducing population-level variation, thus driving a population toward common sizes and optimal yields. The question remains, however, as to whether plants actually have this degree of strategic plasticity.

## 2 Optimal Control Theory

We will use the notation **x**(*t*) = (*x*_1_(*t*), *x*_2_(*t*), …, *x_n_*(*t*)) for the function 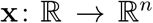 and **g**(*t*, **x**, **u**) = (*g*_1_(*t*, **x**, **u**), *g*_2_(*t*, **x**, **u**), …, *g_n_*(*t*, **x**, **u**)) for the function 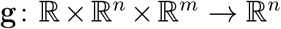. Of particular importance to us are a well-known class of optimal control problems of the form

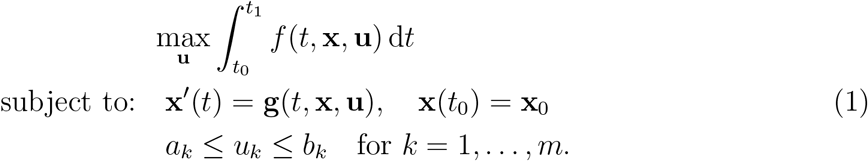

Here we form a Hamiltonian with *n* adjoints λ*_i_*, one for each of the states *x_i_*:

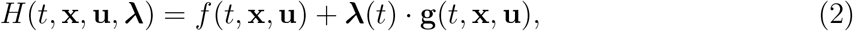

where each adjoint satisfies the following:

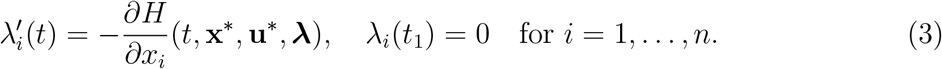

An optimal pair (**x**\*, **u**\*) is characterized by a set of necessary conditions for optimality (4). We refer the reader to Lenhart and Workman (2007) for a thorough treatment of this class of optimal control problems.

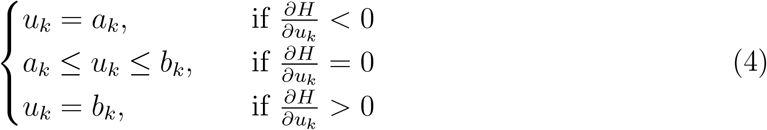

The control problem we will study is an extension of this class of problems, with additional constraints on the controls. We will discuss how the necessary conditions change when they are introduced, as well as include the derivations in Appendix A.

## 3 A Description of the Model

We now turn our attention to the model for resource allocation in annual plants we will discuss as an extension of the work of Iwasa and Roughgarden (1984) to a scenario where the growth of the roots and shoots relies on two resources, rather than one. For the sake of convenience, we will use the terms ‘carbon’ and ‘nitrogen’ to refer to more complicated classes of carbohydrates and mineral nutrients. As a simplifying assumption, we assume that fruits only require carbon for growth.

This assumption makes the mathematical analysis more feasible, and aligns biologically with plants that encase small seeds in large carbon-rich fruits. While this may match our colloquial definition of fruits, e.g. watermelons, it is not an accurate representation of many types of annual plants for which the term ‘fruits’ refers to seeds, nuts, etc., and for which nitrogen content is not negligible.

### 3.1 Model Setup

We consider an annual plant with three organs - shoots, roots, and fruits. Shoots consist of all above-ground vegetative biomass and roots all below-ground vegetative biomass. Fruits refer to any reproductive biomass, be it in the form of colloquial ‘fruits’, nuts, seeds, etc. Biomass of each organ is measured in units of ‘carbon’ where we used ‘carbon’ as a catchall term for carbohydrates produced by the shoots. The functions *S*(*t*), *R*(*t*), and *F*(*t*) give the biomass of shoots, roots, and fruits, respectively, at time *t* throughout a fixed growing season [0, *T*]. We assume that the plant relies on two resources, which we refer to as ‘carbon’ and ‘nitrogen,’ though as previously mentioned we use these terms loosely to refer to more complicated classes of carbohydrates and soil nutrients. We assume that carbon is fixed by the shoots at a rate of *C*(*S*) and nitrogen is absorbed by the roots at a rate of *N*(*R*). Note that this is choice we made to simplify the model, as, in reality, the rate of carbon fixation depends on both shoots and roots via transpiration. Throughout the growing season, fractions of carbon *u_SC_*(*t*), *u_RC_*(*t*), and *u_FC_*(*t*) and fractions of nitrogen *u_SN_*(*t*), *u_RN_*(*t*), and *u_FN_*(*t*) are allocated to the shoots, roots, and fruits, respectively, at time *t*. The resources pass through a synthesizing unit (SU) in each organ, where they are converted into biomass. We use Kooijman’s parallel complementary synthesizing unit (PCSU) function from Kooijman (2010), employing the same simplification seen in (Ledder et al. 2020, Appendix A), given by

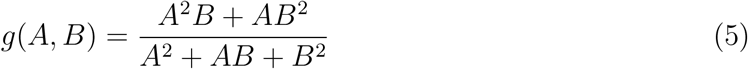

to provide the rate of tissue production when resources are provided at rates *A* and *B*. We can view these rates as representing the maximum rate of tissue production given full utilization of that particular resource, and so when resources are available in the stoichiometric ratio we have that *A* = *B*. As the tissue is measured in units of carbon, we need conversion factors *ν_S_*, *ν_R_*, and *ν_F_*, which give the fixed stoichiometric C : N ratios in the shoots, roots, and fruits, respectively. We specify initial conditions *S*(0) = *S*_0_, *R*(0) = *R*_0_, and *F*(0) = 0 which leads to the following model:

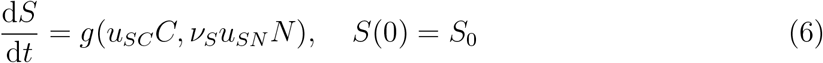

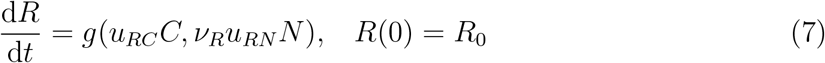

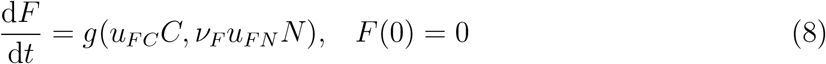

As we are considering the case where fruits only require carbon, we implement this assumption by taking *ν_F_* → ∞, and note that for any nonzero *u_FN_N*, we have the limit

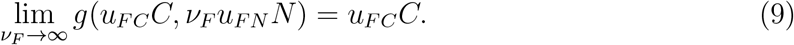

This leads to the following simplified model, which will form the basis for our analysis.

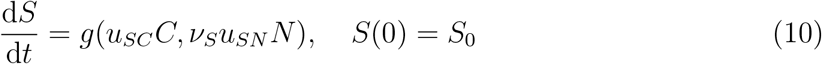

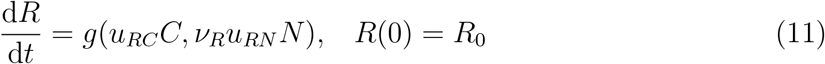

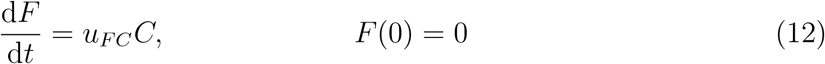

Note that we suppress the arguments in most functions for convenience. This model is shown schematically in Figure 1,

**Figure 1:**
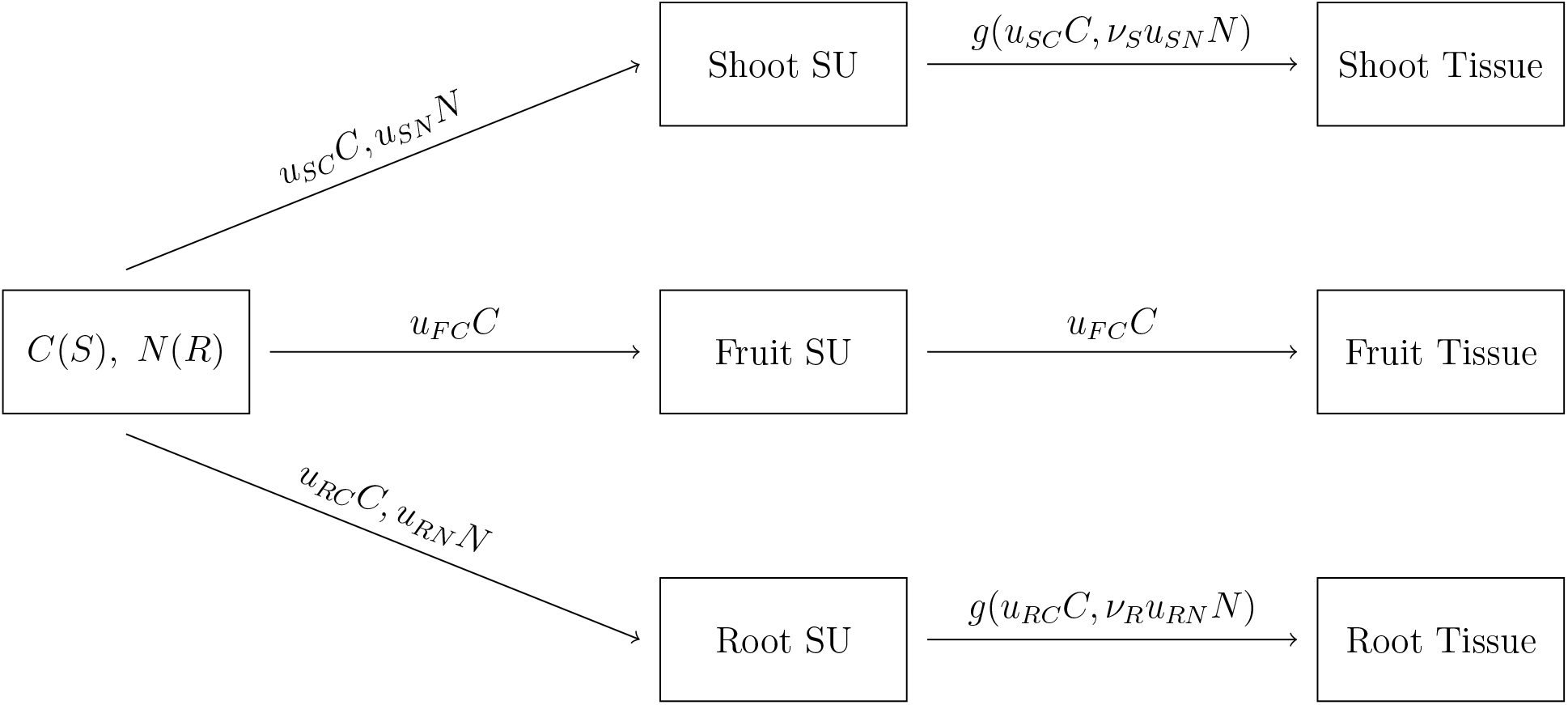
Schematic for model described by equations 10, 11, and 12

Because the functions *u_FC_*, *u_SC_*, *u_RC_*, *u_SN_*, and *u_RN_* represent fractions of carbon and nitrogen, respectively, we impose several restrictions. First, we require that each function be piecewise continuous and bounded between 0 and 1. Furthermore, as we assume full utilization of each resource, we assume that for all times *t* ∈ [0, *T*]

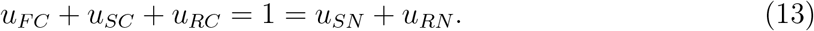

Additionally, to be biologically realistic, we assume that the plant’s capacity to ‘collect’ resources increases continuously with biomass, meaning that we require both *C* and *N* to be continuously differentiable, and

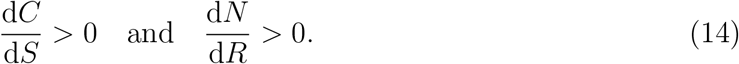

We also assume that the plant experiences possibly diminishing returns with increased biomass, meaning that

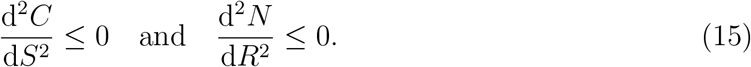

### 3.2 The PCSU

Before we introduce the optimal control framework we will use for determining the optimal resource allocation strategy, there are several important features of the PCSU that merit discussion. As introduced in (5), for resource fluxes *A* and *B*, the PCSU function

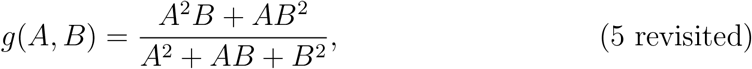

gives the rate of tissue production. Note that when only one resource is present, the tissue production rate is zero as *g*(*A*, 0) = 0 = *g*(0, *B*) for non-zero *A* and *B*. Additionally, this is still true when neither resource is present. In particular, writing (5) in polar coordinates allows us to show that *g*(*A*, *B*) → 0 as (*A*, *B*) → (0, 0). Because of this, we will take *g* to be the continuous extension of this function to the origin such that *g*(0, 0) = 0.

Now, by changing variables to either 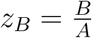 or 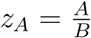, we can rewrite *g* as

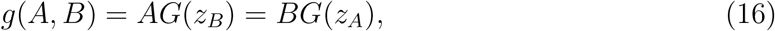

where

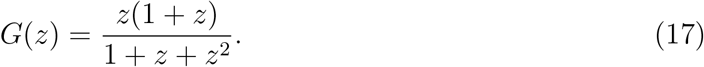

The fact that we can represent the PCSU function in this manner will aid in our analysis by restricting the nonlinearities present to a function of a single variable. By the above discussion, we observe that although *z_B_* and *z_A_* may be undefined when either *A* or *B* or both are zero, both *AG*(*z_B_*) and *BG*(*z_A_*) are zero when at least one of *A* or *B* is zero. It is important to note, however, that whereas *g* is continuous at the origin, *G*(*z_B_*) and *G*(*z_A_*) are not. Additionally, it will facilitate later analysis of the model to note that

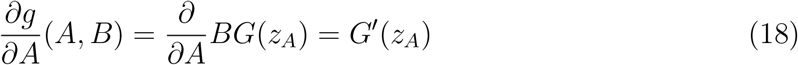

so long as (*A*, *B*) ≠ (0, 0) and, recalling that *g*(0, 0) = 0,

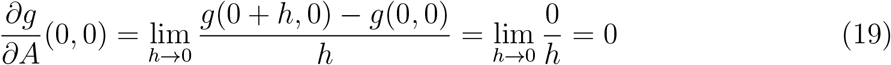

although neither *G*′(*z_B_*) nor *G*′(*z_A_*) is continuous at the origin. We can obtain 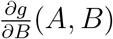 in the same manner, and so we have

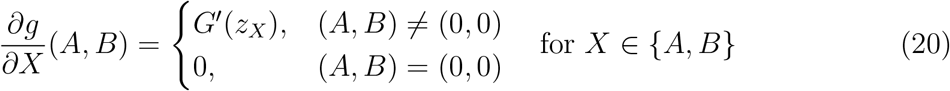

where

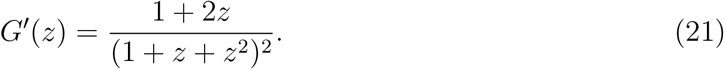

Finally, it is worth noting that both *G* and *G*′ are bounded between 0 and 1 and that

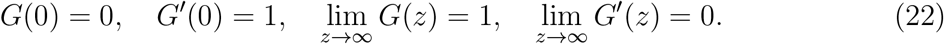

#### 3.2.1 PCSU Identities

There are two identities related to the PCSU function and its derivatives that will be repeatedly cited in later analysis. We will start with the following notation:

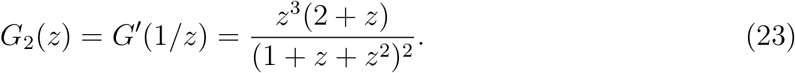

Employing (23), we have the following. We omit the proofs as each can be verified directly.

##### PCSU Identity 1

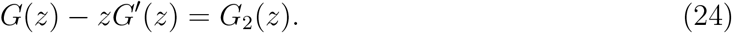

##### PCSU Identity 2

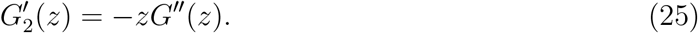

### 3.3 Optimal Control Problem

In this section, we will translate the original problem of finding the growth trajectory to maximize fruit yield to an optimal control problem. Noting that

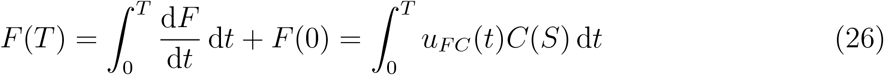

we can reframe our goal as maximizing the right hand side of (26). By imposing the previously discussed constraints that the fractions of each resource, henceforth referred to as the controls, be bounded in [0,1] and fractions of the same resource sum to unity, along with the differential equations for the biomass of each vegetative organ, we can associate the optimal growth trajectory with the solution to the following optimal control problem.

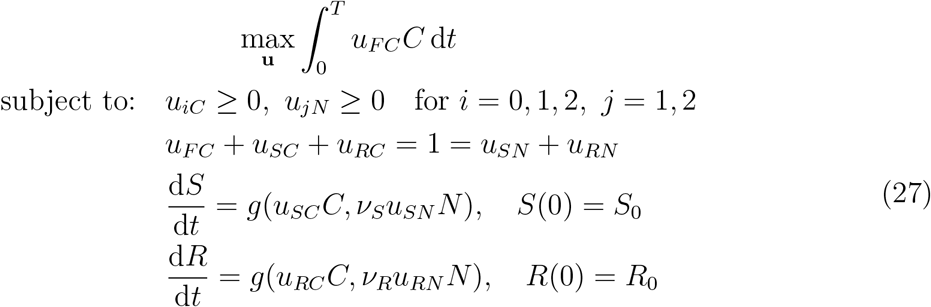

Note that here we only require that the controls be non-negative, because implicit in the combination of those constraints and the equality constraints is the requirement that each control must also be less than one.

### 3.4 Necessary Conditions

We will solve the optimal control problem (27) using a set of necessary conditions that must be satisfied by the solution. The presence of the two equality constraints makes this problem non-standard, so we take the time to derive the necessary conditions for this type of problem. This derivation is found in Appendix A. Since we can essentially think of the carbon controls and the nitrogen controls separately, the necessary conditions for (27) are a combination of the conditions for the two types of problems discussed in Appendix A. We begin by forming a Hamiltonian with two piecewise differentiable adjoints, λ*_S_*(*t*) and λ*_R_*(*t*):

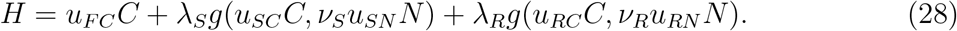

The necessary conditions for optimality are as follows:

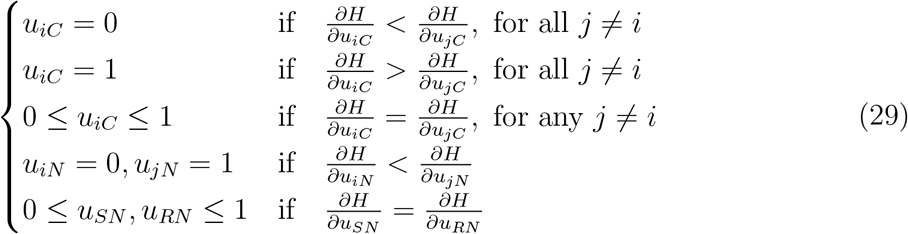

What this essentially says is that all the carbon is being allocated to the organ such that the partial derivative of the Hamiltonian with respect to that organ’s carbon control is larger than the partial derivatives of the Hamiltonian with respect to the other two carbon controls. Likewise for the nitrogen controls. Note also that the Hamiltonian must be constant along the optimal trajectory because the optimal control problem (27) is autonomous (Lenhart and Workman 2007).

As mentioned in Section 3.2, we can rewrite the PCSU function *g* by a change of variables. In particular, we can rewrite (10), (11), and (12) via (16) and the substitutions

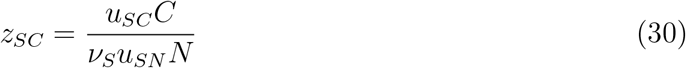

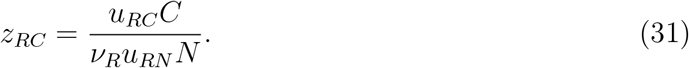

This results in the following system

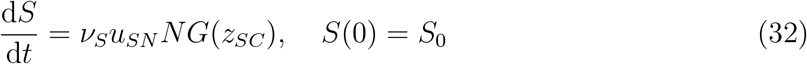

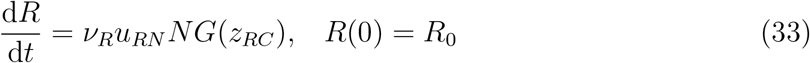

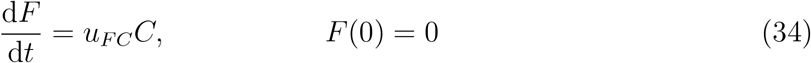

where *G* is given by (17). Therefore, the Hamiltonian can be rewritten as

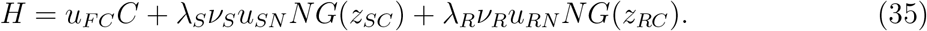

Now, as we previously mentioned, we can essentially think of the carbon controls and nitrogen controls separately. The above formulation provides an avenue for restricting the appearance of the carbon controls to the argument of *G* by letting us write

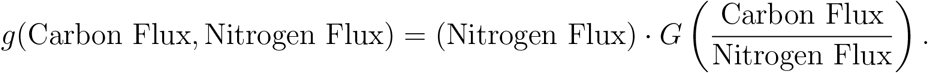

When working directly with the nitrogen controls, it will be useful to write

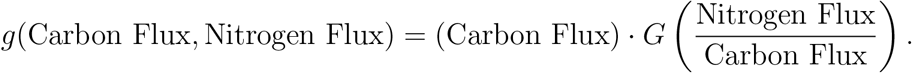

With this in mind, we can make the substitutions

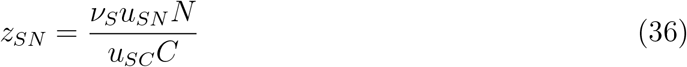

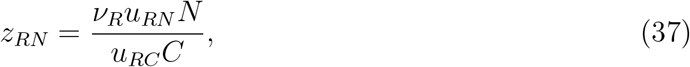

and rewrite (10), (11), and (12) as

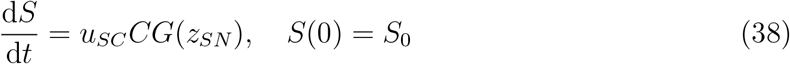

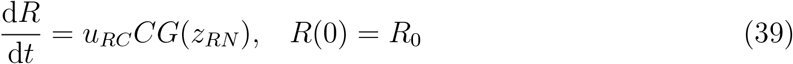

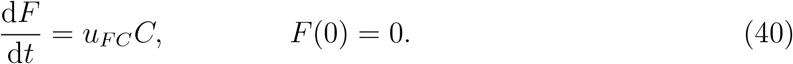

As before, the Hamiltonian can be rewritten as

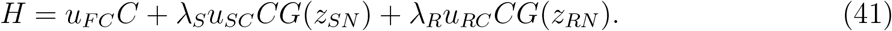

Using (35) and (41), we can compute the following partial derivatives:

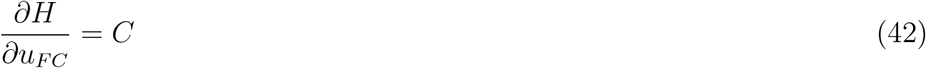

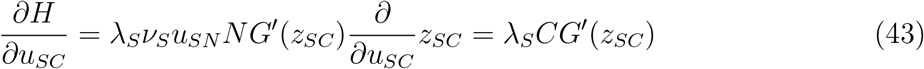

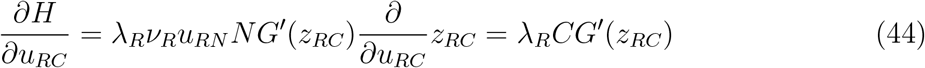

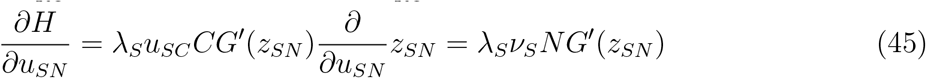

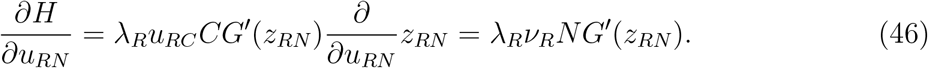

It is important to recall that by (20), these formulations of 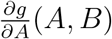 and 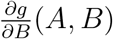 are only valid so long as (*A*, *B*) ≠ (0, 0), in which case these partial derivatives vanish.

The last piece of the optimal control framework concerns the adjoints λ*_S_* and λ*_R_*. By the derivation of the necessary conditions in Appendix A, we have that

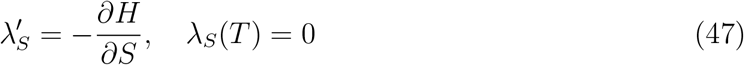

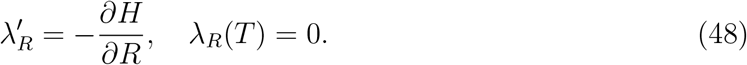

Making use of (35) and (41) we can express (47) and (48) as follows.

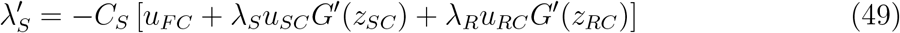

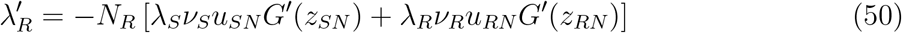

## 4 Four-Phase Structure

In this section we will discuss the structure of the solution to (27). Generally speaking, the solution exhibits a four-phase structure. First, there is an initial phase of vegetative growth in which the plant addresses deficiencies in either shoots or roots. Second, the plant undergoes a phase of balanced growth when both the shoots and the roots are growing. Third, there is a phase of mixed vegetative/reproductive growth, during which the shoots continue to grow and the fruits begin growing simultaneously. The plant completes the growing season with a period of reproductive growth, during which only the fruits are growing.

We will begin with the analysis of the four-phase structure of the optimal solution. Because we know that the adjoints, λ*_S_* and λ*_R_*, vanish at the end of the growing season, we use that as the starting point for our analysis and proceed in reverse.

### 4.1 Final Interval - Reproductive Growth

We begin by showing that there exists a switching time after which all carbon is allocated to fruit production. We will refer to the interval between the switching time and *T* the final interval. Note that by (47) and (48), we have that λ*_S_*(*T*) = 0 = λ*_R_*(*T*). Furthermore, recall that *G*′ is bounded. Therefore, writing *C*(*S*(*T*)) = *C*\*, we have by (42), (43), and (44) that

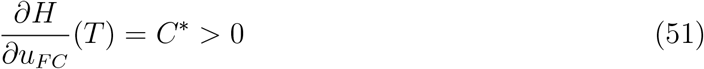

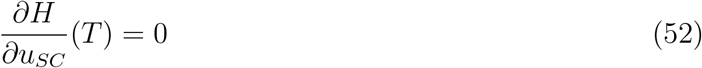

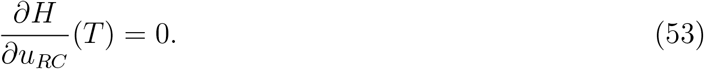

This implies that at *t* = *T*

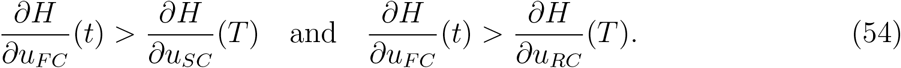

Therefore, by (29), we have that *u_FC_*(*T*) = 1 and *u_SC_*(*T*) = 0 = *u_RC_*(*T*). That is, at the end of the growing season the plant is allocating all of the available carbon to the fruits, which in turn means that only the fruits are growing at this time.

Now, because both λ*_S_* and λ*_R_* are continuous, λ*_S_*(*T*) = 0 = λ*_R_*(*T*), and *G*′ is bounded, there must be some *ε* > 0 such that for all *t* in [*T* – *ε*, *T*] we have that (54) still holds. So, we have the existence of a (potentially small) interval of fruit-only growth. Now, writing 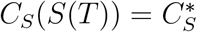, during this interval we have by (49) that

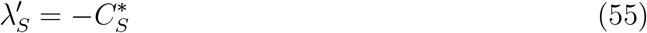

because here we have *u_FC_* = 1, *u_SC_* = 0 = *u_RC_* and *G*′ is bounded. Because λ*_S_*(*T*) = 0 this implies that during this interval

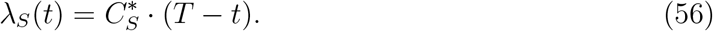

Additionally, because during this interval we have a scenario where for *i* = 1, 2 either only *u_iC_* is zero or both *u_iC_* and *u_iN_* are zero, we have by (50) that

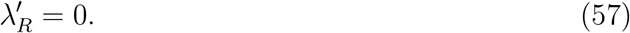

This is because either *u_iC_* alone is zero, in which case *z_iN_* → ∞ and by (22) we have that *G*′(*z_iN_*) → 0, or in the case that both *u_iC_* and *u_iN_* are zero then *u_iN_G*′(*z_iN_*) = 0 because *G*′ is bounded. In either case, we get (57). So, because λ*_R_*(*T*) = 0, we have that the following holds throughout the interval:

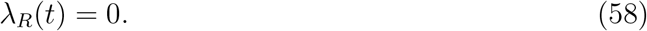

Therefore, because *G*′ is bounded, we have by (46) that

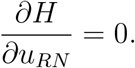

Since (45) is non-negative in this interval, it must be the case that

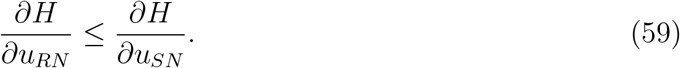

If this inequality were strict, then by (29) we would have *u_SN_* = 1. In this case, because *u_SC_* = 0, we have that *z_SN_* → ∞, and so by (22) we have that *G*′(*z_SN_*) → 0. This, however, implies that

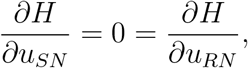

and so the inequality (59) cannot be strict, and both partial derivatives are zero in this interval. In particular, *G*′ (*z_SN_*) = 0, which means that *z_SN_* → ∞ and so *z_SC_* = 0, which by (22) implies that *G*′(*z_SC_*) = 1. In summary then, during this interval we have

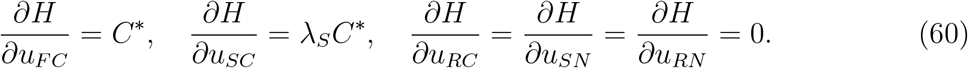

Now, since λ*_S_* is decreasing by (55), we see from (60) that the partial derivatives will maintain the same ordering as long as λ*_S_* < 1. We define the switching point *t*\* to be the time when the plant switches to fruit-only growth. We can identify the switching point as the time when λ*_S_*(*t*) = 1, and we can use (56) to define *t*\* implicitly by the equation

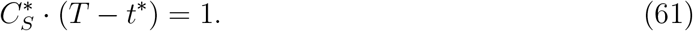

By solving for *t*\* in (61) we obtain

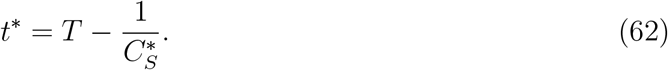

In doing so, note that we have extended this period of fruit-only growth from the interval [*T* – *ε, T*] to [*t*\*, *T*]. Note also that during the final interval we have by (28) that

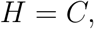

and because the Hamiltonian must be constant along the optimal trajectory, we have that

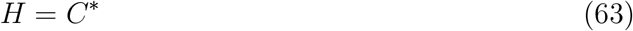

throughout the growing season.

### 4.2 Penultimate Interval - Vegetative/Reproductive Growth

We will now continue backwards to show that, prior to the final interval of fruit-only growth, we have a period of mixed vegetative/reproductive growth, during which both the shoots and fruits grow simultaneously. This interval will be referred to as the penultimate interval. Recall that at the beginning of the final interval we have that λ*_S_* = 1, and so (60) becomes

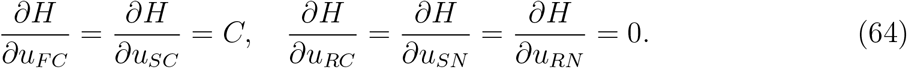

In order to apply (29) we need to determine the ordering of the partial derivatives of *H* in an interval immediately prior to *t*\*. We will accomplish this by a sequence of three lemmas, the proofs of which are included in Appendix B.1. We begin with the following inequality.

#### Lemma 3.

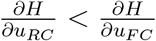 *in an open interval immediately prior to t*\*.

Note that, regardless of where 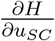 falls in the order of partial derivatives, the strict inequality in Lemma 3 means that there is no root growth during this period of time immediately prior to *t*\*. Next, we have the following, which indicates that allocation to shoots and to fruits at this point in the growing season are equally important to overall fitness.

#### Lemma 4.

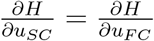 *in an open interval immediately prior to t*\*.

Putting together Lemmas 3 and 4, we get

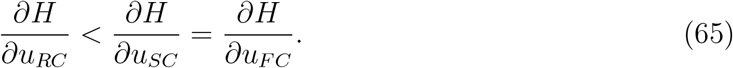

By (29), this means that *u_RC_* = 0 and by (42) and (43) we obtain

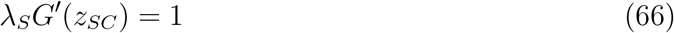

during this interval. Next, we state the following result which means that the shoots are growing during this stage. This will then be used to show that this interval is indeed marked by simultaneous fruit and shoot growth.

#### Lemma 5.

*u_SN_* = 1 *and u_SC_* > 0 *in an open interval immediately prior to t*\*.

At this point, note that we have only showed that

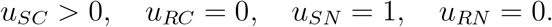

Lastly then, as λ*_S_* > 1 for this open interval before *t*\*, λ*_S_*(*t*\*) = 1, and λ*_S_* is continuous, we have that

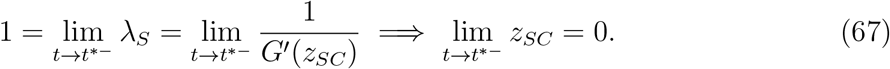

As we have already verified that *u_SN_* = 1 this means that

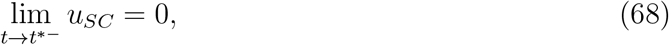

and so *u_SC_* is continuous at *t*\*. This also means that 0 < *u_SC_* < 1 during an open interval immediately prior to *t*\*, so here *u_FC_* > 0. Therefore, we have established the existence of a penultimate interval, during which

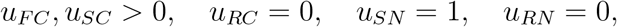

that is fruits and shoots alone grow simultaneously.

### 4.3 Marginal Values

Before discussing the stage which precedes the penultimate interval, we will briefly digress to discuss a key interpretation which will be relevant to the remaining portions of the growth path. One of the most elegant results from the work of Iwasa and Roughgarden (1984) was the relationship between the optimal growth trajectory and the marginal values of the sizes of the organs. In this context, the marginal value of the size of an organ at time *t* essentially measures the additional fruits present at time *T* given a unit increase in the size of that organ at time *t*. In their model, optimal growth was characterized as directing photosynthate to whichever organ had the highest marginal value, and only in the case that two organs had the same marginal value could resources be shared.

Iwasa and Roughgarden identified the adjoints in their model as the marginal values of the shoots and roots, and we can make a similar identification here. Let *J*(*S*_0_, *R*_0_, *t*_0_) be the maximum for the following optimal control problem.

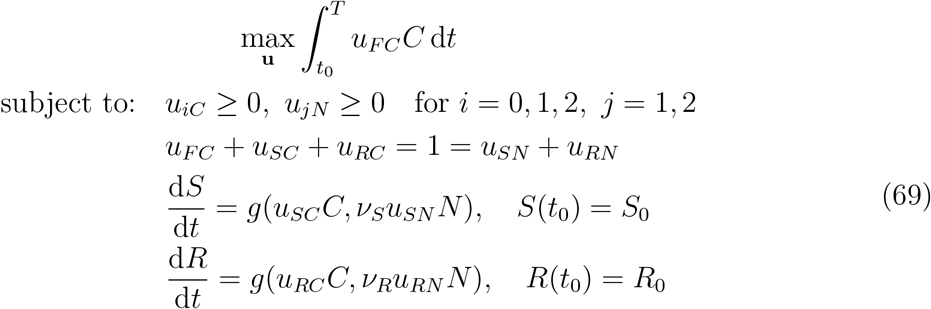

Note that this only differs from (27) in that the initial time is allowed to vary, and so if we initialize (69) with a point along the optimal trajectory for (27), the optimal trajectories for the two problems will coincide (see Lenhart and Workman 2007). In this context, given *t* ∈ [0, *T*] and *S*(*t*), *R*(*t*) optimal for (27), we have that

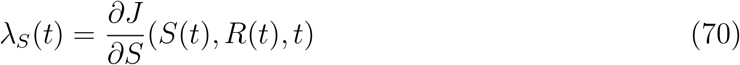

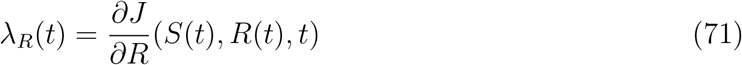

which means that λ*_S_*(*t*) and λ*_R_*(*t*) measure the increase in the objective function given a small instantaneous increase in either shoots or roots, respectively. See Kamien and Schwartz (1991) for a more general treatment. Therefore, as *J*(*S*(*t*), *R*(*t*), *t*) = *F*(*T*), we can identify λ*_S_*(*t*) and λ*_R_*(*t*) as the marginal values of the sizes of shoots and roots at time *t*, respectively. Furthermore, as resources invested in fruits are directly converted to tissue, we can identify 1 as the marginal value of the size of fruits.

In Iwasa and Roughgarden’s model (1984), the organ with the highest marginal value was the organ that received the full supply of photosynthate. However, when we consider the optimal trajectory laid out so far, we see that this is not the case in our model, which begs the question why. In particular, in Section 4.2 we see by (66) that λ*_S_G*′(*z_SC_*) = 1 throughout the interval. Because *z_SC_* ≠ 0 on the interior of this interval, we know that *G*′(*z_SC_*) < 1 and so λ*_S_*(*t*) > 1. Furthermore, as λ*_R_* = 0 at the end of the penultimate interval, we have a region where λ*_S_*(*t*) > 1 > λ*_R_*(*t*), that is the marginal value of the size of shoots is greater than either of the other marginal values, and yet both shoots and fruits are growing.

The reason for this apparent contradiction stems from the addition of the synthesizing units in our model. Consider again the end of the penultimate interval. Here we see that because the marginal value of shoots is highest, an instantaneous increase in shoots would be better for the plant than either an instantaneous increase in roots or fruits of the same amount. Note, however, that only the control functions can increase instantaneously, and it can be shown that during this phase the shoot SU alone cannot out-perform a mixed shoot/fruit strategy. We will make this more precise below.

Keeping our discussion limited to the penultimate interval, there are two thing in particular that we will examine to help us understand why resources are not necessarily directed solely to the organ with the highest marginal value, in this case shoots. First, we will give a biological interpretation of the necessary conditions during this interval, and second we will use a linear approximation to directly compare the trade off between a shoot-only strategy and a mixed fruit/shoot strategy during the penultimate interval.

To this end, recall that during the penultimate interval we have

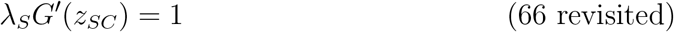

which relates the marginal values for shoots and fruits. To understand the biological relevance of this statement we need an interpretation of *G*′(*z_SC_*). By (20), we have that

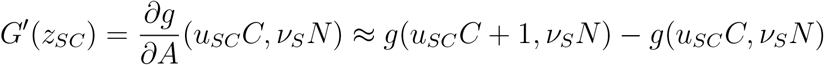

during the penultimate interval. That is, *G*′(*z_SC_*) is approximately the additional units of shoot present per unit time given a unit increase in the rate at which carbon is sent to the shoots. Multiplying this by the marginal value of shoots, λ*_S_*(*t*), approximates the additional fruits at time *T* given the additional shoots produced by a unit increase in carbon allocation to shoots. We can think of the marginal value of fruits being 1 in the same way, in that if we increase the fruit carbon flux, *u_FC_C*, by 1 we have an additional

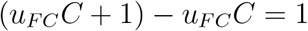

units of fruit per unit time, which yields an additional unit of fruit at time *T*. So, (66) essentially has the biological meaning that neither the shoot SU nor the fruit SU can outperform the other, and so a mixed strategy is best. In Iwasa and Roughgarden’s model (1984), raising the rate of carbon flux to an organ by one unit was approximately equivalent in the long run to increasing the biomass of that organ by one unit, which is why comparing marginal values was equivalent to using the necessary conditions.

### 4.4 Strategy Comparison

It is interesting to consider what happens when we attempt to construct a solution based on the marginal values alone during the penultimate interval. To this end, consider a point *t*^#^ in the penultimate interval such that λ*_S_* (*t*^#^) > 1 > λ*_R_* (*t*^#^). A growth strategy based solely on the marginal values would suggest shoot-only growth, whereas we know that a mixed fruit/shoot strategy is optimal here based on the necessary conditions. We will examine what happens if at time *t*^#^ there is a length-*ε* deviation from the optimal solution during which the mixed fruit/shoot strategy is replaced with a shoot-only strategy. To facilitate this analysis we have the following theorem.

#### Theorem 6.

*Let t*^#^ *be a point in the penultimate interval such that* λ*_S_*(*t*^#^) > 1 > λ*_R_*(*t*^#^) *and such that this ordering of the marginal values holds on* [*t*^#^, *t*^#^ + *ε*] *for some ε* > 0. *Let F*(*T*) *be the value obtained by following a mixed fruit/shoot strategy on* [*t*^#^, *t*^#^ + *ε*] *and then following the optimal trajectory on* (*t*^#^ + *ε*, *T*]. *Let F_alt_*(*T*) *be the value obtained by following a shoot-only strategy S_alt_*(*t*) *on* [*t*^#^, *t*^#^ + *ε*] *and then following the optimal trajectory on* (*t*^#^ + *ε*, *T*]. *Then, to leading order*,

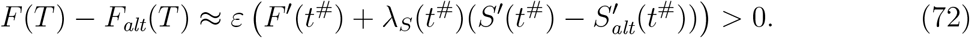

Note that this not only gives us an approximation for the trade off between the two strategies, but also means that to leading order the solution we have derived is better than the alternative solution suggested solely based on the marginal values. A full proof of Theorem 6 can be found in Appendix B.2, but here we will give an intuitive explanation for the approximation, as well as some numerical evidence.

We can use the marginal values for shoots and fruits at time *t*^#^ to approximate the impact each strategy on the interval [*t*^#^, *t*^#^ + *ε*] will ultimately have on the value of *F*(*T*). First, we will consider a mixed fruit/shoot strategy on [*t*^#^, *t*^#^ + *ε*] then we have

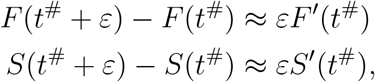

and so the total contribution to *F*(*T*) during the interval [*t*^#^, *t*^#^ + *ε*] is approximately

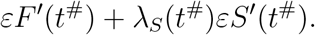

Likewise, if we consider a shoot-only strategy on [*t*^#^, *t*^#^ + *ε*] then we have

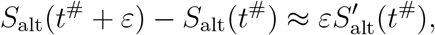

and so the total contribution to *F*_alt_(*T*) here is approximately 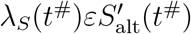. Therefore, assuming that *ε* is small enough that any difference between the optimal growth paths that the two solutions follow after *t* = *t*^#^ + *ε* is negligible we conclude that

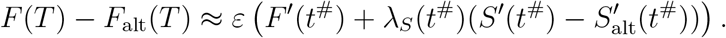

We use a numerical experiment to illustrate (72). Using methods we will discuss in Section 6, we directly compared the difference *F*(*T*) – *F*_alt_(*T*) with the approximation 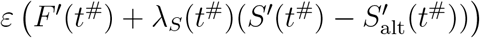. Here we used the terminal conditions *S*(*T*) = 222.11 and *R*(*T*) = 111.55 (see Section 7.2). Using ten linearly spaced points in the penultimate interval where λ*_S_* > 1 > λ*_R_*, and starting at each point we simulated shoot-only growth on a length-*ε* interval, with *ε* = 10^−3^, and then continued along the trajectory that is optimal for the new states at the end of each interval. This allowed us to compute *F*(*T*) – *F*_alt_(*T*) directly. We also computed the estimate 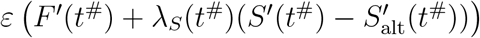 for each of the ten sampled points, and then compared the computed value to the estimate. The results of this experiment are shown in Figure 2. Observe that the estimate follows the computed values, which illustrates the accuracy of our estimate. Note also that that *F*(*T*) – *F*_alt_(*T*) > 0, meaning that the mixed fruit/shoot strategy out-performs the shoot-only strategy, as claimed.

**Figure 2:**
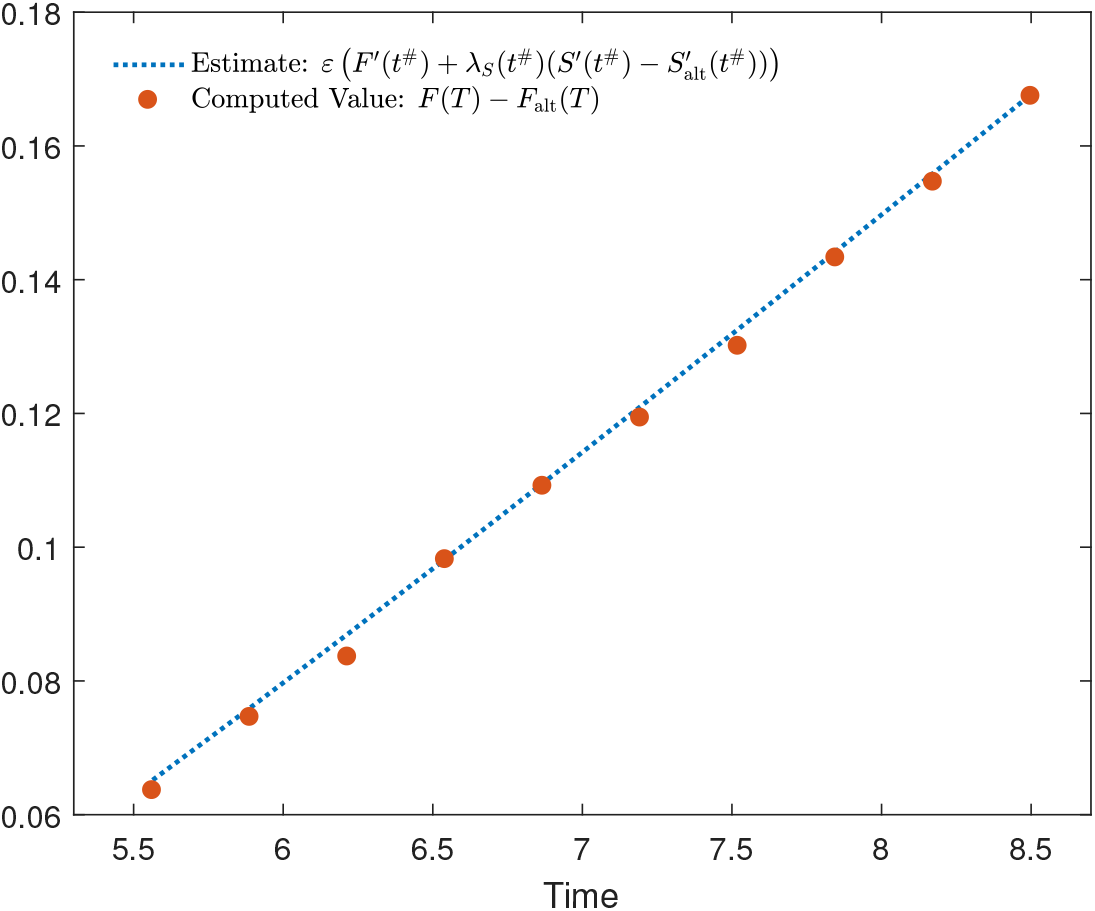
Numerical verification of (72) for ten times during the penultimate interval

### 4.5 Balanced Growth - Mixed Vegetative Growth

At this point, we have established the existence of a final interval during which only the fruits are growing, and a period before this when both the shoots and fruits are growing together. Continuing backwards, we have arrived at a juncture where there are seemingly several options. This is made complicated by the fact that *z_RC_* is not defined during the penultimate interval, and so it is not clear directly from the necessary conditions (29) which growth pattern is optimal here.

Prior to this period of mixed shoot/fruit growth, there are several possibilities. These are: fruit growth, root/fruit growth, root/shoot/fruit growth, root/shoot growth, shoot-only growth, and root-only growth. Because we assume that fruits are only carbon-dependent, it is never optimal for the plant to invest fully in fruits unless the shoots are done growing. We can also rule out either shoot-only growth or root-only growth here. As we will subsequently see, the first stage of growth consists of either shoot-only growth or root-only growth, and while it may be the case that the length of this second phase is infinitesimally small, we still consider it as distinct from the first phase. That said, it is not optimal to alternate between growing a only shoots and only roots, because after addressing any initial imbalance between shoots and roots, to do so would only create further resource limitation for the growing organ.

This leaves us with three remaining options: root/fruit growth, root/shoot/fruit growth, or root/shoot growth. We will use an argument based on our discussion of marginal values in Section 4.3 to eliminate both root/fruit and root/shoot/fruit growth. The necessary conditions (29) provide conditions that must be met on an open interval in each case:

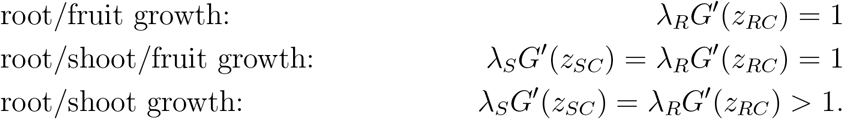

As with λ*_S_G*′(*z_SC_*) in Section 4.3, we can think of λ*_R_G*′(*z_RC_*) as approximating the additional units of fruit at time *T* given the additional roots produced by a unit increase in carbon allocation to roots. When the growth path follows λ*_R_G*′(*z_RC_*) = 1, this means that, with respect to carbon, the root SU can do no better for the plant in the long run than the fruit SU can do at present. In the same vein, when a growth path follows λ*_R_G*′(*z_RC_*) > 1, the long-term returns of investing carbon in roots exceeds the immediate returns of investing carbon in fruits. This means that root/shoot growth is optimal here, which is consistent with the work of Iwasa and Roughgarden (1984).

### 4.6 Initial Phase - Shoot or Root Growth

It is not optimal to have an interval without fruit growth between two intervals which include fruit growth. Such a growth path could be improved by swapping the first two intervals so that the fruits grow continuously. This means that there is no fruit growth prior to the balanced growth phase. As we have already discussed the idea that alternating phases of shoot-only and root-only growth is not optimal, we are left with one option for the initial phase, namely that it consists of either shoot-only growth or root-only growth. Here the plant addresses deficiencies present in either vegetative organ depending on the initial conditions. Note that this is also the first stage that we see in the single-resource case described in Iwasa and Roughgarden (1984).

## 5 Phase Dynamics and Transitions

In this section, we will present the basic equations governing the dynamics in each phase of the solution to (27) as well as show that *z_SC_* and *z_RC_* are continuous between any two consecutive phases in which they are defined. To keep this section organized, we will go through the dynamics for each phase in chronological order and then go on to discuss the transitions between phases. We will also limit this section to include only phase-specific versions of the differential equations for states and adjoints, relevant algebraic constraints due to the necessary conditions, and any additional equations used for understanding the transitions between phases. When we discuss the numerical scheme, we will make use of several additional equations which govern dynamics in various stages but are not relevant here. In what follows, we will also make frequent use of the notation *G*_2_(*x*) = *G*′(1/*x*).

### 5.1 Initial Phase: Shoot-Only Growth

During shoot-only growth we have

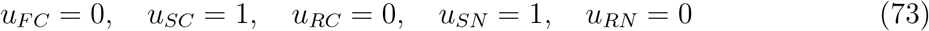

and only *S*, λ*_S_*, and λ*_R_* are changing, with differential equations in time given by

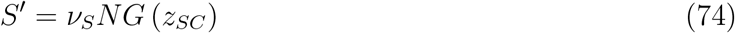

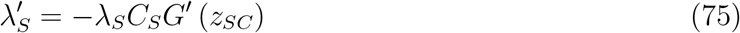

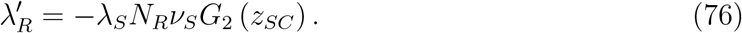

Here we have that

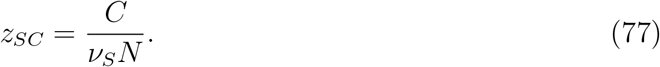

Note also that in this stage both *N* and *N_R_* are constant because *R* is constant. Furthermore, because we know by (63) that *H* = *C*\*, the Hamiltonian (35) during this phase becomes

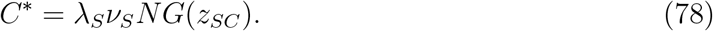

By solving for λ*_S_*, and employing (24), we obtain two expressions for λ*_S_* during this phase:

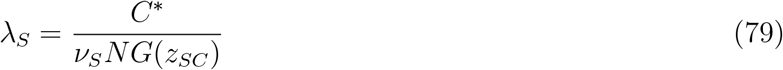

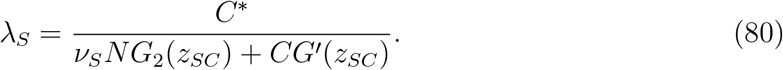

### 5.2 Initial Phase: Root-Only Growth

During root-only growth we have

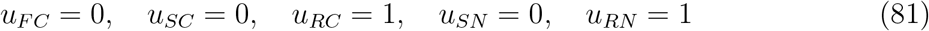

and only *R*, λ*_S_*, and λ*_R_* are changing, with differential equations in time given by

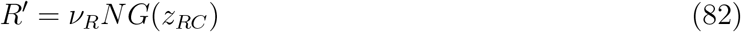

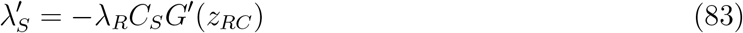

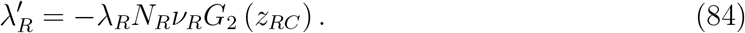

Here we have that

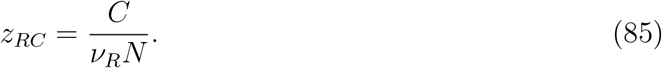

Note also that in this stage both *C* and *C_S_* are constant because *S* is constant. Furthermore, because we know by (63) that *H* = *C*\*, the Hamiltonian (35) during this phase becomes

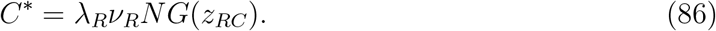

By solving for λ*_R_*, and employing (24), we obtain two expressions for λ*_R_* during this phase:

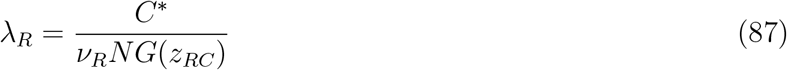

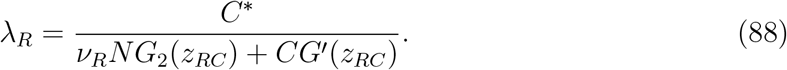

### 5.3 Balanced Growth - Shoot/Root Growth

During balanced growth we have

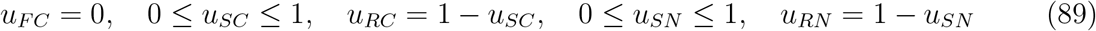

and so by (29) we have that the following two conditions must hold during this phase

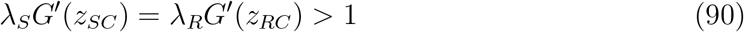

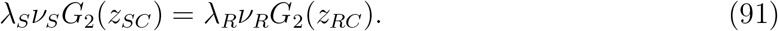

During this stage *S*, *R*, λ*_S_*, and λ*_R_* are changing. The differential equations in time for these four during this phase are given by

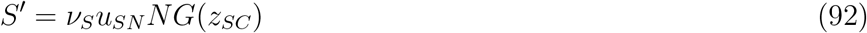

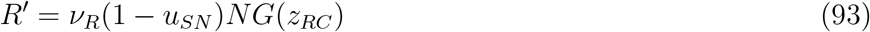

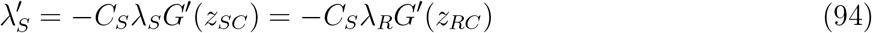

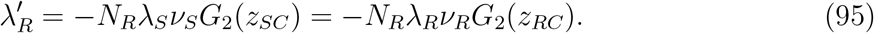

Furthermore, again taking advantage of the fact that (63) gives us *H* = *C*\*, we can rewrite the Hamiltonian (35) during this phase as

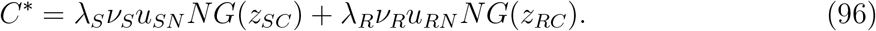

Rewriting this using (24) we get

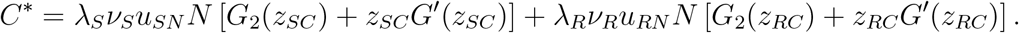

Simplifying, and using (90) and (91) to write everything in terms of *z_SC_*, gives us

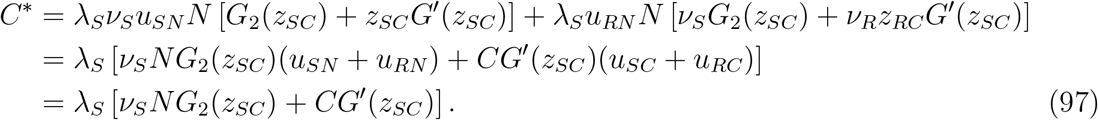

Solving for λ*_S_* gives us

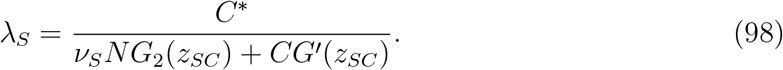

Using (90) and (91) to rewrite (97) in terms of *z_RC_* instead leads to

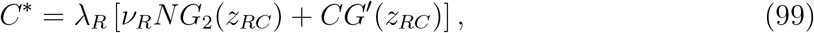

which upon solving for λ*_R_* gives us

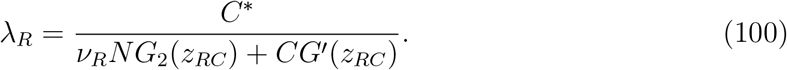

Note that (98) and (100) are consistent with (80) during shoot-only growth, and (88) during root-only growth, respectively. While we do not assume that either *z_SC_* or *z_RC_* is continuous between phases (because the controls need not be), we will use this consistency later to show continuity between the initial and balanced growth phases.

### 5.4 Penultimate Interval - Shoot/Fruit Growth

During the penultimate interval, we have

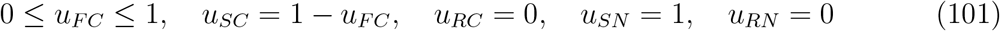

and so by (29) we have that

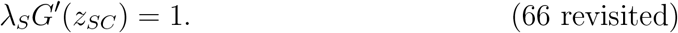

Note that here we have

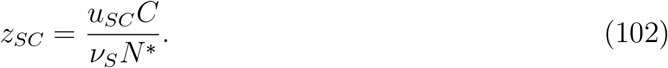

During this interval *S*, *F*, λ*_S_*, and λ*_R_* are all changing. The differential equations in time for these four during this stage are

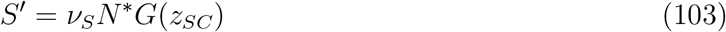

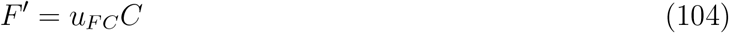

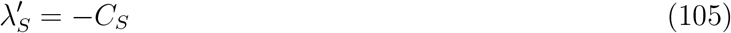

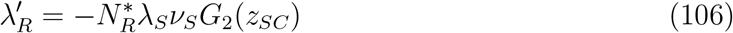

Furthermore, again taking advantage of the fact that (63) gives us *H* = *C*\*, we can rewrite the Hamiltonian (35) during this phase as

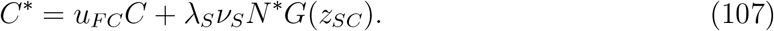

Using (24) and (102) we get

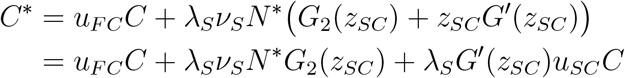

which by (66) becomes

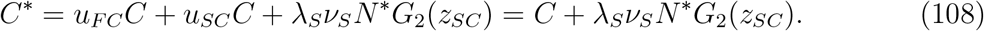

Now, at this point we can either solve for λ*_S_* and obtain

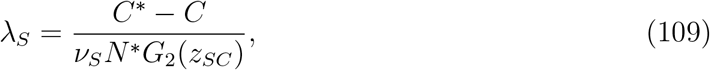

or we can use (66) to rewrite (108) as

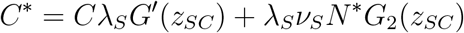

and solve to obtain the familiar expression

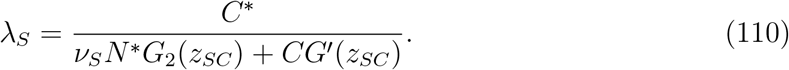

As we commented earlier, the reappearance of this expression for λ*_S_* suggests that *z_SC_* may be continuous between the balanced growth phase and penultimate interval. We will prove this later with the help of (109).

Another important result in the penultimate interval which we prove here is that *z_SC_* is strictly decreasing throughout this phase. This will be especially important because, as we will see in Section 6.3, it will be advantageous for us to think of *z_SC_*, rather than *t*, as the integration variable for the numerical scheme during the penultimate interval. In particular, we will see that numerically solving the differential equations for this phase in *t* requires integrating a singularity, whereas solving the differential equations in *z_SC_* does not.

#### Lemma 7.

*z_SC_ is strictly decreasing during the penultimate interval*.

*Proof*. Recall that during the penultimate interval we have the relationship

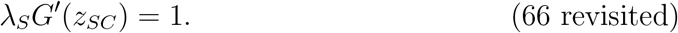

Differentiating both sides with respect to *t* yields

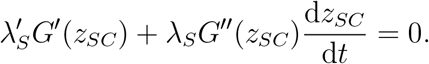

Using the fact that 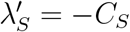, we have

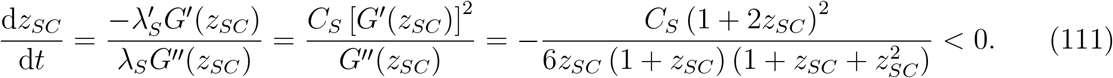

Therefore, *z_SC_* is strictly decreasing throughout the penultimate interval.

### 5.5 Final Interval - Fruit-Only Growth

During the final interval of fruit-only growth we have

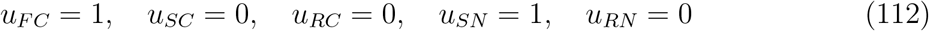

and only *F* and λ*_S_* are changing, with differential equations in time given by

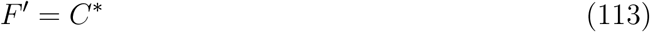

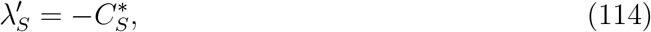

Because λ*_S_* (*T*) = 0 we also have

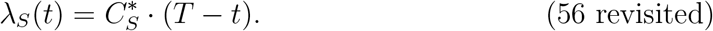

Lastly, recall that λ*_R_*(*T*) = 0 and 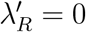 during this phase, and so λ*_R_* = 0 here as well.

### 5.6 Initial Phase to Balanced Growth Transition

Now that we have established the main dynamics in each phase, we will turn our attention to the transitions between phases. Again, we will proceed in chronological order, beginning with the transition from the initial phase to balanced growth. We can use the fact that the states and adjoints are continuous at this boundary to characterize this transition. In particular, we will show that *z_SC_* is continuous at this transition when the first stage consists of shoot-only growth and *z_RC_* is continuous at this transition when the first stage consists of root-only growth. In the first case this will mean that the ratio of the shoot carbon flux to shoot nitrogen flux is continuous across this boundary, and in the second case the ratio of root carbon flux to root nitrogen flux is continuous.

Due to the symmetry between equations (98) and (80) and equations (100) and (88) we will streamline the argument by considering a ‘generalized’ version of these equations:

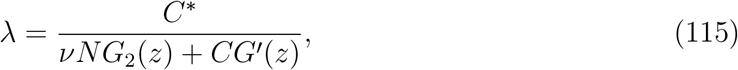

where (λ, *ν*, *z*) is either (λ*_S_*, *ν_S_*, *z_SC_*) or (λ*_R_*, *ν_R_*, *z_RC_*) depending on whether the initial stage is shoot-only growth or root-only growth, respectively. We call this transition point 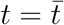, let 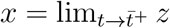, and note that 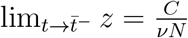. Now, taking limits of (115) from both sides:

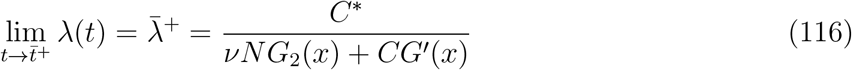

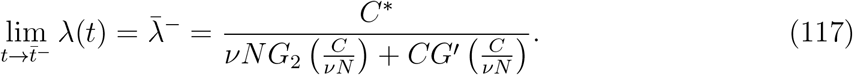

Note that while we don’t know the values of *C* and *N* at 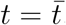, in what follows it will only matter that they are the same in both (116) and (117). At 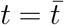 we have 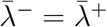 and so

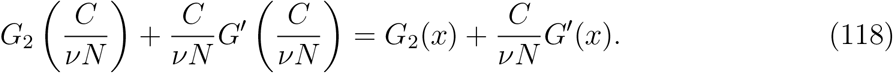

Let

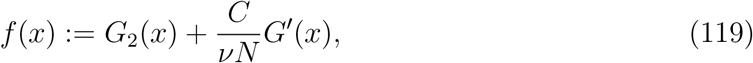

and note that is clear from (118) that 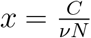 is a solution to

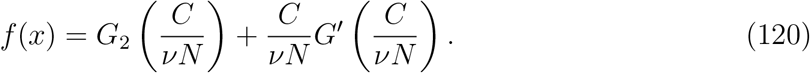

Here we will show that this solution is unique. Consider *f*′(*x*), recalling (25):

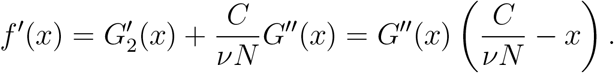

Note that because *G*″(*x*) ≤ 0 for *x* ≥ 0, we have that

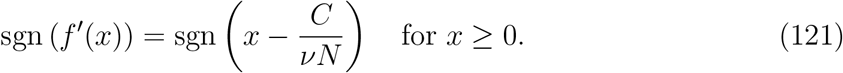

Now, as *f* is continuous and, by (121), we have that *f* is decreasing on 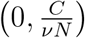 and increasing on 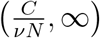, it must be the case that 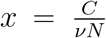 is the only solution to (120). Therefore, we have that *z* is continuous at 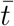 which verifies our claim that *z_SC_* is continuous at the transition from shoot-only growth to balanced growth and *z_RC_* is continuous at the transition from root-only growth to balanced growth.

### 5.7 Balanced Growth to Penultimate Interval Transition

As in Section 5.6, we can use the fact that the states and adjoints are continuous to show that *z_SC_* is continuous at the boundary between the balanced growth and penultimate intervals. Let 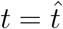 denote this transition point, and let 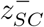 and 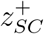 be the left and right limits of *z_SC_* at 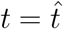, respectively. We have, then, the following theorem.

#### Theorem 8.

*z_SC_ is continuous at* 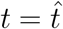.

We include the details of the proof in Appendix C and only give a sketch of the proof here. To that end, note that by combining (98), (109), and (66), and writing 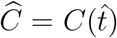, we have that 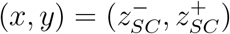 must solve

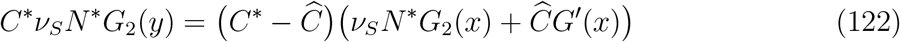

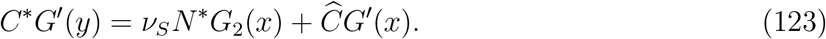

Because *G*_2_ and *G*′ are both one-to-one, we have that *y* is a function of *x* in (122) and (123).

Figure 3 illustrates the three types of solutions to (122) and (123), depending on the values of the parameters. Note that the intersection points (*x*, *y*) of the two curves correspond to possible pairs 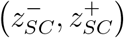. In particular, there are two common features of each plot:

1. In each plot there is one *y* value that solves (122) and (123); this determines 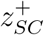.
2. There are at most two intersection points, one is always at 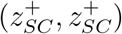.

**Figure 3:**
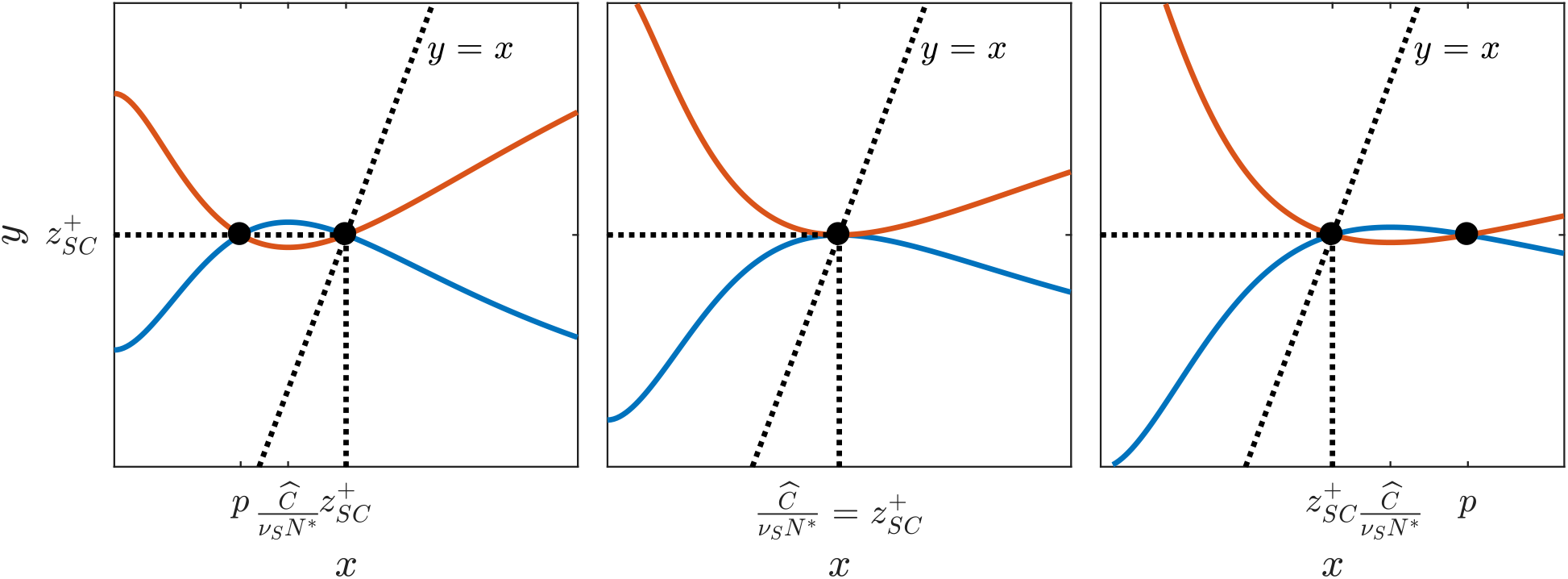
Three cases for graphs of (122) and (123). Case 1 (left): 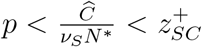 Case 2 (center): 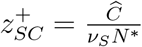. Case 3 (right): 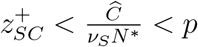

For each case in Figure 3, the only admissible solution is 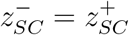. In Case 2 (Figure 3, center) this is the only possibility. In Case 1 (Figure 3, left) we observe that

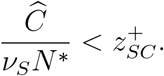

However, as we know that

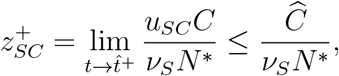

this would mean that

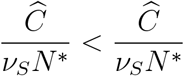

which is impossible. Therefore, Case 1 is not an admissible configuration of solutions to (122) and (123). In Case 3 (Figure 3, right), there is a possible intersection point at 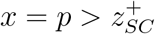. However, if 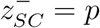, then as *G*′ is strictly decreasing we have that 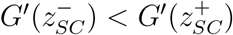, and

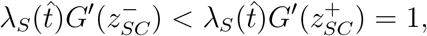

which is impossible by (90). Therefore, the only possible solution to (122) and (123) is 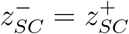, which means that *z_SC_* is continuous at 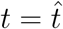. It remains to show that these three cases actually characterize the solution, the details of which are included in Appendix C.

### 5.8 Penultimate Interval to Final Interval Transition

Note that in the final interval neither *z_SC_* nor *z_RC_* is defined, so we won’t be addressing continuity of either one here. We will, however, analyze the controls at this transition and conclude that at this particular junction we actually have continuity of the controls.

First, recall that in Section 4.2, we showed that 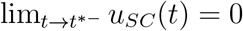. Because during the penultimate interval *u_FC_* = 1 – *u_SC_* we have that 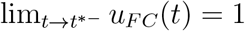 as well. Note also that during the penultimate interval we have from (101) that *u_RC_* = 0, *u_SN_* = 1, and *u_RN_* = 0, and by (112) we have that all the controls maintain their values in the final interval as well. Therefore, all the controls are continuous at *t* = *t*\*.

Next, we will prove the following lemma which shows that the transition to fruit-only growth occurs rapidly at the end of the penultimate interval.

#### Lemma 9.

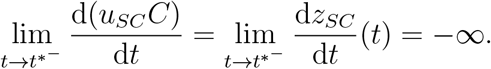

*Proof*. The first equality follows from the penultimate interval definition of *z_SC_* (102). For the second equality, recall that by (67), *z_SC_* → 0^+^ as 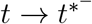. Lastly, by (111), we have

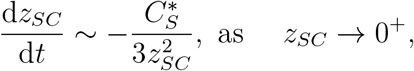

which because 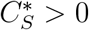 means that

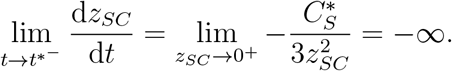

## 6 Numerical Scheme

In this section we will outline the numerical scheme we developed for solving the optimal control problem (27). Because the solution exhibits a multi-phase structure, standard numerical methods such as forward-backward sweep or shooting (see Lenhart and Workman 2007) aren’t well-suited. Our approach is to construct a numerical scheme for solving the problem backward in time. There are two primary components to the numerical scheme. First, we obtain the map

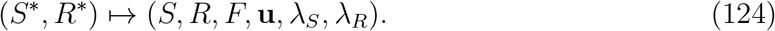

Second, we use MATLAB’s built-in nonlinear equation solver `fsolve` to find the map

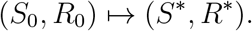

Ultimately, then, we obtain the map

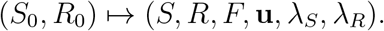

We will now direct out attention to the numerical scheme for finding (124), proceeding phase-by-phase in reverse order. During each phase, we use the fourth-order Runge-Kutta method (RK4) to solve differential equations which govern the phase dynamics, and use algebraic equations to update the controls, thus allowing us to avoid having to update the controls iteratively. We will use the fact that the Hamiltonian is constant along the optimal trajectory, as well as the information we have about transitions from Section 5, to find the boundaries between phases and ultimately stitch the four phases together to form one complete solution. We will discuss how the numerical scheme works without getting into the fine numerical details. The actual MATLAB code is included in Appendix E.

### 6.1 Penultimate Interval

We begin by using equation (62) to find the end of the penultimate interval (*t*\*). Now, as the dynamics during the penultimate interval depend on *z_SC_*, we would ideally use (111) and (103) to solve for *z_SC_* and *S* simultaneously, and use (102) to update the controls. However, recall that by Lemma 9, *z_SC_* approaches a vertical tangent as 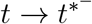, which makes it difficult to solve for *z_SC_* numerically. Note, however, that by the same reasoning we have that 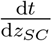 approaches 0 as *z_SC_* → 0^+^. So, to remedy this numerical difficulty, we take advantage of this fact, along with the fact that, by Lemma 7, *z_SC_* is strictly decreasing throughout the penultimate interval, to think of *z_SC_* as our ‘time’ and derive differential equations for *t*, *S*, and λ*_R_* in terms of *z_SC_*. This, combined with the fact that during the penultimate interval *R* is constant, λ*_S_* is algebraically related to *z_SC_* via (66), and *F* depends on the value of 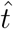, gives us everything we need to find the solution during this phase numerically. Note that because *z_SC_* is decreasing, solving forward in *z_SC_* is equivalent to solving backward in time.

To this end, note that by (111) we have that

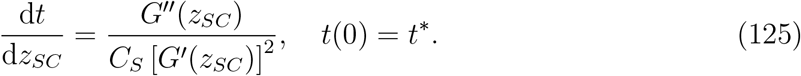

Furthermore, using (103), (106), and (125) we obtain

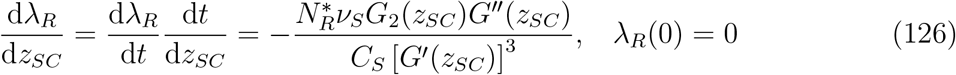

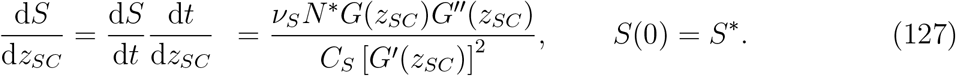

We use RK4 to solve (125), (126), and (127) forward in *z_SC_*, and use (102) to update *u_SC_*. Also, recall that λ*_S_* = 1/*G*′(*z_SC_*), *u_FC_* = 1 – *u_SC_*, and the other three controls are constant by (101). Upon reordering by *t*, this gives us all of the states, adjoints, and controls during the penultimate interval, with the exception of fruits, which we will determine after identifying the correct transition point from balanced growth to the penultimate interval 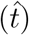.

We stop solving forward in *z_SC_* when *u_SC_* becomes unbounded, as this gives us an upper bound on the value of *z_SC_* (lower bound for the value of *t*) for which the transition from balanced growth to the penultimate interval can occur. Furthermore, because during the balanced growth phase we have by (90) that λ*_R_G*′(*z_RC_*) > 1, it must be the case that 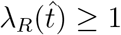 because *G*′(*z*) ≤ 1. Finding where λ*_R_* =1 gives us a lower bound on the value of *z_SC_* (upper bound for the value of *t*) for which the transition from balanced growth to the penultimate interval can occur. This gives us an interval 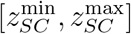 in which to search for 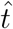.

### 6.2 Locating the Start of the Penultimate Interval

This portion of the numerical scheme begins with the interval 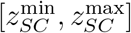 identified in Section 6.1. Since we know that the Hamiltonian must be constant along the optimal trajectory, we use the balanced-growth specific formulation of *H* in terms of λ*_R_* and *z_RC_*, given by (99):

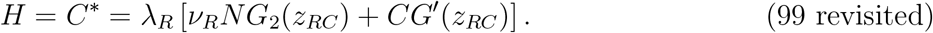

Starting with 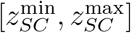, we use a binary search to find the smallest interval, relative to current RK4 step size, which contains the point 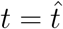 at which (99) is satisfied. Next, we use a smaller step size, and follow the procedure outlined in Section 6.1 to increase the resolution of the solution and repeat the binary search until we find 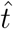 to within some specified tolerance. This also gives us the values of the states, adjoints, and controls at 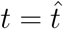.

One crucial feature on the process that we have not mentioned so far is the fact that since *z_RC_* is not defined in the penultimate interval, we need to compute its limit from balanced growth at every step in the binary search. This is made possible by the fact that *z_SC_* is continuous at 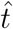, as established in Section 5.7. Therefore, we can use (90) and (91) to obtain

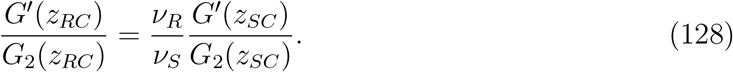

Using MATLAB’s built-in nonlinear equation solver `fsolve`, we use the value of *z_SC_* at a particular point to calculate what *z_RC_* would be if balanced growth ended at that point. This value of *z_RC_* is then used to evaluate the right-hand side of (99) in the binary search.

### 6.3 Fruits - Penultimate Interval

In Section 6.1 we were unable to solve for *F* because we didn’t know the value of 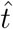. This was resolved in Section 6.2, so at this point we use RK4 to solve (104) forward in time during the penultimate interval. This completes the numerical solution for the penultimate interval.

### 6.4 Final Interval

Before continuing backward in time we take advantage of the fact that, by the process described in Section 6.3, we now know the value of *F* at the beginning of the final interval. Here the controls are constant by (112), *S*, *R*, and λ*_R_* are constant, and λ*_S_* is given by (56). Lastly, using the value of *F*(*t*\*) obtained as described in Section 6.3, and the fact that *F*’ is constant during the final interval by (113), we have that

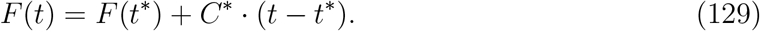

### 6.5 Balanced Growth

Upon locating 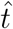 as discussed in Section 6.2, we have the value of the states, adjoints, *z_SC_*, and *z_RC_* at the end of the balanced growth stage. Recall that the controls need not be continuous, so we do not immediately know their values at the end of balanced growth. We can, however, start with the equations for *z_SC_* and *z_RC_* during balanced growth

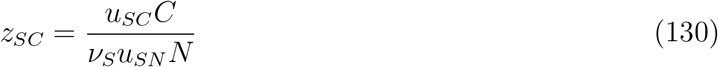

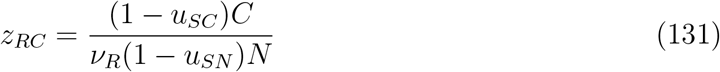

and solve for *u_SN_* and *u_SC_*:

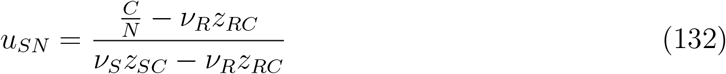

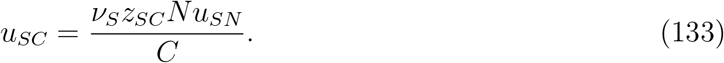

We derive differential equations for *z_SC_* and *z_RC_* during this phase, and ultimately use (132) and (133) to obtain the controls. The differential equations in time for *z_SC_* and *z_RC_* are given below, and the derivation is included in Appendix D.

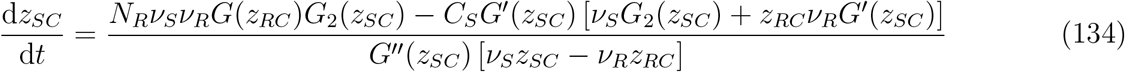

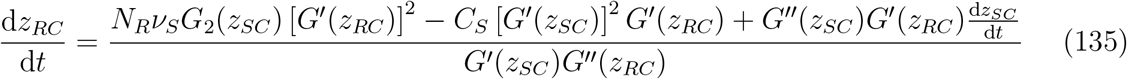

Beginning at 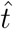, we use RK4 to simultaneously solve (92), (93), (94), (95), (134), and (135) backward in time to obtain *S*, *R*, λ*_S_*, λ*_R_*, *z_SC_*, and *z_RC_*, respectively. We use (133) and (132) to eliminate *u_SC_* and *u_SN_* in (92) and (93) so that these six differential equations are expressed exclusively in terms of these six variables. We then use (133) and (132) to obtain *u_SC_* and *u_SN_*, as well as the rest of the controls via (89). We continue until one of the controls exits [0,1], which gives us the earliest time for the beginning of the balanced growth phase.

### 6.6 Locating the Start of Balanced Growth

The next piece of the numerical scheme finds the start of balanced growth, 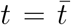. Recall that in Section 5.6, we showed that either *z_SC_* or *z_RC_* is continuous at 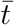, depending on which is defined in the initial phase. In either case, the transition must occur at a point where

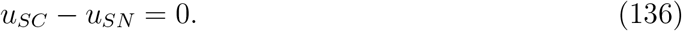

This is because in the case where the initial phase consists of shoot growth we have

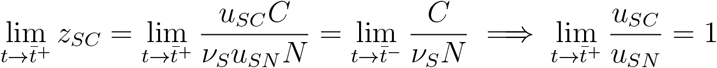

and in the case where the initial phase consists of root growth we have

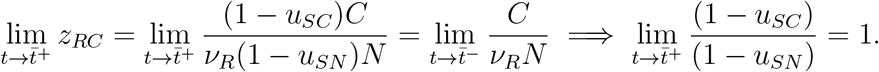

We use essentially the same procedure as in Section 6.2 for refining the transition point between the balanced growth and the penultimate intervals, with the exception that finding 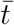 doesn’t require computing the limits of any quantities from the earlier phase, as we had to do with *z_RC_* at 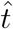. We use a binary search to locate the smallest interval about which (136) is satisfied, and then use RK4 to solve equations (92), (93), (94), (95), (134), and (135) backward in time on a smaller integration mesh. We repeat this process until we have found a point at which (136) is met to within some specified tolerance. We call this point 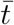.

Note that if no such point exists then the plant begins in balanced growth, in which case there are only three phases. If 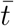 exists, the next step in the numerical scheme is to determine the makeup of the initial phase. Using (79) and (87), we compute the following at 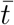:

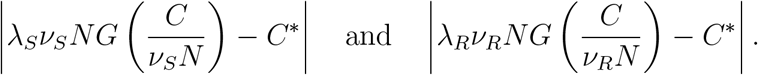

If the first is smaller then the initial phase consists of shoot growth, and if the second is smaller the initial phase consists of root growth.

### 6.7 Initial Phase

#### 6.7.1 Shoot-Only Growth

If the first phase consists of shoot-only growth, we use RK4 to solve (74), (75), and (76) simultaneously backward in time until we reach *t* = 0. This gives us *S*, λ*_S_*, and λ*_R_*, respectively. The controls are constant here and given by (73), and 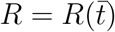.

#### 6.7.2 Root-Only Growth

If the first phase consists of root-only growth, we use RK4 to solve (82), (83), and (84) simultaneously backward in time until we reach *t* = 0. This gives us *R*, λ*_S_*, and λ*_R_*, respectively. The controls are constant here and given by (81), and 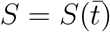.

## 7 Numerical Results

In this section, we will present some of the primary results from the numerical simulations of the model. In particular, we will look at several ‘representative’ simulation results that showcase the different strategies a plant can employ to maximize fruit production, as well as help us understand the relationship between initial and terminal conditions. In each simulation, we made the simplifying assumptions that *C*(*S*) = *S* and *N*(*R*) = *R*, which ignore any possibility of self-shading as the plant grows. We chose *ν_R_* = 1 for convenience, and chose *ν_S_* = 1/3, which had the effect of exaggerating the lengths of the various phases, resulting in easier-to-interpret plots. Note that this results in 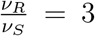, which is likely a bit higher than data suggests (see Hicks 1928; Minden and Kleyer 2014). Lastly, we chose *T* =10 because in the case that *C*(*S*) = *S*, we have by (62) that *t*\* = 9, which again facilitated our interpretation of the numerical results. We will begin with four examples of different strategies which all reach the same optimal value of fruits at time *T*.

### 7.1 Initial Shoot Growth

Here we chose terminal conditions *S*\* = 223.20 and *R*\* = 112.86. This results in *F*(*T*) = 900, *S*_0_ = 17, and *R*_0_ = 84. In this case we see the full four-stage structure of the solution. There is an initial phase of shoot growth, followed by a period of balanced growth between shoots and roots, a penultimate interval of shoot and fruit growth, and finally a period of fruit-only growth at the end of the growing season. The states and controls are shown in Figure 4.

**Figure 4:**
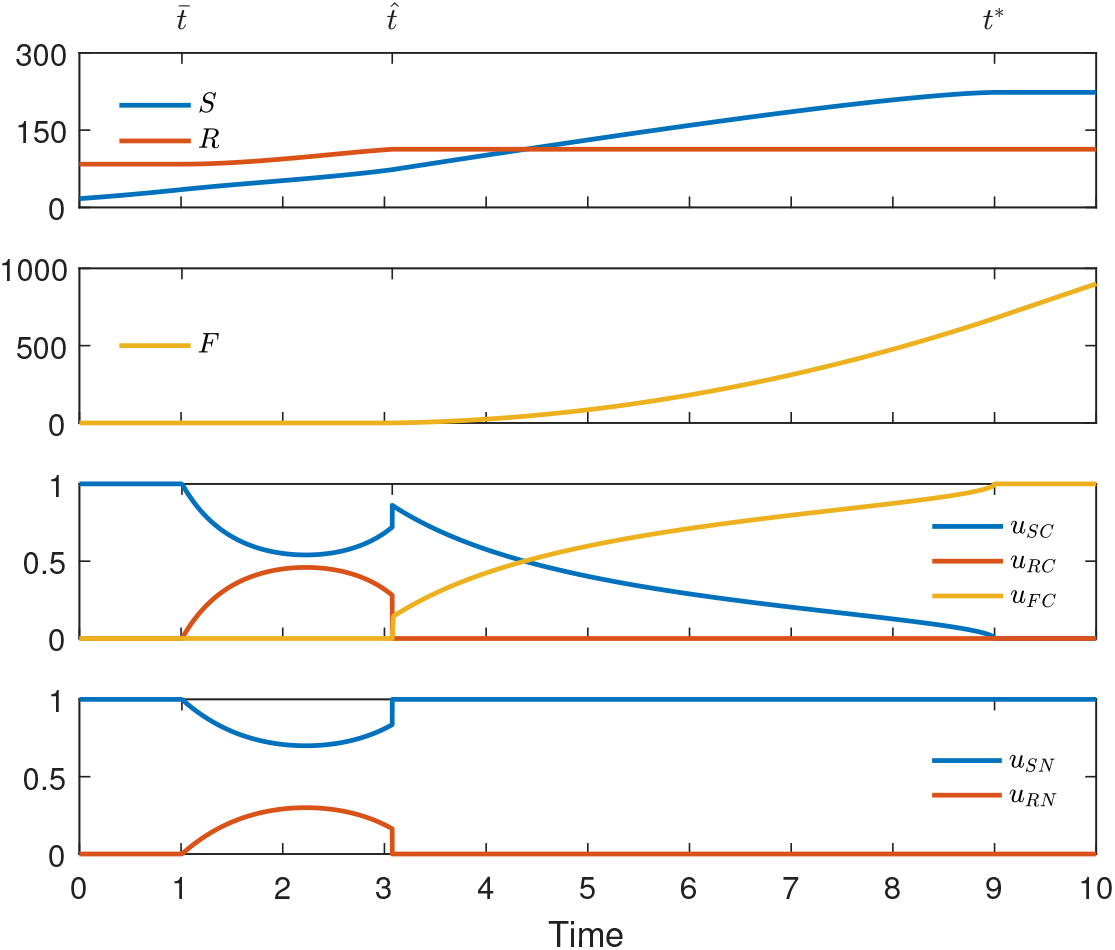
Initial shoot growth. From top to bottom: Shoots and roots, fruits, carbon controls, nitrogen controls. Shoots, roots, and fruits in units of carbon and controls dimensionless

The lower two plots in Figure 4 show the carbon and nitrogen allocation strategies for this plant. Following the initial period of shoot growth, there is a short period during which *u_RC_* and *u_RN_* are increasing, signifying a period of increasing root production. Shortly after *t* = 2, we see *u_RC_* and *u_RN_* begin to decrease, signifying that although it is still advantageous to invest resources in roots, the plant ultimately needs to prioritize shoot production again to prepare for mixed shoot/fruit growth. During the penultimate interval, we see the plant gradually stop investing in shoots, before switching to fruit-only growth at time *t* = 9.

### 7.2 Initial Root Growth

Here we chose terminal conditions *S*\* = 222.11 and *R*\* = 111.55. This results in *F*(*T*) = 900, *S*_0_ = 48.9, and *R*_0_ = 28.9. In this simulation we again see all four stages, however here we see an initial phase of root growth instead of the initial phase of shoot growth we saw previously. The initial conditions of the two simulations are quite different, but the terminal conditions are nearly the same. This, as we will discuss later, suggests that optimal growth tends to smooth out initial transients. The states and controls are shown in Figure 5. The two graphs at the bottom of Figure 5 show a different allocation strategy than appeared in the case of initial shoot growth in Figure 4. In particular, following the initial period of root growth, we see a decline in both carbon and nitrogen allocation to the roots throughout the entire balanced growth phase, and the steady increase in allocation to shoots.

**Figure 5:**
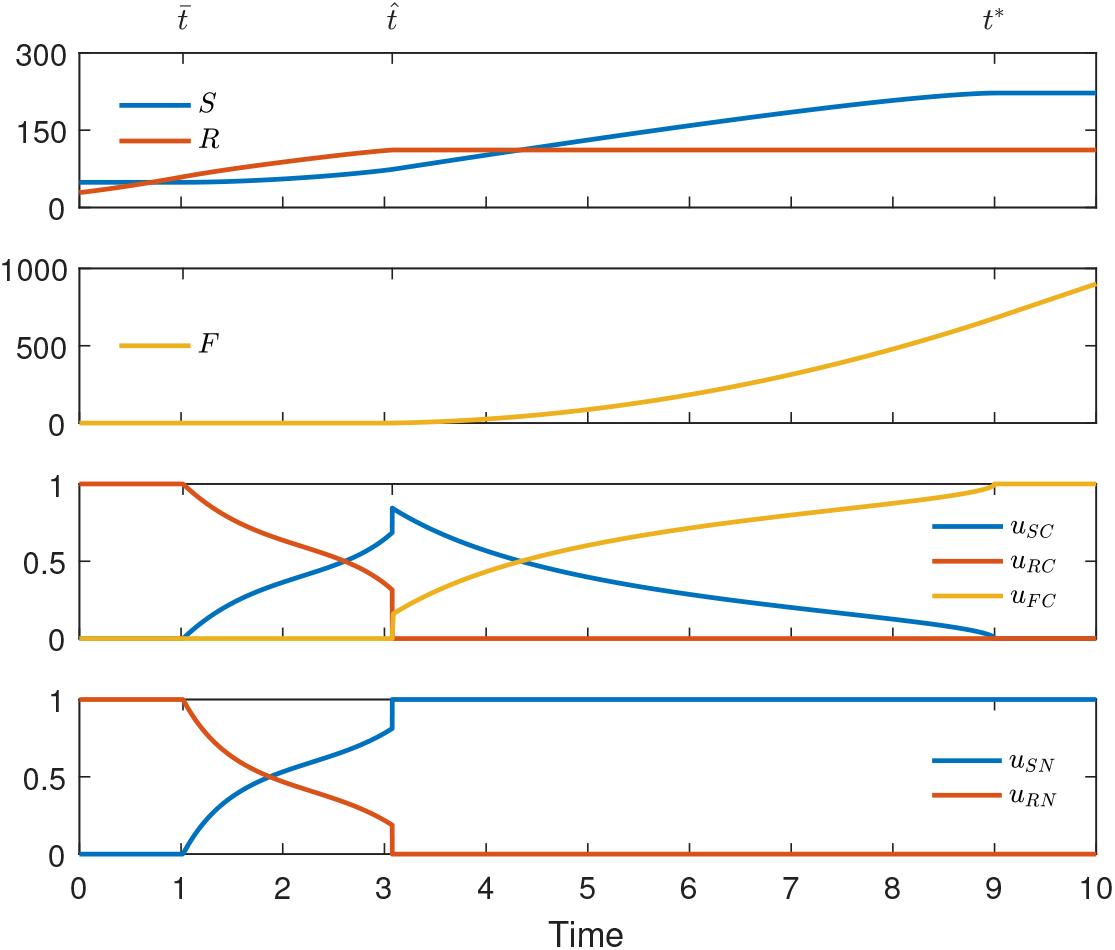
Initial root growth. From top to bottom: Shoots and roots, fruits, carbon controls, nitrogen controls. Shoots, roots, and fruits in units of carbon and controls dimensionless

### 7.3 Balanced Growth First - Type S

Here we chose terminal conditions *S*\* = 222.71 and *R*\* = 112.27. This results in *F*(*T*) = 900, *S*_0_ = 30.3, and *R*_0_ = 59.74. The states and controls are shown in Figure 6. In this simulation, we see a plant which starts in balance, and therefore forgoes the initial phase of shoot-only or root-only growth. That said, comparing Figures 6 and 4, we see a similar allocation strategy during balanced growth. In particular, both show that allocation to shoots initially decreases before increasing again. This behavior corresponds to initial conditions that are more biased toward shoot deficiency than root deficiency. For this reason, we refer to this type of initially balanced growth as ‘Type S.’

**Figure 6:**
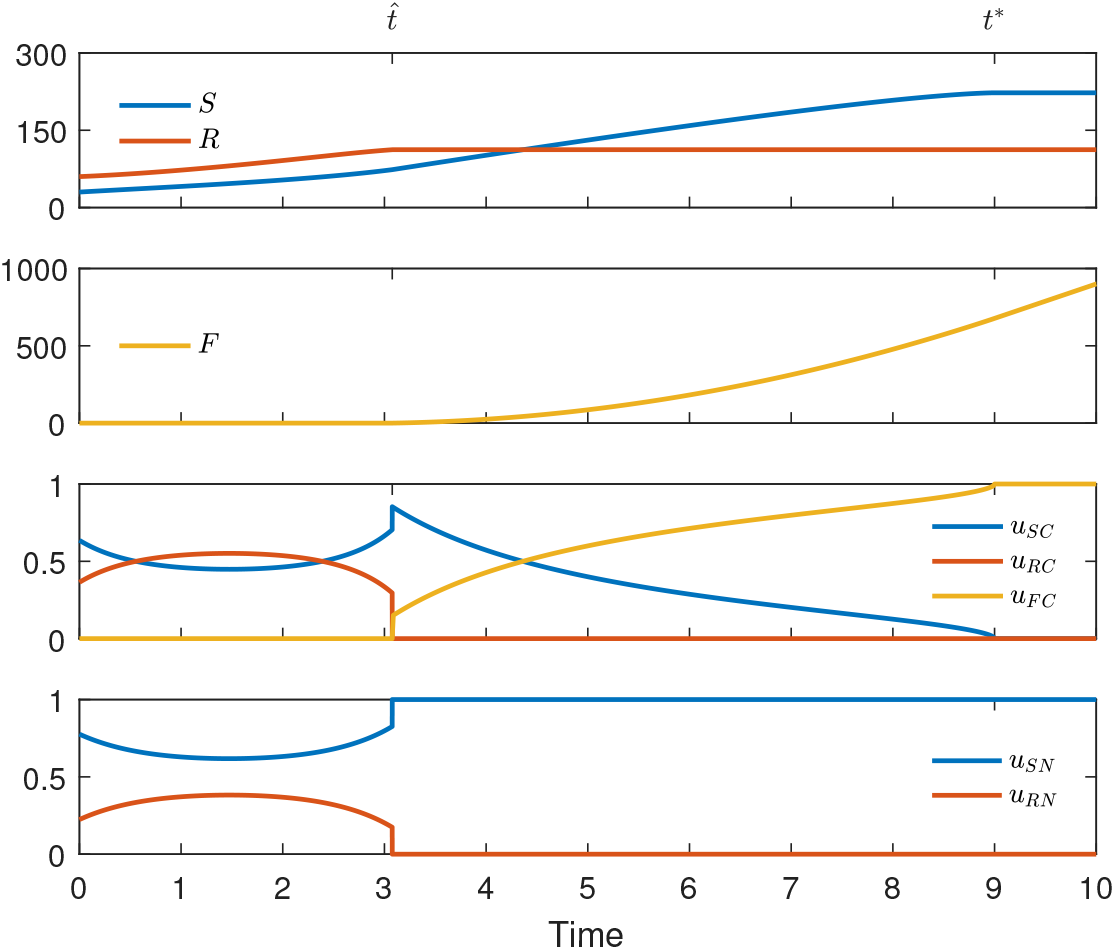
Type *S* initially balanced growth. From top to bottom: Shoots and roots, fruits, carbon controls, nitrogen controls. Shoots, roots, and fruits units of carbon and controls dimensionless

### 7.4 Balanced Growth First - Type R

Here we chose terminal conditions *S*\* = 222.60 and *R*\* = 112.14. This results in *F*(*T*) = 900, *S*_0_ = 36.16, and *R*_0_ = 49.58. The states and controls are shown in Figure 7. Here we see another example of a plant that begins in balanced growth. Unlike Figure 6, the balanced growth phase seen here has a similar structure to that in Figure 5, where the plant begins with root-only growth. In both examples, the phase consists of steadily increasing allocation to shoots while simultaneously decreasing allocation to roots. In this case the plant is initially more biased toward root deficiency than shoot deficiency and so the early growth sees a greater investment in roots. We refer to this type of initially balanced growth as ‘Type R.’

**Figure 7:**
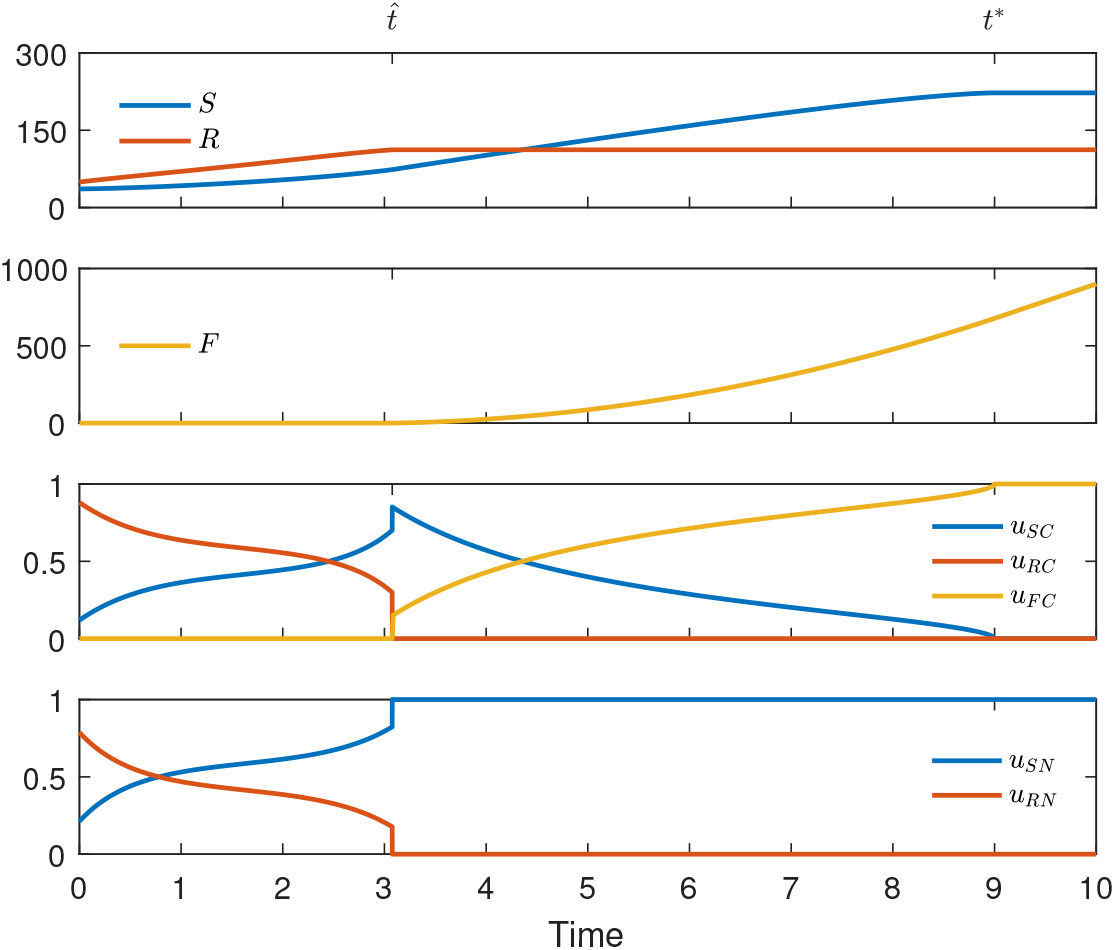
Type R initially balanced growth. From top to bottom: Shoots and roots, fruits, carbon controls, nitrogen controls. Shoots, roots, and fruits in units of carbon and controls dimensionless

### 7.5 Optimal Fruit Yield Contours

In order to better understand the relationship between initial conditions and optimal fruit yield, we looked for points in the (*S*_0_, *R*_0_)-plane for which *F*(*T*) is 700, 800, or 900. For a given value of *S*_0_, we used MATLAB to find the corresponding value of *R*_0_ for which the numerical scheme outlined in Section 6 would yield either 700, 800, or 900 for *F*(*T*). We plotted the resulting contours in the (*S*_0_, *R*_0_)-plane as seen in Figure 8. We also note that the four ‘representative’ simulations we have examined occur on the *F*(*T*) = 900 contour.

**Figure 8:**
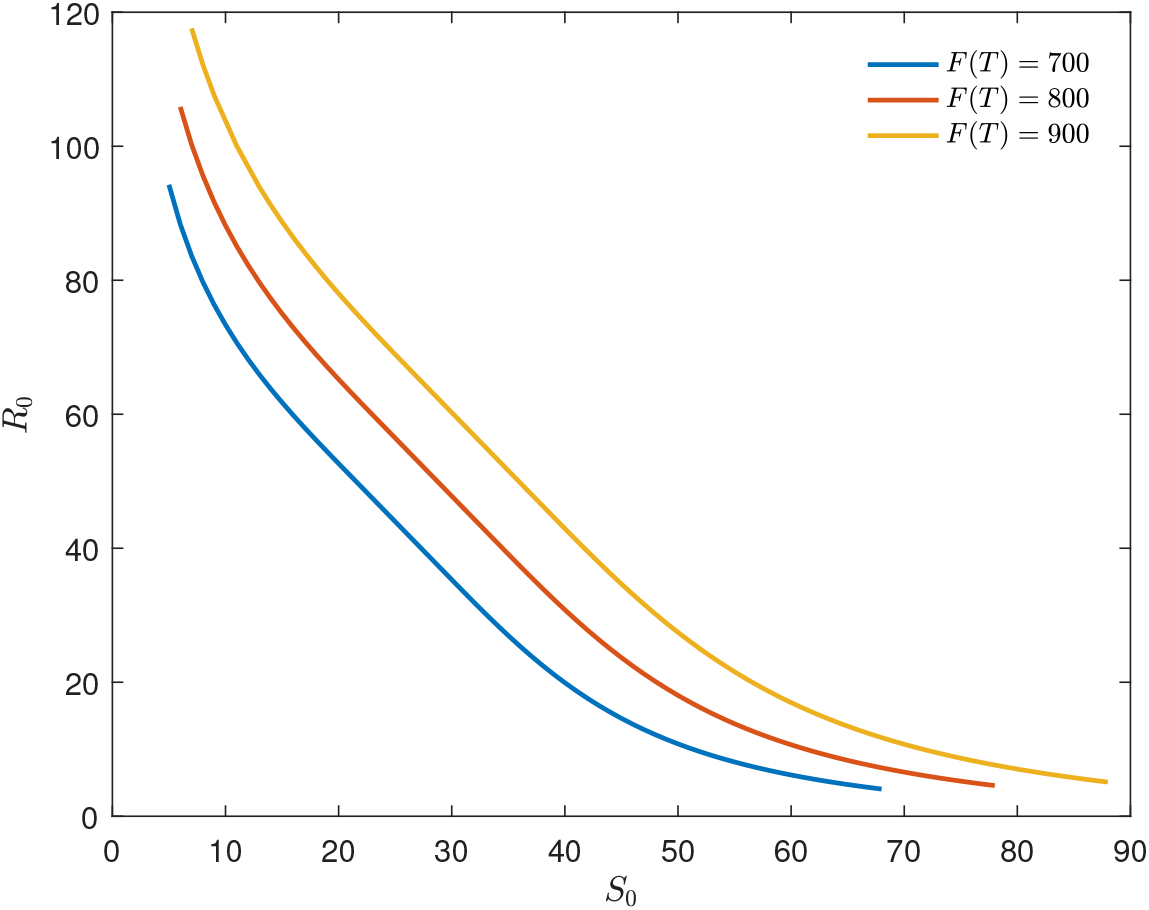
Optimal fruit yield contours for *F*(*T*) = 700,800,900

There are several key features of this plot to notice. First is the wide range of initial conditions for which a particular fruit yield is optimal. Depending on the allocation strategy, plants with very different initial conditions may share an optimal fruit yield. Additionally, for any given point in Figure 8, we can see how much more initial shoots or roots would be necessary to move the plant to a higher-yield contour. Looking at the ends of the contours, it takes relatively little additional initial structure to move from one contour to the next, whereas in the middle of the contours we see that a larger increase in initial biomass is required to move to the next contour. Looking at Figures 6 and 7, we see that these middle regions contain initial conditions from which the plant begins in balance. Nearer to the ends of the contours, where the concavity becomes more pronounced, we see that for the plant must be more biased toward the vegetative organ with the most biomass.

## 8 Discussion

By incorporating nitrogen into the model, but keeping the fruits solely reliant on carbon, we have considered a case in which *ν_F_* → ∞. While not realistic for most annual plants, this provides a mathematically approachable framework in which to begin to analyze optimal allocation of two resources. It is worth noting, however, that if we exchange the assumption that fruits are C-only with the assumption that the fruits are N-only, we expect the model to predict a reversal in the roles of shoots and roots. The assumption that fruits are C-only, however, is a more natural extension of Iwasa and Roughgarden’s work (1984).

An obvious outcome of our model, not present in Iwasa and Roughgarden (1984), is the penultimate interval of mixed shoot/fruit growth. As the *ν_F_* > *ν_S_*, *ν_R_*, we see a phase during which shoot production is more important overall to eventual fruit yield, but an increased capacity to assimilate nitrogen is not useful so any excess carbon is invested in fruits. We also note that this behavior is consistent with recent predictions in the literature (see e.g. Koshkin et al. 2021).

The theoretical results in Section 5 confer a degree of biological relevance to the model that adds credence to the model despite the narrow scope imposed by the assumption that fruits are built solely from carbon. In particular, we showed that *z_SC_*, *z_RC_*, and *u_FC_* are continuous between any two phases in which they are defined and non-zero. Note that while *z_SC_* and *z_RC_* are dimensionless, they are multiples of the ratio of C flux to N flux in shoots in roots. So, the fact that *z_SC_* and *z_RC_* are continuous here means that these ratios are continuous in the shoots and roots. This is striking, given that, while these individual fluxes are continuous at the beginning of balanced growth, they are markedly discontinuous at the end in Figures 4, 5, 6, and 7. So, as long as either shoots or roots is growing, the amount of allocated C per unit of allocated N varies continuously, which is reasonable to expect from biochemical processes. Furthermore, by (34), the fact that *u_FC_* is continuous between the penultimate and final intervals means that the rate of fruit growth is continuous here.

The results presented in Section 7 provide several avenues for drawing more general conclusions about the nature of plant growth that optimizes fruit yield. In Figures 4, 5, 6, and 7, we see that allocation is a balancing act between avoiding limitation and investing in the organ which most directly contributes to increased fruit yield. The plant invests in the most deficient organ first until it can efficiently invest in both shoots and roots, which it does so that by the end of balanced growth we see increased C flux to shoots. During the penultimate interval, then, fruits are a better investment than roots, but fruits still benefit from increased C flux. Therefore, the plant continues to invest in shoots while gradually transitioning to fruit-only growth. These ideas are also reminiscent of our discussion of the marginal values and necessary conditions for optimality in Section 4.3, in that they suggest that two organs can grow simultaneously only if neither SU can out perform the other.

Recall that Figures 4, 5, 6, and 7 represent ‘snapshots’ of optimal growth along the *F*(*T*) = 900 contour represented in Figure 8. While the general trends discussed above hold across the contour, there is a broad spectrum of optimal strategies. What is striking is the high level of variation in solutions with initial conditions along the contour, and the fact that these initial transients are absent by time *T*. We see this in the values of *S*(*T*), *R*(*T*), and *F*(*T*) in Figures 4, 5, 6, and 7, but the trend generally holds along the contour. In some sense, then, we can think of optimal growth under this model as being an equalizing agent that reduces initial variation in a population.

## A Necessary Conditions

In this appendix we will derive the necessary conditions presented in (29) in Section 3.4. Note that because the carbon controls do not directly depend on the nitrogen controls we can obtain the necessary conditions by considering an *n*-state two-control problem and an *n*-state three-control problem separately. In our particular situation *n* = 3, but we derive the conditions in more generality because there is minimal complexity added to the derivation. In all the following we consider *f*, **g** to be continuously differentiable in all arguments and **u** to be piecewise continuous. We begin with an *n*-state two-control problem in Appendix A.1 followed by an *n*-state three-control problem in Appendix A.2

### A.1 *n* States, 2 Controls, Interval [0, *T*]

We consider the optimal control problem give by (137), with the goal of deriving the associated necessary conditions.

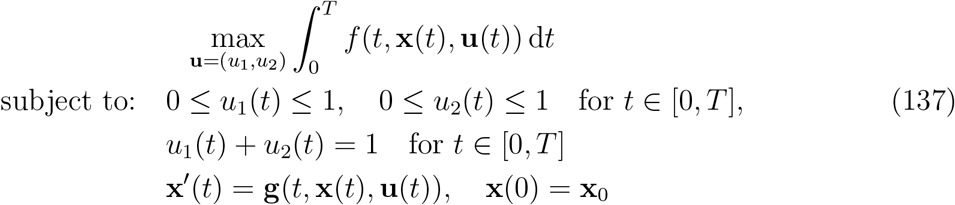

To this end, let *U* be the space of admissible controls, and define the functional 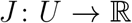 by

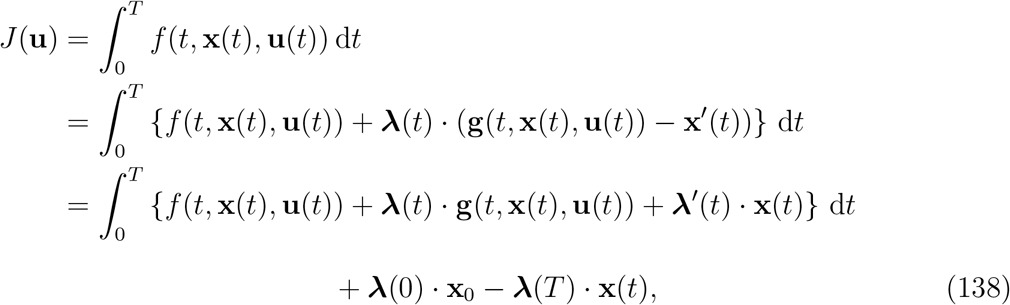

where **λ**(*t*) is a piecewise differentiable function to be specified later, and the final equality was obtained by integrating by parts. Note that by **λ**’ we mean the function whose *n*^th^ component is the time derivative of the *n*^th^ component of **λ**. We use the same notation for **x**′(*t*). Let **u**\*, **x**\* be optimal, and for a piecewise continuous variations *h*_1_, *h*_2_ and 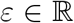 define 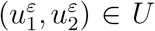 by 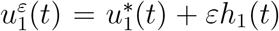 and 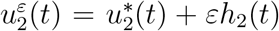, and let **x**^ε^ be the corresponding state. Note that admissibility requires 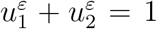, so we have *h*_1_ = −*h*_2_. Because **u**\*, **x**\* are optimal, we have

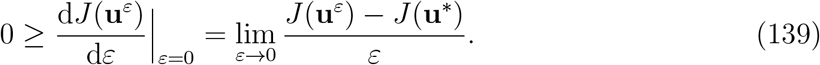

We have inequality rather than equality above because the boundedness of the controls makes it possible that optimality is only global rather than local, and the numerator in the difference quotient is nonpositive because *J*(**u**\*) is maximal over admissible controls. We differentiate (138) with respect to *ε* and note that conditions on the functions involved in the integrand allow us to interchange the derivative and integral via the Dominated Convergence Theorem. Suppressing the arguments from here on,

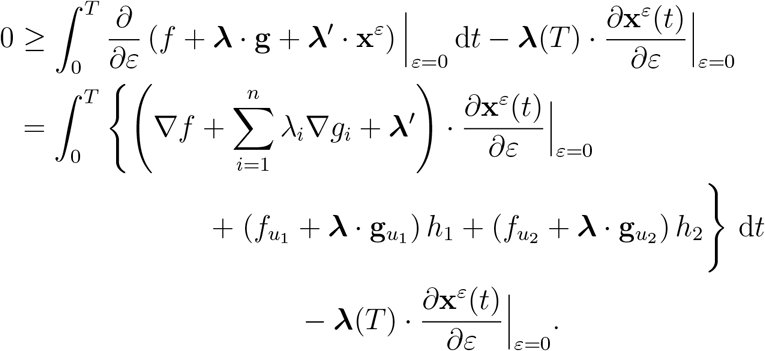

Note that here we are using ∇*f* to mean the vector of derivatives of *f* with respect each component of **x**, and likewise for ∇*g_i_*. Choosing **λ** to satisfy

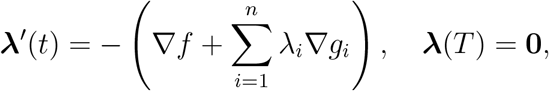

the inequality reduces to

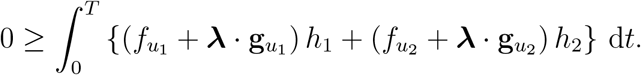

Using the fact that *h*_1_ = – *h*_2_, we obtain

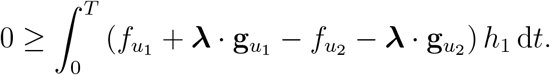

Next, we use this inequality to obtain necessary conditions for optimality. Let *s* be a point of continuity for 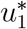 and 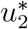 such that 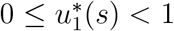, and suppose for the sake of contradiction that *f*_*u*_1__ + **λ** · **g**_*u*_1__ – *f*_*u*_2__ – **λ** · **g**_*u*_2__ > 0 at *s*. As **λ** is continuous, and *f* and **g** are continuously differentiable, by the continuity of 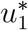 and 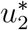 at *s* we have that *f*_*u*_1__ + **λ** · **g**_*u*_1__ – *f*_*u*_2__ – **λ** · **g**_*u*_2__ is continuous at *s* as well. Therefore we can find a closed interval *I* about *s* such that 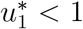 and *f*_*u*_1__ + **λ** · **g**_*u*_1__ – *f*_*u*_2__ – **λ** · **g**_*u*_2__ > 0 on *I*. Let

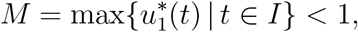

and choose *h*_1_ and *h*_2_ to be

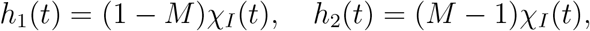

where *χ_I_* is the characteristic function on *I*. Note that this gives us

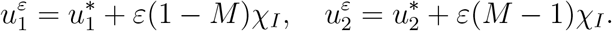

As 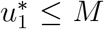 on *I*, then it must be the case that 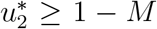 on *I* since 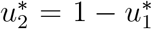. This means that for all *ε* ∈ [0,1] we have

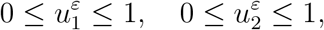

and as the variations were chosen such that *h*_1_ + *h*_2_ = 0, we have that these variations lead to admissible controls. We have then

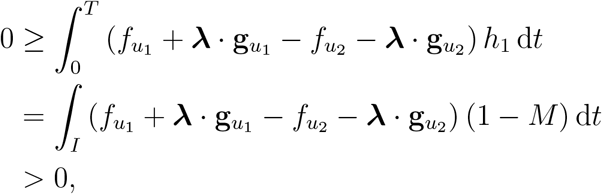

a contradiction. So, it must be the case that *f*_*u*_1__ + **λ** · **g**_*u*_1__ – *f*_*u*_2__ – **λ** · **g**_*u*_2__ ≤ 0. Note that because the controls sum to one we have that 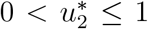 implies 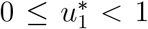, and so we actually have the following condition

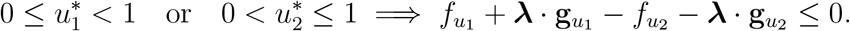

Interchanging the roles of *u*_1_ and *u*_2_ in the argument above also gives us

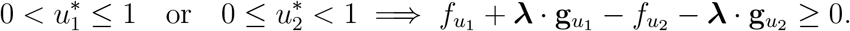

So, together we have the that

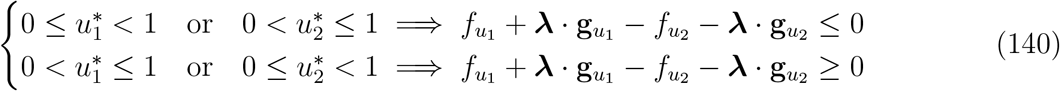

Now, suppose that we are in the case where *f*_*u*_1__ + **λ** · **g**_*u*_1__ – *f*_*u*_2__ – **λ** · **g**_*u*_2__ < 0. By (140) we can rule out the possibility that either 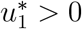 or 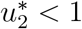 because either choice would lead to a contradiction. Therefore, in this case we can conclude that 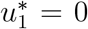 and 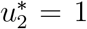. Similar arguments lead to the following conditions:

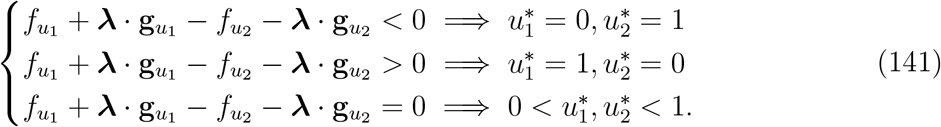

We can convert this into conditions involving a Hamiltonian as follows. Define *H* by

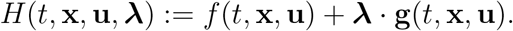

For distinct indicies *i, j* ∈ {1, 2}, rewriting (141) in terms of *H* yields the necessary conditions

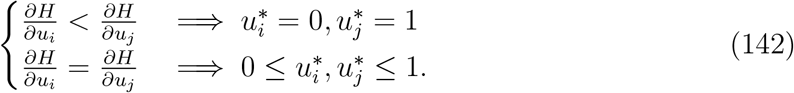

We can also express the differential equations for **x** and **λ** in terms of the Hamiltonian as

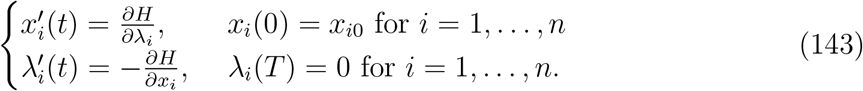

Note that if we were to reduce the problem to one control by writing *u*_2_ = 1 – *u*_1_ these conditions would become exactly the standard necessary conditions for a problem with one bounded control and *n* states (see Lenhart and Workman 2007). Going through this derivation with two controls, however, gives us a starting point to approach the analogous problem with three controls.

### A.2 *n* States, 3 Controls, Interval [0, *T*]

We now consider the following optimal control problem with *n* states and three controls. We will again derive the necessary conditions, following a similar procedure to that used in the simpler case with only two controls. The problem is stated as follows.

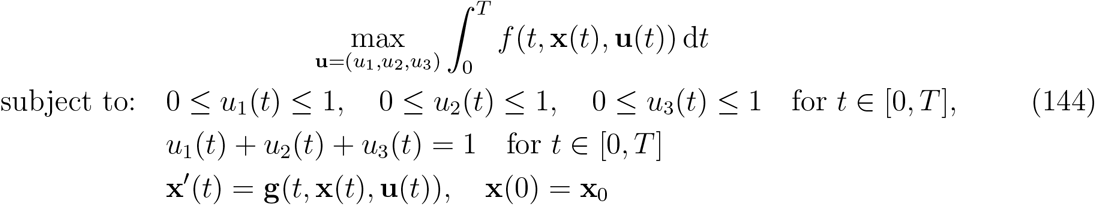

The derivation of the necessary conditions is similar to the two-control case. We let *U* be the space of admissible controls for (144) and define 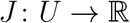 by (138), restated below.

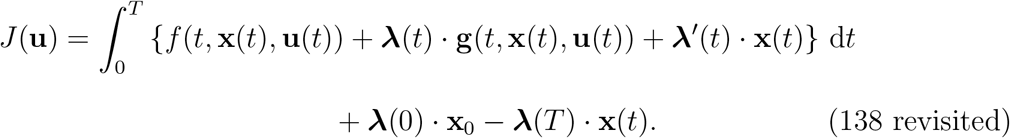

Suppose that **u**\*, **x**\* are optimal, and let *h*_1_, *h*_2_, and *h*_3_ be piecewise continuous variations. Then for 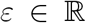 we define **u**^*ε*^ ∈ *U* by 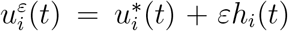 for *i* = 1, 2, 3, and let **x**^*ε*^ be the corresponding state. Admissibility here requires that 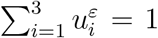 so we have that *h*_3_ = −(*h*_1_ + *h*_2_). By the optimality of **u**\*, **x**\* we again get (139), restated below:

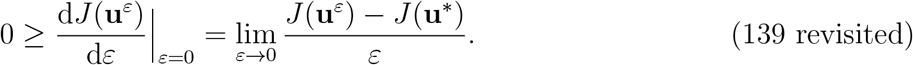

We again differentiate (138), using the DCT to interchange the order of differentiation and integration, this time arriving at

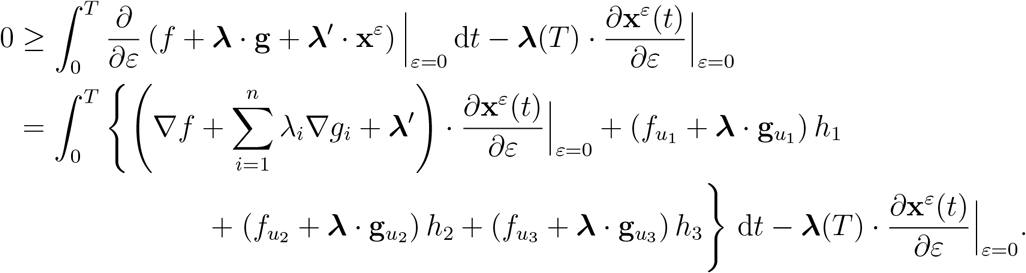

As before, we choose **λ** so that

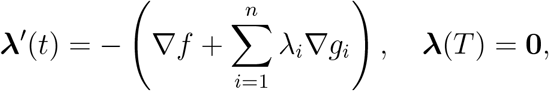

and so we arrive at the inequality

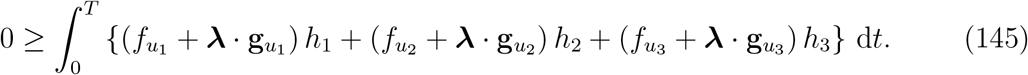

We will use this inequality to derive the necessary conditions for optimality. Due to the similarity between the various cases we will only show one in detail. We begin by using the substitution *h*_3_ = – (*h*_1_ + *h*_2_) to rewrite (145) as

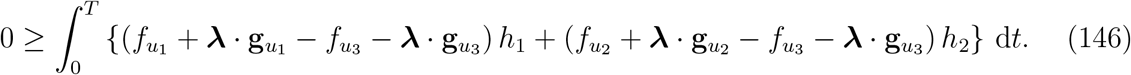

Next, let *s* be a point of continuity for all controls so that 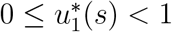 and 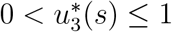. Note that because the controls sum to one this also means that 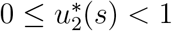. Additionally, assume for the sake of contradiction that *f*_*u*_1__ + **λ** · **g**_*u*_1__ – *f*_*u*_3__ – **λ** · **g**_*u*_3__ > 0 at *s*. As **λ** is continuous, and *f* and **g** are continuously differentiable, by the continuity of the controls we have that *f*_*u*_1__ + **λ** · **g**_*u*_1__ – *f*_*u*_3__ – **λ** · **g**_*u*_3__ is continuous at *s* as well. Therefore, we can find a closed interval *I* about *s* such that 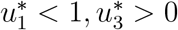, and *f*_*u*_1__ + **λ** · **g**_*u*_1__ – *f*_*u*_3__ – **λ** · **g**_*u*_3__ > 0 on *I*. Next, let

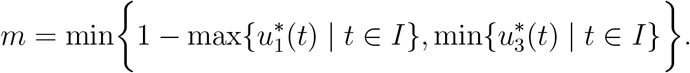

and note that *m* > 0. Next, we choose variations to be

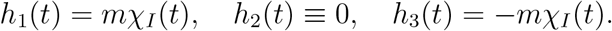

This gives us the controls

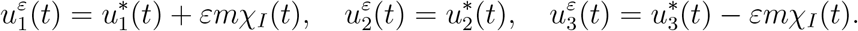

Note that because 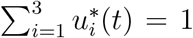 and 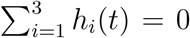 we have that 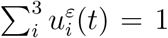 as well. Furthermore, restricting our attention to *t* ∈ *I* and *ε* ∈ [0, 1], we have

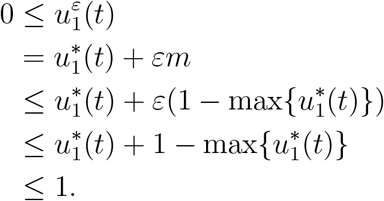

As this bound also holds outside of *I* by assumption we have that 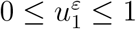. Now, for 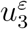, again restricting our attention to *t* ∈ *I* and *ε* ∈ [0,1] we have

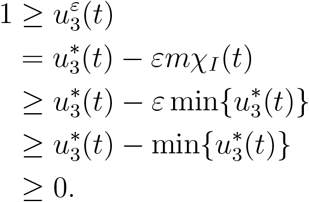

Again, as this bound holds outside of *I* by assumption we have that 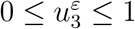. Therefore, the controls 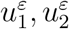, and 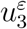 are admissible for all *ε* ∈ [0,1]. We have then

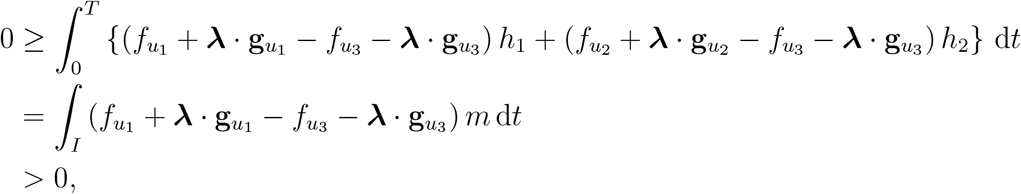

a contradiction. Therefore it must be the case that *f*_*u*_1__ + **λ** · **g**_*u*_1__ – *f*_*u*_3__ – **λ** · **g**_*u*_3__ ≤ 0.

Now, by swapping the indicies 1 and 2 in the argument above we can reach the additional conclusion that *f*_*u*_2__ + **λ** · **g**_*u*_2__ – *f*_*u*_3__ – **λ** · **g**_*u*_3__ ≤ 0 for the same set of assumptions. Note that this is because if we assume that 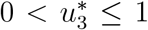 then we can conclude both 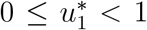 and 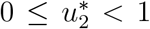 regardless of which assumption is used for a particular argument. Therefore, we have that if we are in the case that 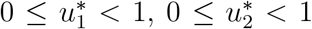, and 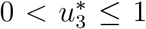 then we have that both *f*_*u*_1__ + **λ** · **g***u*_1_ – *f*_*u*_3__ – **λ** · **g**_*u*_3__ ≤ 0 and *f*_*u*_2__ + **λ** · **g**_*u*_2__ – *f*_*u*_3__ – **λ** · **g**_*u*_3__ ≤ 0. We can permute the roles of *u*_1_, *u*_2_, and *u*_3_, or, equivalently, repeat the argument above with the substitutions of *h*_1_ = −(*h*_3_ + *h*_2_) or *h*_2_ = −(*h*_3_ + *h*_1_) into (145) to obtain the following.

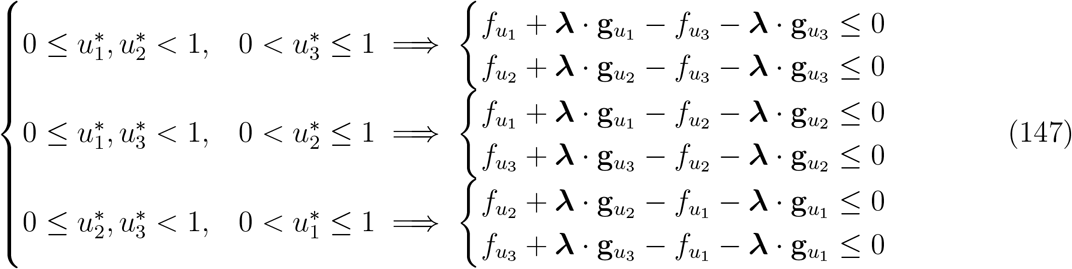

We will now work to develop implications going in the other direction. We will illustrate this for a few cases and the remaining cases follow from permuting the controls. First, consider the case that at a point *t* ∈ [0, *T*] we have *f*_*u*_1__ + **λ** · **g**_*u*_1__ – *f*_*u*_3__ – **λ** · **g**_*u*_3__ < 0 and *f*_*u*_2__ + **λ** · **g**_*u*_2__ – *f*_*u*_3__ – **λ** · **g**_*u*_3__ < 0, and suppose for the sake of contradiction that 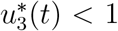. Then it must be the case that at *t* either 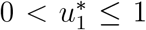 and 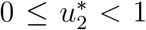 or 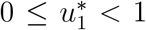 and 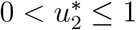. Using (147) we see that the first case implies that *f*_*u*_3__ + **λ** · **g**_*u*_3__ – *f*_*u*_1__ – **λ** · **g**_*u*_1__ ≤ 0 at *t* and the second case implies that *f*_*u*_3__ + **λ** · **g**_*u*_3__ – *f*_*u*_2__ – **λ** · **g**_*u*_2__ ≤ 0 at *t*, both of which contradict our assumptions. So, it must be the case that 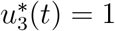.

Next, consider the case that at a point *t* ∈ [0, *T*] we have *f*_*u*_1__ + **λ** · **g**_*u*_1__ – *f*_*u*_3__ – **λ** · **g**_*u*_3__ < 0 and *f*_*u*_1__ + **λ** · **g**_*u*_1__ – *f*_*u*_2__ – **λ** · **g**_*u*_2__ < 0, and suppose for the sake of contradiction that 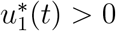. Then it must be the case that at *t* we have 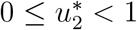 and 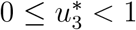. Using (147) we see that this implies that *f*_*u*_2__ + **λ** · **g**_*u*_2__ – *f*_*u*_1__ – **λ** · **g***u*_1_ ≤ 0 and *f*_*u*_3__ + **λ** · **g**_*u*_3__ – *f*_*u*_1__ – **λ** · **g***u*_1_ ≤ 0 at *t*, both of which contradict our assumptions here. Therefore, it must be the case that 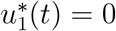. A similar argument can be used to show that if *f*_*u*_2__ + **λ** · **g**_*u*_2__ – *f*_*u*_1__ – **λ** · **g**_*u*_1__ < 0 and *f*_*u*_2__ + **λ** · **g**_*u*_2__ – *f*_*u*_3__ – **λ** · **g**_*u*_3__ < 0 at a point *t* ∈ [0, T] then it must be the case that 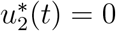.

By permuting the roles of the controls in the arguments above we can reach the following conclusions. For distinct indicies *i*, *j*, *k* ∈ {1, 2, 3} we have that

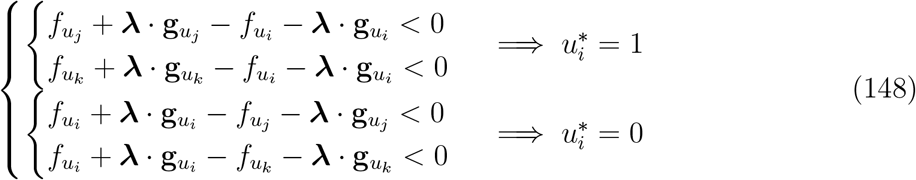

Note that this also tells us that the only way for a control to take on a value other than zero or one is for at least one of the inequalities in (148) to be an equality instead. We can convert this into conditions involving a Hamiltonian as follows. Define *H* by

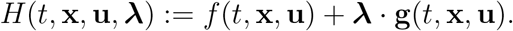

Rewriting (148) in terms of *H* yields the necessary conditions

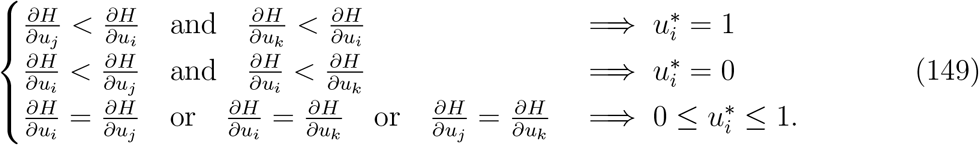

We can also express the differential equations for **x** and **λ** in terms of the Hamiltonian:

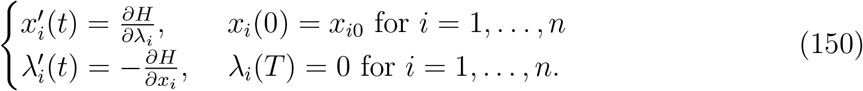

## B Penultimate Interval Proofs

In this appendix we include the proofs of Lemma 3, Lemma 4, Lemma 5, and Theorem 6.

### B.1 Existence of Penultimate Interval

#### Lemma 10.

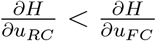 *in an open interval immediately prior to t*\*.

*Proof*. Because λ*_R_*(*t*\*) = 0, λ*_R_* is continuous, and *G*′ is bounded, there exists *ε* > 0 such that for all *t* in (*t*\* – *ε*, *t*\*) we have

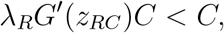

which in terms of the Hamiltonian (35) gives the desired result:

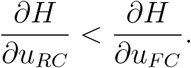

#### Lemma 11.

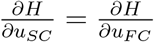 *in an open interval immediately prior to t*\*.

*Proof*. We will begin by showing that

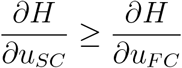

in an open interval immediately prior to *t*\*. Suppose to the contrary that for some *ε* > 0 we have for all *t* in (*t*\* – *ε*, *t*\*) that

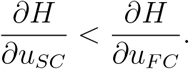

By Lemma 3 we have that

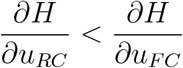

as well, and so by (29) this means that *u_FC_* = 1 and *u_SC_* = 0 = *u_RC_*. As in the final interval, this yields

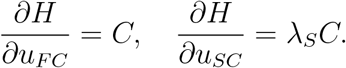

However, as λ*_S_*(*t*\*) = 1 and by (49) we have that 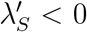, it must be the case that λ*_S_* > 1. This then means that

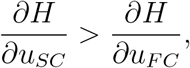

a contradiction. Therefore we have shown that during this interval we have

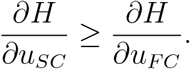

Suppose now that the inequality is strict, and that in fact we have

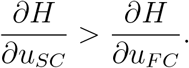

Now, combining this supposition with Lemma 3 gives us

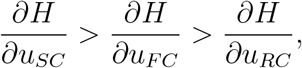

which by (29) means that

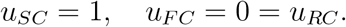

This then gives us

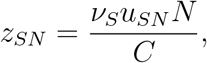

which, for fixed *C* and *N*, is bounded between 0 and 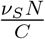. Therefore, we have that *G*′(*z_SN_*) ≠ 0. Furthermore, recall that we know that λ*_R_*(*t*\*) = 0 and *G*′ is bounded. This means that there is a (potentially smaller) open interval immediately prior to *t*\* in which we have

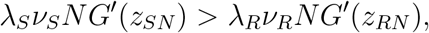

or in terms of the Hamiltonian,

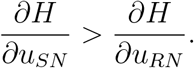

By (29), this means that *u_SN_* = 1 and *u_RN_* = 0 during this interval. With this configuration of the controls, we have that *N* = *N*\* is constant because *R*′ = 0 and this interval reaches this final interval. So, we have that

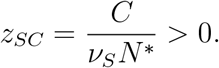

This then means that *G*′(*z_SC_*) < 1, and so as λ*_S_*(*t*\*) = 1 and both *z_SC_* and λ*_S_* are continuous here there exists *ε* > 0 such that for all *t* in (*t*\* – *ε*, *t*\*) we have

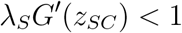

that is

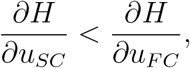

a contradiction. Therefore, we have

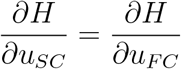

in an open interval immediately prior to *t*\*, as desired.

#### Lemma 12.

*u_SN_* = 1 *and u_SC_* > 0 *in an open interval immediately prior to t*\*.

*Proof*. We will first show that *u_SN_* is non-zero. Recall that by (65) and (29), we have that *u_RC_* = 0 in an open interval immediately prior to *t*\*. Suppose furthermore that *u_SN_* = 0 during this interval as well. By (10) and (11), this means that both *S*′ = 0 and *R*′ = 0. As this interval is followed by a period of fruit-only growth, in order to maximize the integral in (27) it must be the case that *u_FC_* = 1. We therefore have

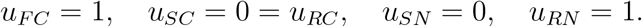

In particular, because here we are again in a case where *u_FC_* = 1, *u_SC_* = 0 = *u_RC_*, and λ*_R_*(*t*\*) = 0 we again arrive at a situation where

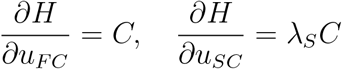

following the same argument as in the justification for the existence of a final interval of fruit-only growth. However, as we know that here λ*_S_* > 1 we have that

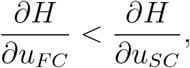

which regardless of the relationship between 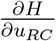 with the other two partial derivatives contradicts the fact that *u_FC_* = 1. Hence, *u_SN_* > 0.

Now, because λ*_S_* > 1 we know by (66) that *z_SC_* > 0 so that *G*′(*z_SC_*) < 1. Because *u_SN_* > 0, this allows us to conclude that *u_SC_* > 0 as well without any concern regarding ambiguous limiting behavior with *G*′(*z_SC_*) when one or both controls are zero. This in turn implies that

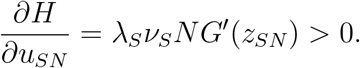

Now, if *u_SN_* < 1 we would also have *u_RN_* > 0. Because *u_RC_* = 0, this would imply that *z_RN_* → ∞, and so by (22) we would have *G*′(*z_RN_*) = 0. This would then imply that

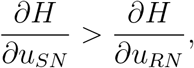

which by (29) means that *u_SN_* = 1, a contradiction. Therefore, it must have been the case that *u_SN_* = 1 in addition to *u_SC_* > 0 in an open interval immediately prior to *t*\*, as desired.

### B.2 Strategy Comparison/Marginal Values

#### Theorem 6.

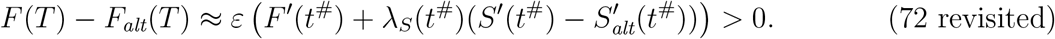

*Proof*. To derive the approximation in Theorem 6, we will first recall a definition we introduced in Section 4.3 when discussing the marginal values. Let *J*(*S*_0_, *R*_0_, *t*_0_) be the maximum for the following optimal control problem.

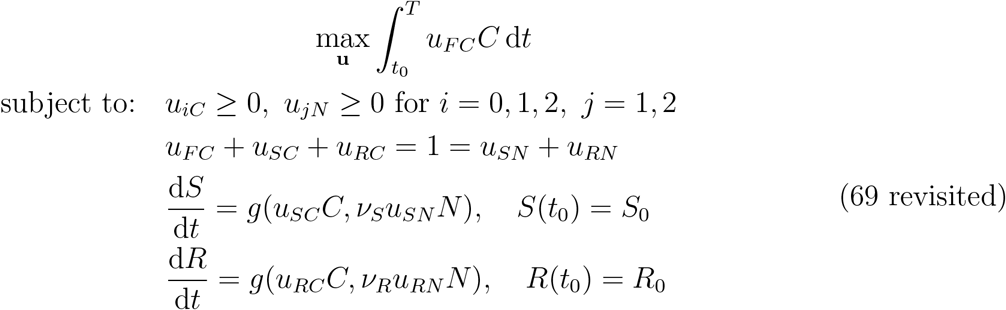

We wish to compute *F*(*T*) – *F*_alt_(*T*), which in terms of *J* is given by

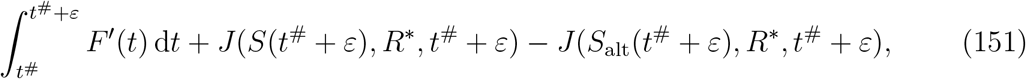

where we use the ‘alt’ subscript wherever we need to distinguish variables which differ from the optimal solution based on the shoot-only growth strategy during [*t*^#^, *t*^#^ + *ε*]. Note also that because here we are in the penultimate interval *R* = *R*\*. We can interpret (151) as the sum of the amount of fruit grown during the interval [*t*^#^, *t*^#^ + *ε*] using the mixed fruit/shoot strategy and the difference between the final value of fruits depending on the different values of *S*(*t*^#^) produced by each strategy on [*t*^#^, *t*^#^ + *ε*]. Now, for small enough *ε*, we have that

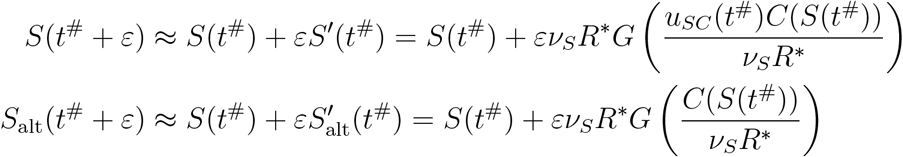

which makes (151) approximately

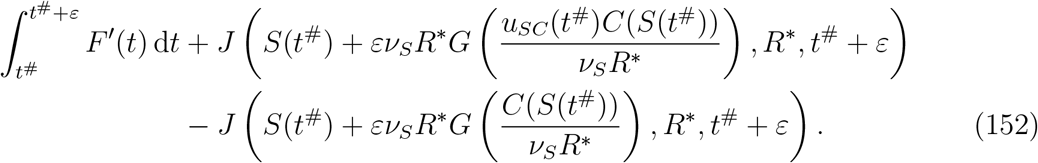

We will simplify the second two terms in (152), using the following labels for convenience:

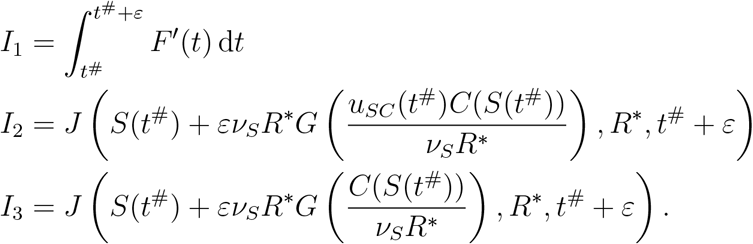

Beginning with *I*_2_, and integrating by parts:

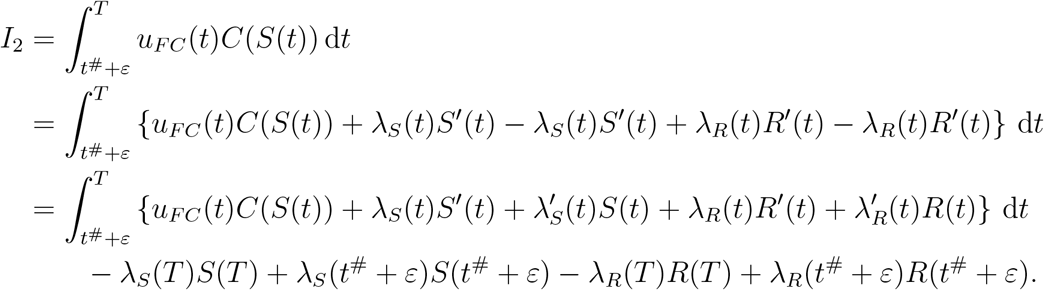

Likewise for *I*_3_, using the same λ*_S_*, λ*_R_*, we have:

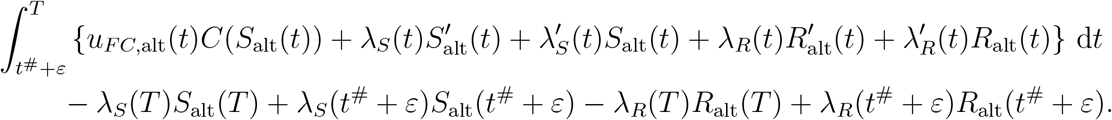

Therefore, *I*_2_ – *I*_3_ becomes

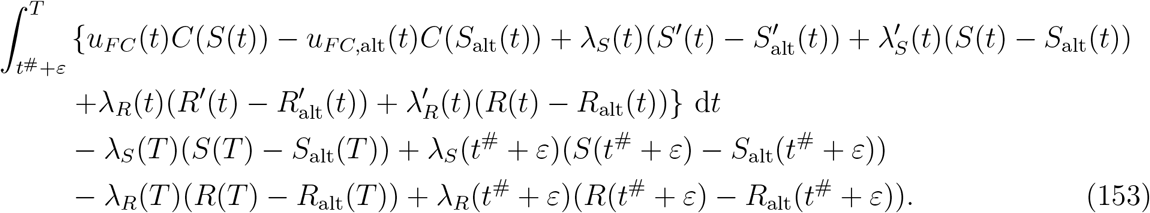

Next, we will approximate the integrand of (153) with it’s Taylor series around the point (*t*, *S*(*t*), *R*(*t*), **u**(*t*)). We only keep track of the first-order terms of the expansion, because as *ε* → 0 the integral already goes to zero. We have then that *I*_2_ – *I*_3_ is approximately

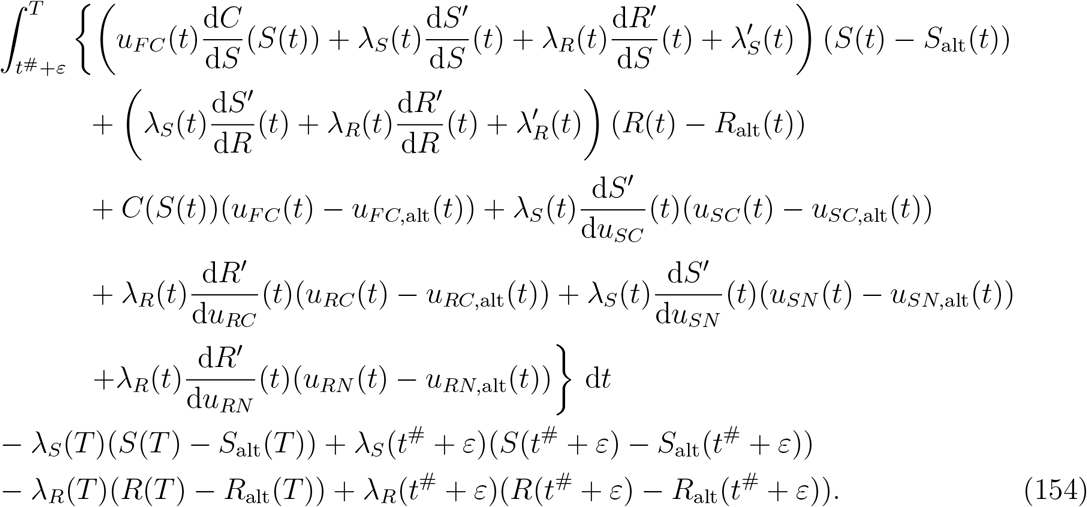

Note that by the differential equations for λ*_S_* and λ*_R_*, and the fact that *R*(*t*^#^ + *ε*) = *R*_alt_(*t*^#^ + *ε*), this simplifies to

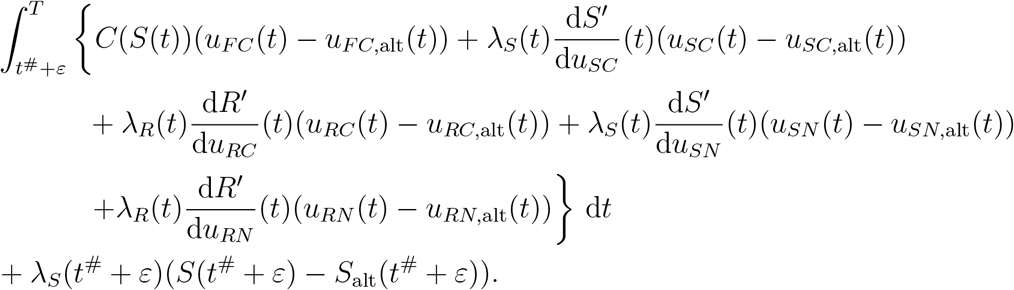

We can rewrite this in terms of the Hamiltonian:

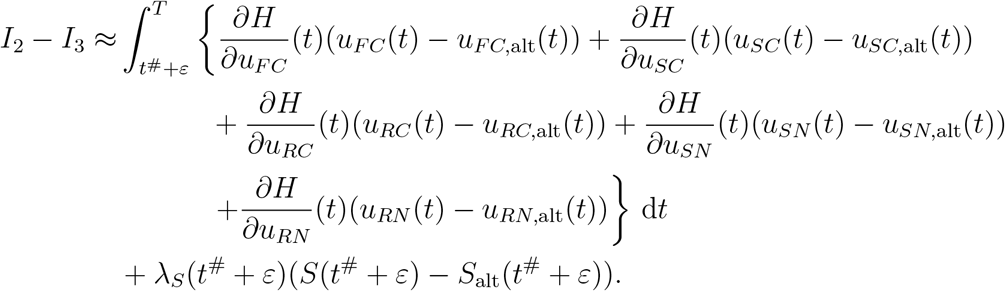

Now, because we know the structure of the optimal solution, namely that during the penultimate interval 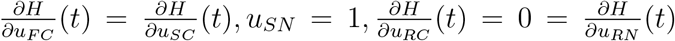, and during the final interval 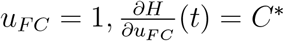, we have that *I*_2_ – *I*_3_ is approximately

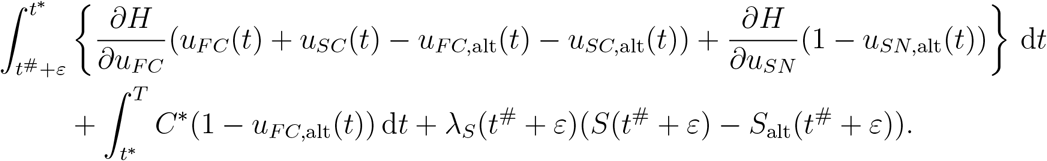

Now, we assume that *ε* is chosen small enough that the optimal solution to the perturbed problem also begins in the penultimate interval at *t* = *t*^#^ + *ε*. This gives us *u*_*FC*,alt_(*t*) + *u*_*SC*,alt_(*t*) = 1 = *u*_*SN*,alt_(*t*), and so

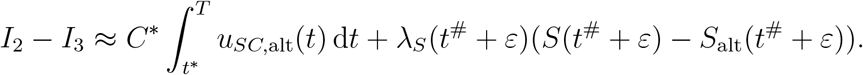

Next, we use

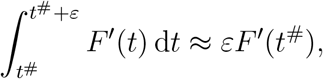

which gives us

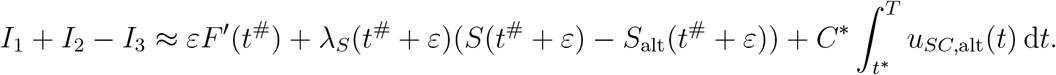

Using a linear approximation, and the fact that *S*(*t*^#^) = *S*_alt_(*t*^#^), we have that to *O*(*ε*)

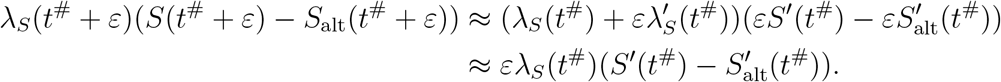

Therefore, we have that

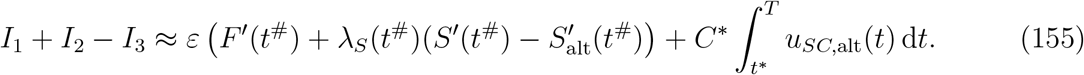

The last step in obtaining the approximation in Theorem 6 is to show that the last term in (155) vanishes. To this end, we will show that 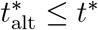, in which case the *u*_*SC*,alt_(*t*) = 0 on [*t*\*, *T*]. To see why this is the case, first note that by the way we constructed the alternate growth path on [*t*^#^, *t*^#^ + *ε*] we know that *S*_alt_(*t*^#^) ≥ *S*(*t*^#^). Because *R*(*t*) = *R*\* = *R*_alt_(*t*) for all *t* in [*t*^#^, *T*], and *S*_alt_ (*t*^#^) ≥ *S* (*t*^#^), it must be the case that *S*_alt_(*T*) ≥ *S* (*T*), else the (*S*, *R*) and (*S*_alt_, *R*_alt_) solutions curves would intersect, and therefore coincide, by the uniqueness of optimal solutions. Now, because *S*_alt_(*T*) ≥ *S*(*T*) and by (15) we know that *C_S_*(*t*) is decreasing, we have that *C_S_*(*S*_alt_(*T*)) ≤ *C_S_*(*S*(*T*)) which by (62) means that 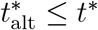. Therefore, the last term in (155) vanishes and we are left with

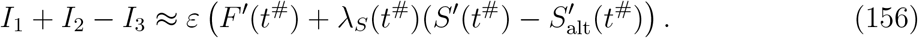

Now that we have obtained (156), it remains to show that *I*_1_ + *I*_2_ – *I*_3_ > 0, i.e. the optimal strategy given by the necessary conditions (shoot/fruit growth) is better than the strategy suggested by the marginal values (shoot-only growth). To this end, dropping the arguments after the first line,

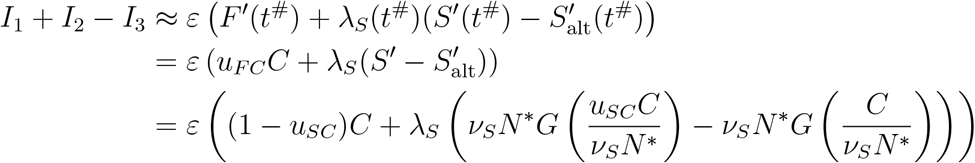

Because here 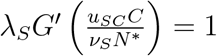 we have

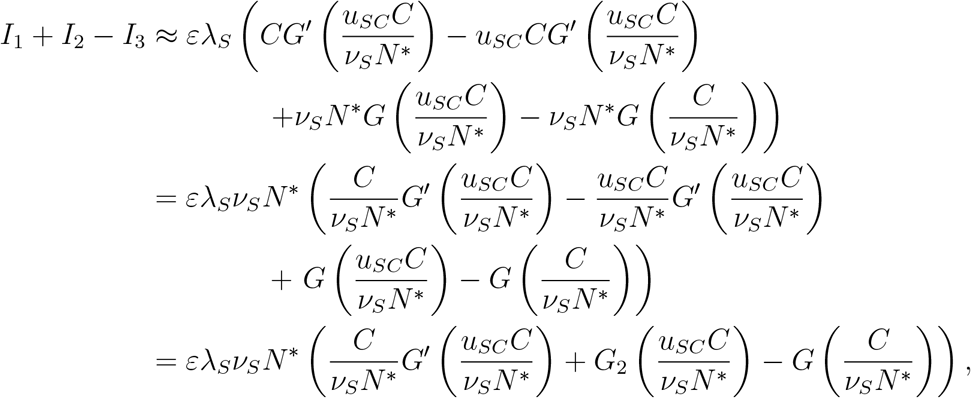

where the last line follows from (24). At this point, showing that for *a* ∈ (0,1), *x* > 0,

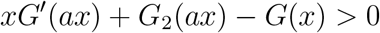

will be sufficient to show that *I*_1_ + *I*_2_ – *I*_3_ > 0. To that end, for *a* ∈ (0,1), *x* > 0,

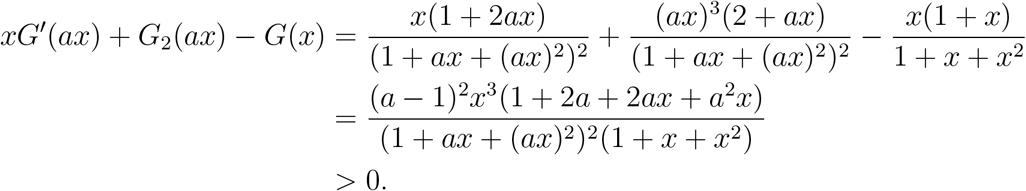

Therefore, we have the desired result, namely that

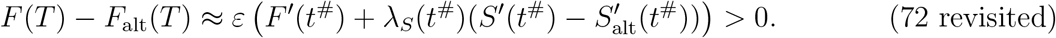

## C Proof of Theorem 8

In this appendix we include the full proof of Theorem 8. Recall that 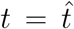 denotes the transition point between the balanced growth phase and the penultimate interval, and 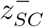 and 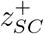 are the left and right limits of *z_SC_* at 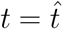, respectively.

### Theorem 8.

*z_SC_ is continuous at* 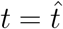.

*Proof*. First, note that by combining (98), (109), and (66), and writing 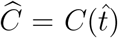, we have that 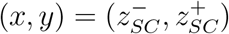 must solve

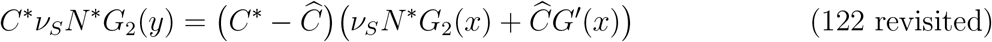

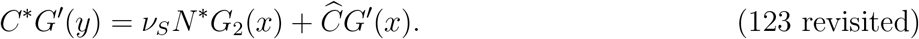

Because *G*_2_ and *G*′ are both one-to-one, we have that *y* is a function of *x* in (122) and (123). It will also be useful to equate the left-hand sides of (122) and (123) to obtain:

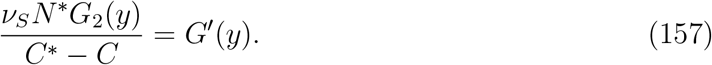

We will break the argument up into three components.

First, we will show that for any given values of 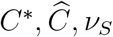, and *N*\*, equation (157) has only one positive real solution. To this end, setting 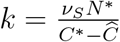, equation (157) gives us

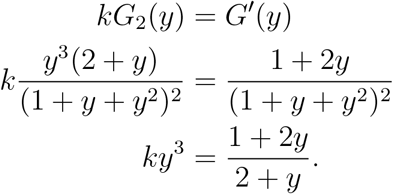

Note that since 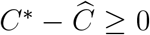 we have that *k* > 0. For any positive choice of *k* it is a simple matter to see that the graphs of

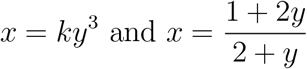

have only one positive intersection point. Note that this tells us that *G*′(*y*)/*G*_2_(*y*) is invertible, and in particular that there is only one admissible value of *y* (depending on *k*) which solves (157). Therefore, 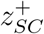 is completely determined by the values of 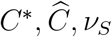, and *N*\*.

Next, we note that *x* = *y* is a solution to equations (122) and (123). We will omit the details of this computation, as for both (122) and (123) it can be verified directly that *x* = *y* is a solution if and only if (157) holds, which we have already established.

Lastly, we will show that *x* = *y* is in fact the only solution to (122) and (123). This, then, combined with the facts that 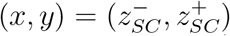 necessarily solves (122) and (123) and 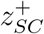 is completely determined by the values of 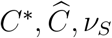, and *N*\* will establish that 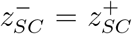, that is *z_SC_* is continuous at 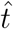.

To show that *x* = *y* is the only solution to (122) and (123) we will use 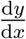 for both these equations as well the PCSU identity (25). We proceed with finding 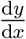 for (122), and so as to avoid confusion we will refer *y* in this equation as *y_A_*. Differentiating (122) gives us

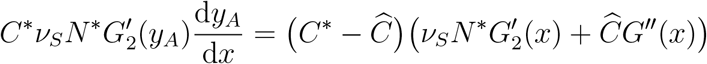

and so

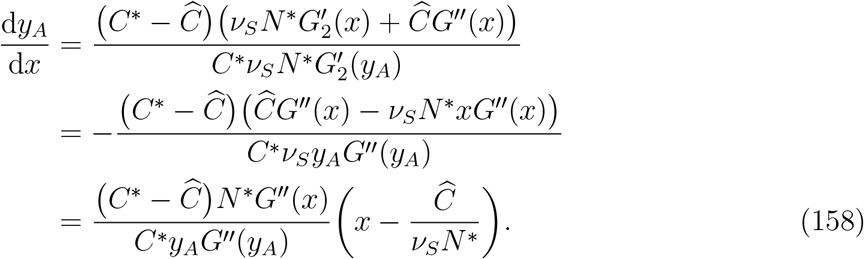

Now,

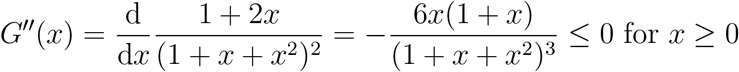

and so

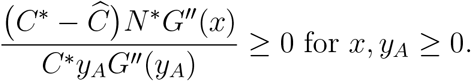

In particular, this means that

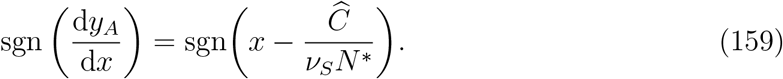

Next, we do the same thing for (123), here using *y_B_* to refer to *y*:

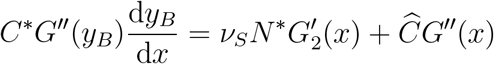

and so

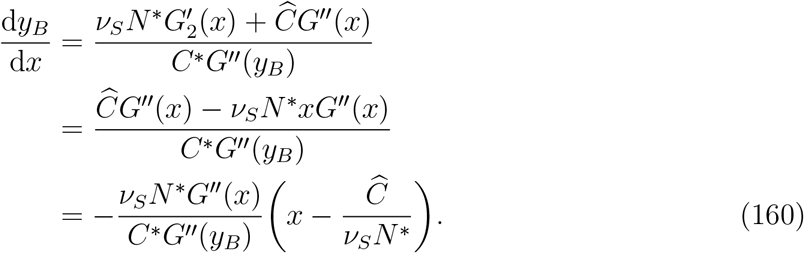

As before,

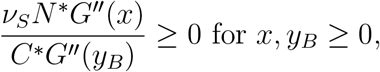

so in particular we have that

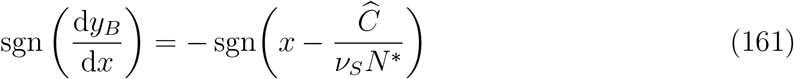

which means that

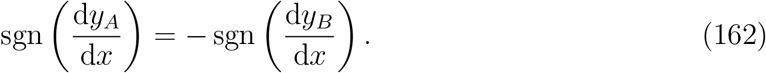

Additionally, note that both derivatives vanish at 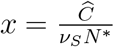. We will use these properties of 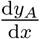 and 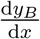 to show that *x* = *y* is the only admissible solution to (122) and (123).

Recall that 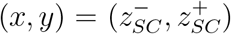 is necessarily a solution to (122) and (123), and we have established that the values of 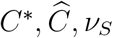, and *N*\* determine a single value of *y*, and therefore 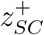, that solves both equations. Now, as 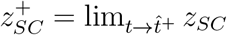, we have that

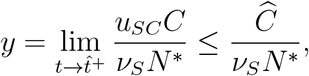

which means that the *y* = *x* intersection point must be at or before 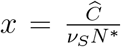, the point where both 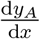 and 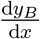 vanish. There are two possible cases to consider: either 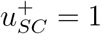 or 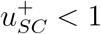, where here 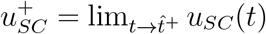.

1. Case 1: 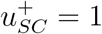 In this case the *y* = *x* intersection point occurs at 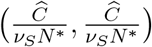. We have from (159) and (161) that 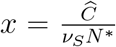 is the unique global minimizer for *y_A_* and the unique global maximizer for *y_B_*, meaning that there are no additional intersection points.
2. Case 2: 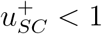 In this case the *y* = *x* intersection point occurs at 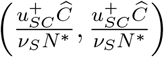, before 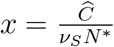 where 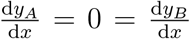. Now, we have from (159) and (161) that *y_A_* strictly decreases until 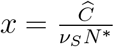 and then strictly increases after that point, whereas *y_B_* strictly increases until 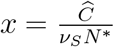 and then strictly decreases after that point, implying the existence of exactly one more intersection point at some 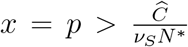. Now, as we established, there is only one solution for *y*, namely 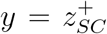. Therefore, in this case there is another intersection point 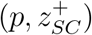 where 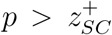. Note that this is potentially admissibly because we would have

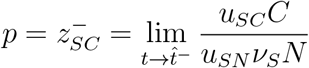

which for admissible choices of the controls could be greater than 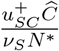. However, if 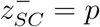, then as *G*′ is strictly decreasing we have that 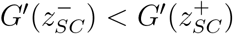, and so

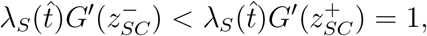

which is impossible by (90). Therefore this second intersection point is not admissible.

In both cases we have shown that the only admissible intersection point for the curves defined by equations (122) and (123) is at the single intersection point where *x* = *y*, which because 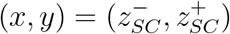 is a solution means that *z_SC_* is continuous at 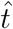.

## D Balanced Growth Differential Equations Derivation

This appendix includes a derivation, referenced in Section 6.5, for the differential equation for *z_SC_* and *z_RC_* during the balanced growth phase. In particular, we will derive equations (134) and (135), revisited below.

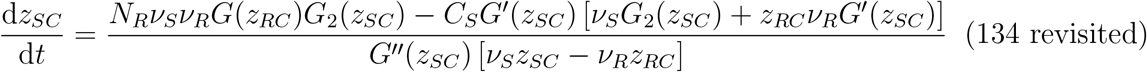

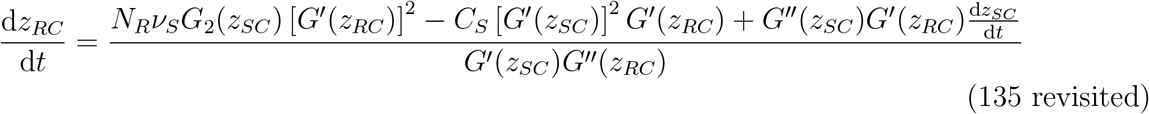

In deriving these differential equations we will use the notation *z_S_* = *z_SC_* and *z_R_* = *z_RC_* for simplicity. We begin by differentiating (90) and (91) with respect to *t*:

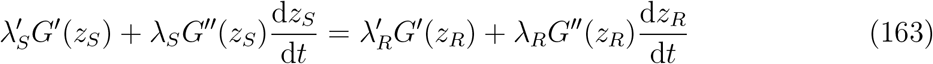

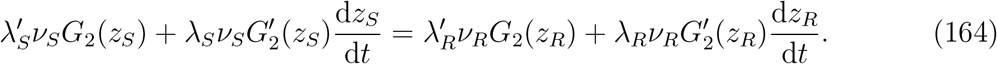

Solving (163) for 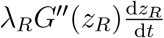 give us

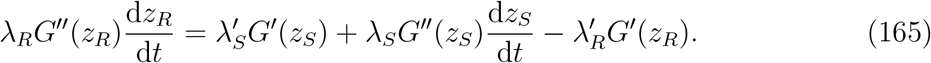

Likewise, making the substitution 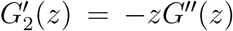 by means of (25), we can solve for 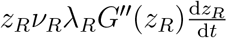 in (164) to get

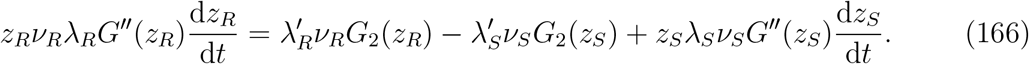

Substituting (165) into (166) gives us

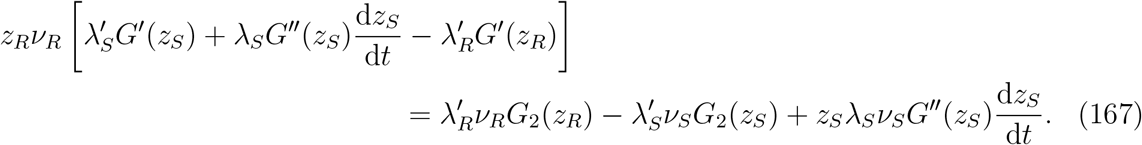

Solving for 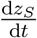, making use of (24), gives us

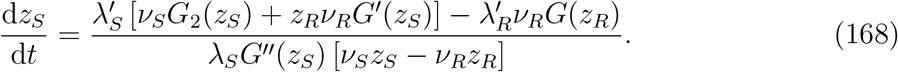

Lastly, then, we use the differential equations for λ*_S_* and λ*_R_* during balanced growth, (94) and (95), to write both 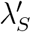 and 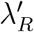 in terms of λ*_S_*. Canceling the resulting common factor of λ*_S_* leaves us with

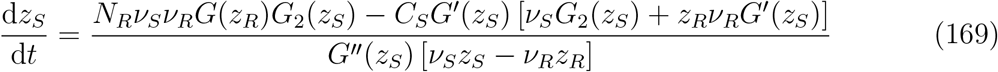

Similarly, we can use (94) and (95) along with (90) to simplify (165), which upon canceling the common factors of λ*_S_* gives us

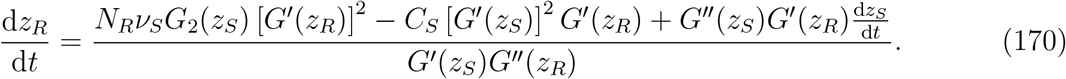

## E MATLAB Script

This appendix includes the MATLAB script for the numerical scheme we developed to solve the optimal control problem (27) associated with the model, as discussed in Section 6. The function `growthpath` is the primary function for constructing the optimal trajectory, and takes the terminal values of shoots and roots as arguments. The primary parameters for `growthpath` are given in Table 1. All functions called by `growthpath` are included at the end of the script. We also note that the script uses slightly different subscript notation than that discussed so far, so we include Table 2 to show the differences.

**Table 1:** Primary parameters for `growthpath`

| Parameter | Value | Meaning |
| --- | --- | --- |
| nu_R | 1 | $\nu_R$ |
| beta | 3 | $\nu_R/\nu_S$ |
| n | $2^{10}$ | RK4 step size is $1/n$ |
| T | 10 | $T$ |
| BGPI_tol | $10^{-10}$ | Tolerance for the balanced growth phase to penultimate interval transition condition (99) |
| IBG_tol | $10^{-10}$ | Tolerance for the initial stage to balanced growth phase transition condition (136) |

**Table 2:**
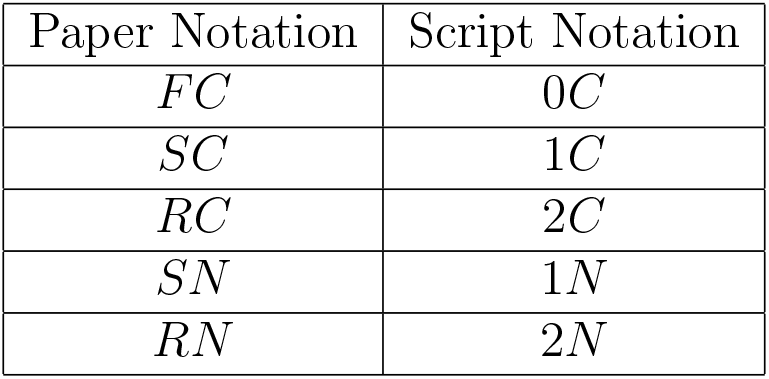
Differences in subscript notation between paper and script

| Paper Notation | Script Notation |
| --- | --- |
| $FC$ | $0C$ |
| $SC$ | $1C$ |
| $RC$ | $2C$ |
| $SN$ | $1N$ |
| $RN$ | $2N$ |

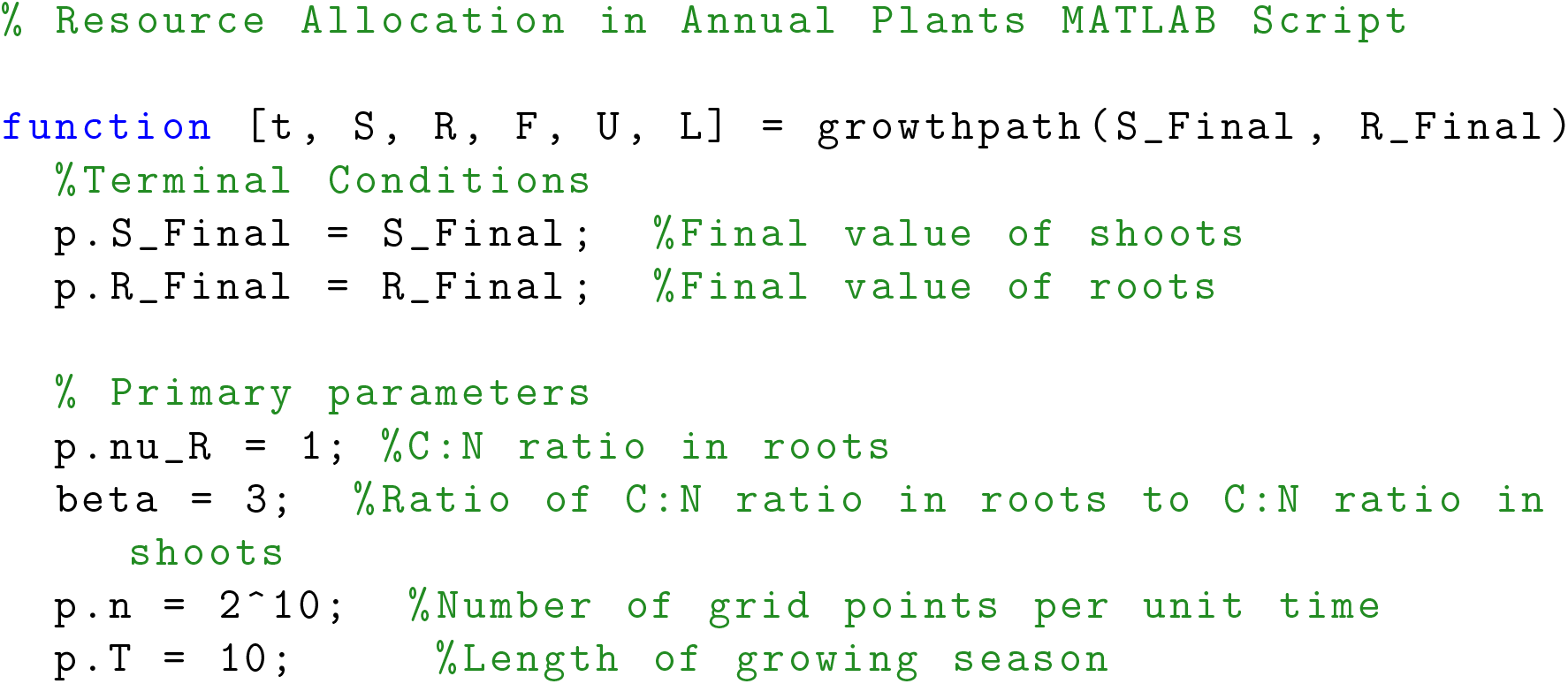
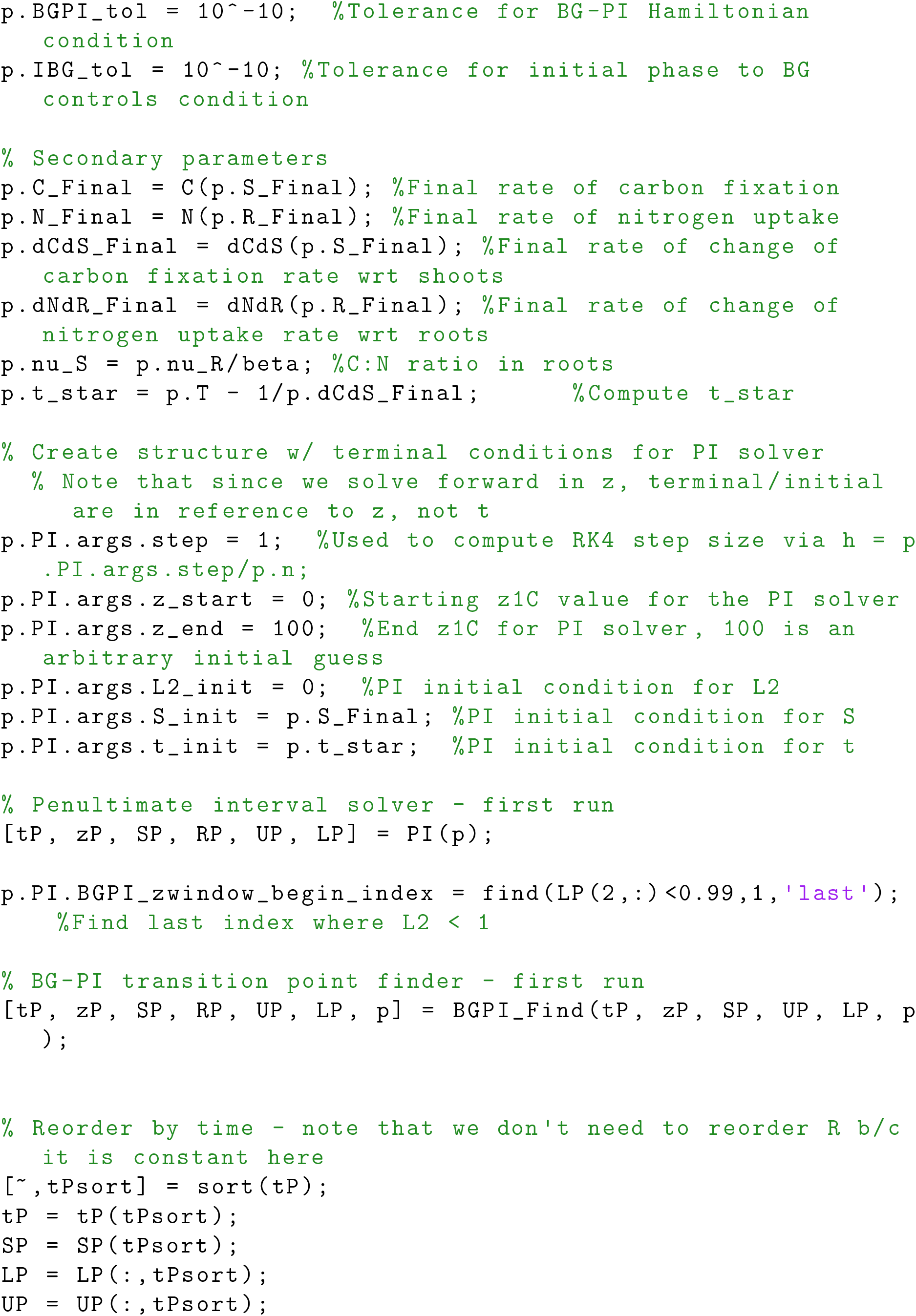
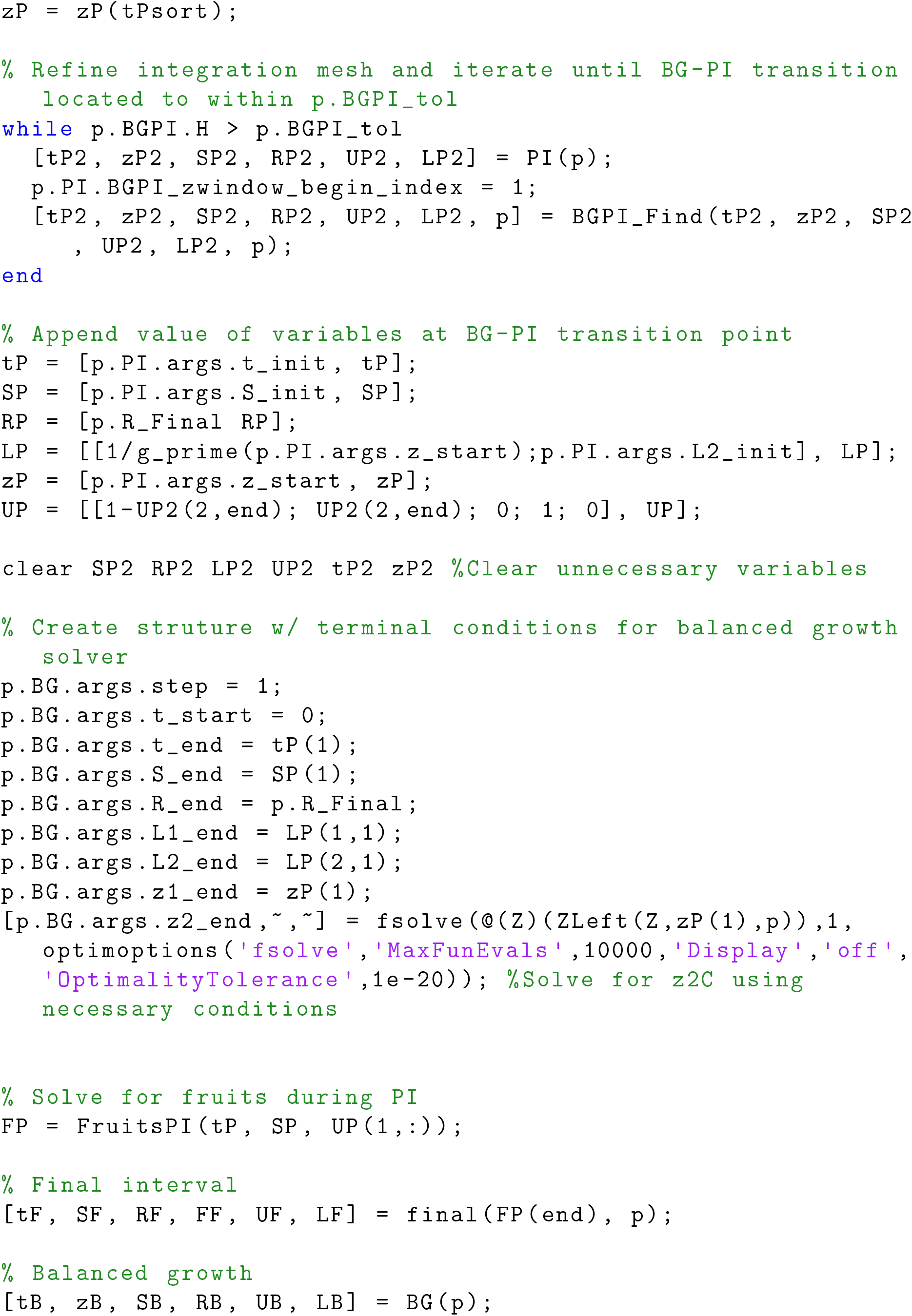
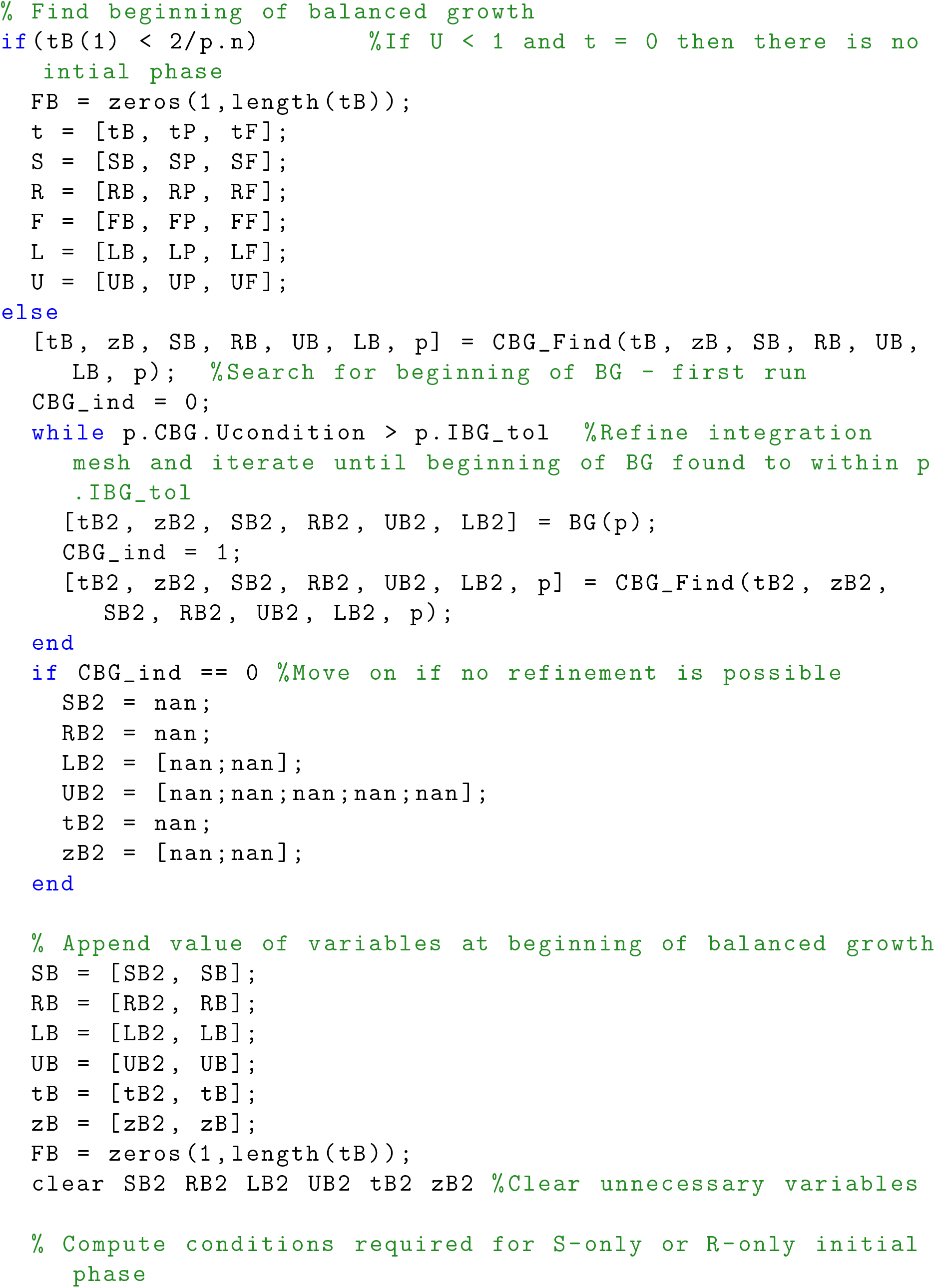
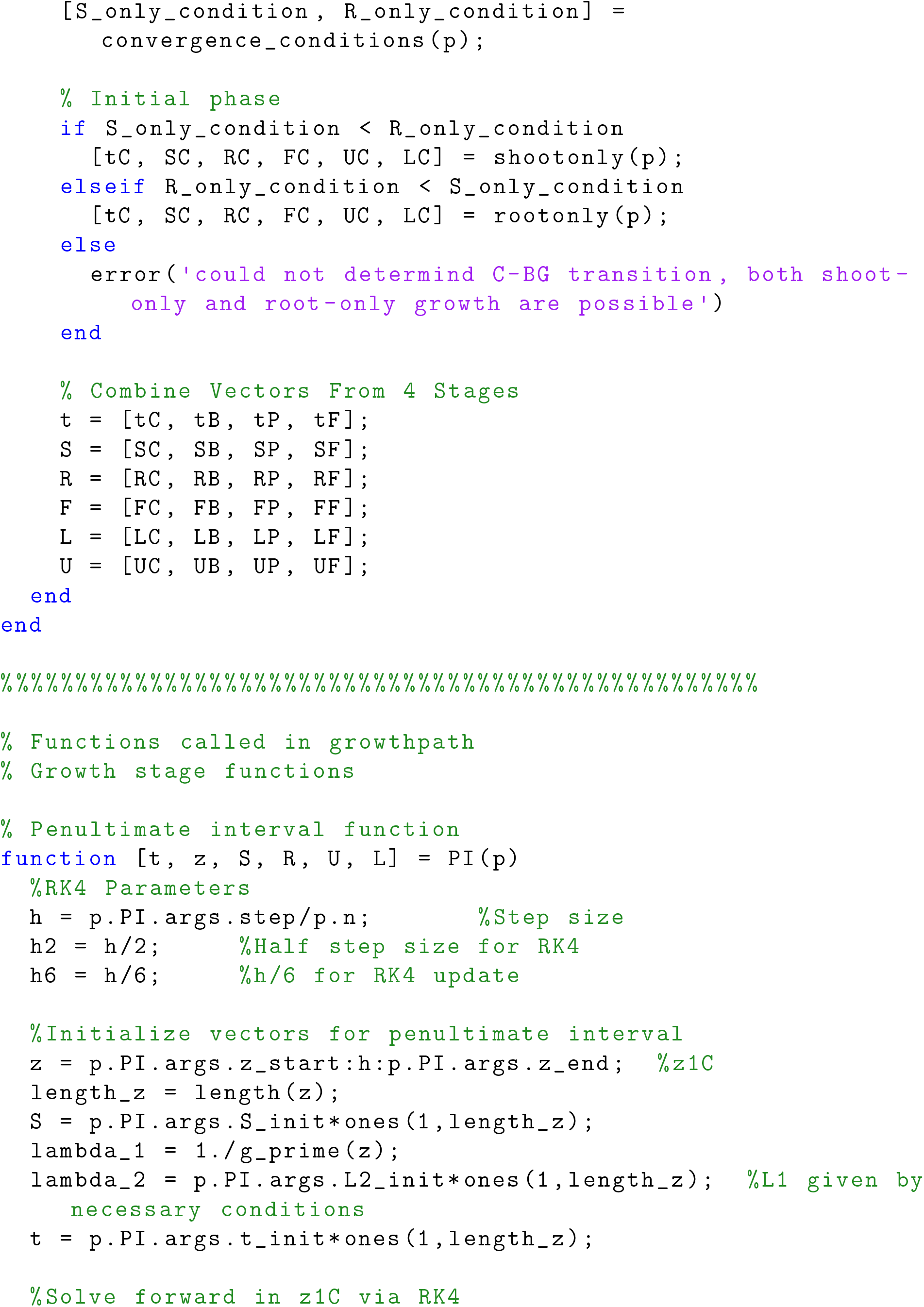
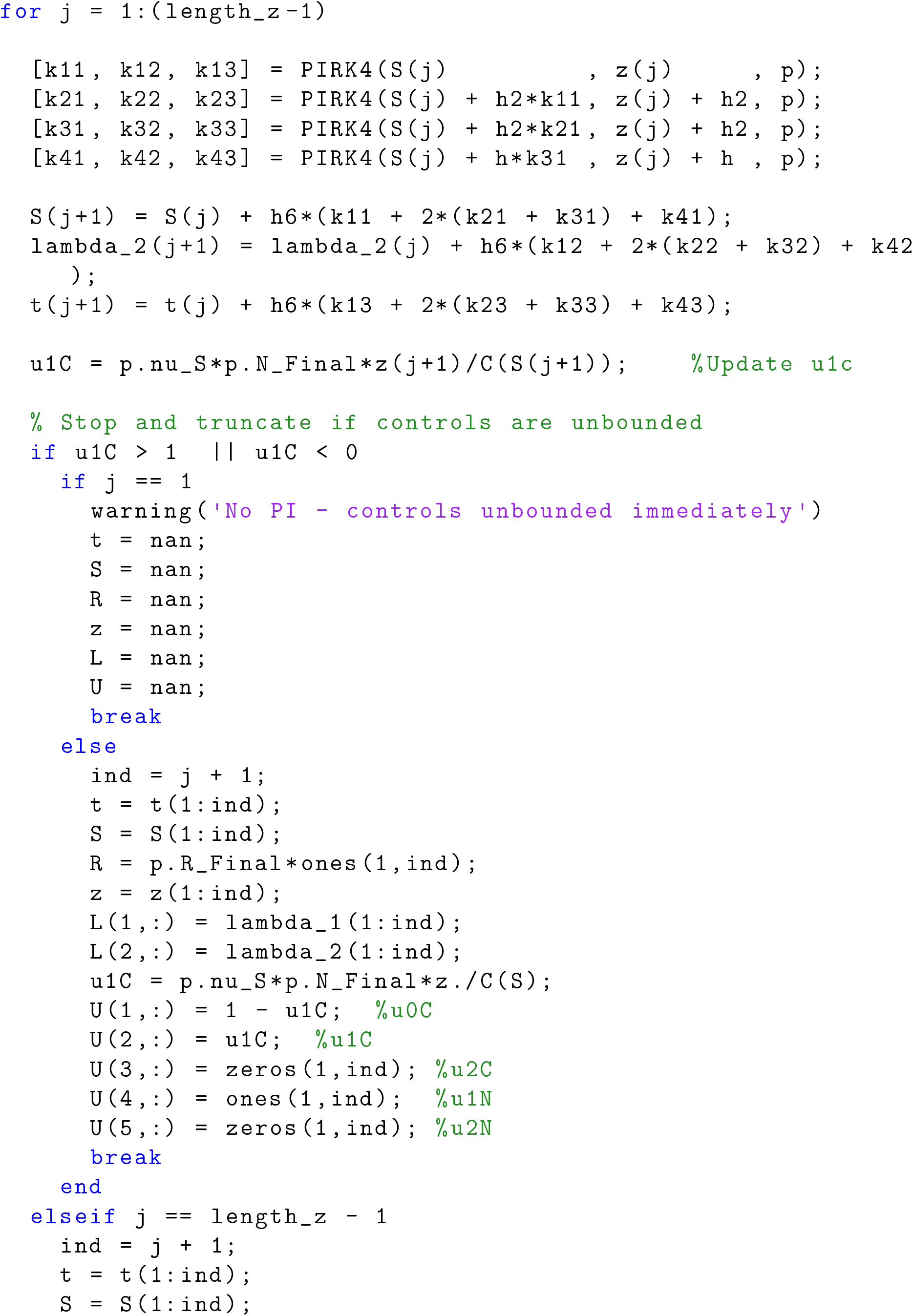
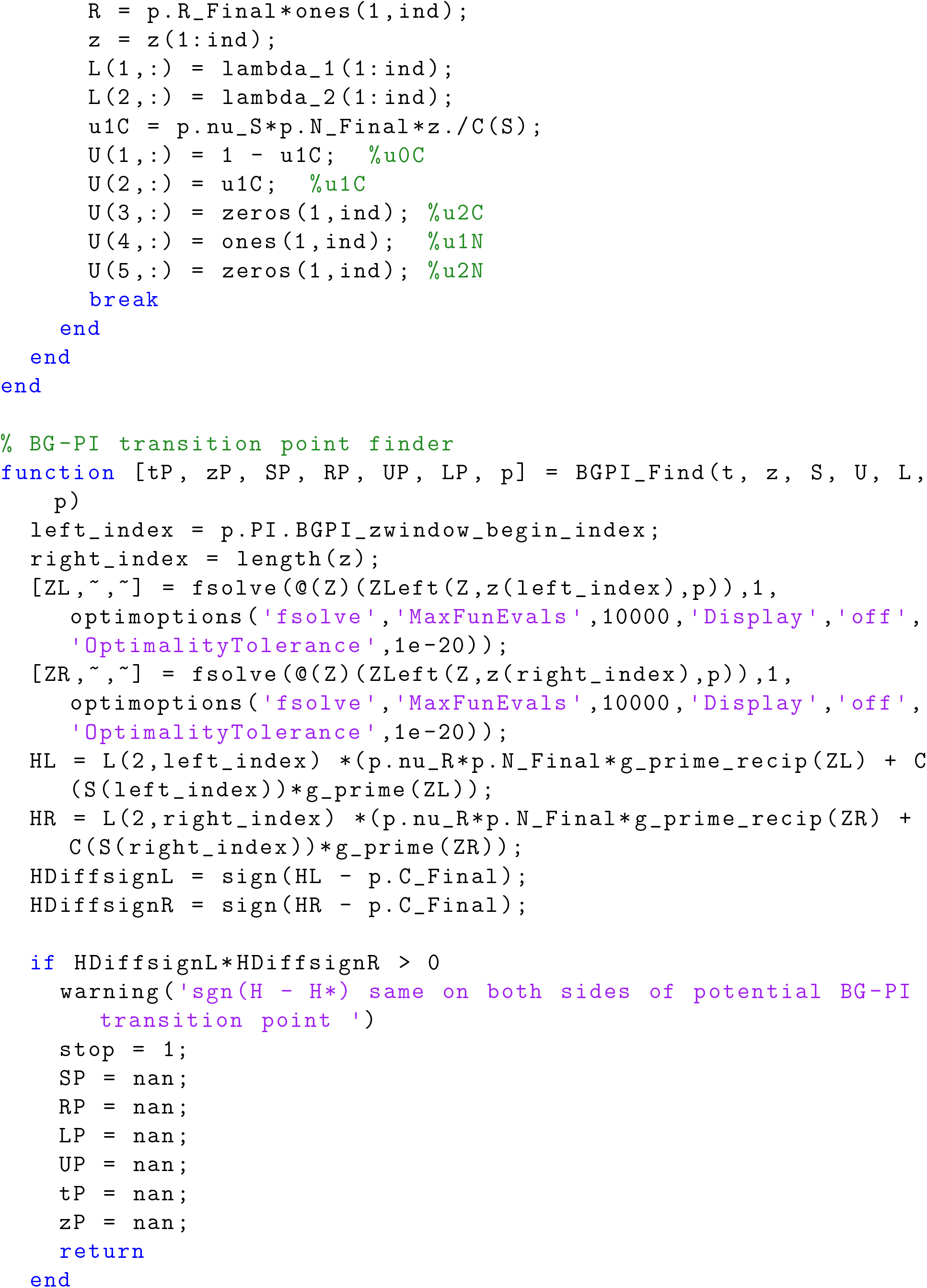
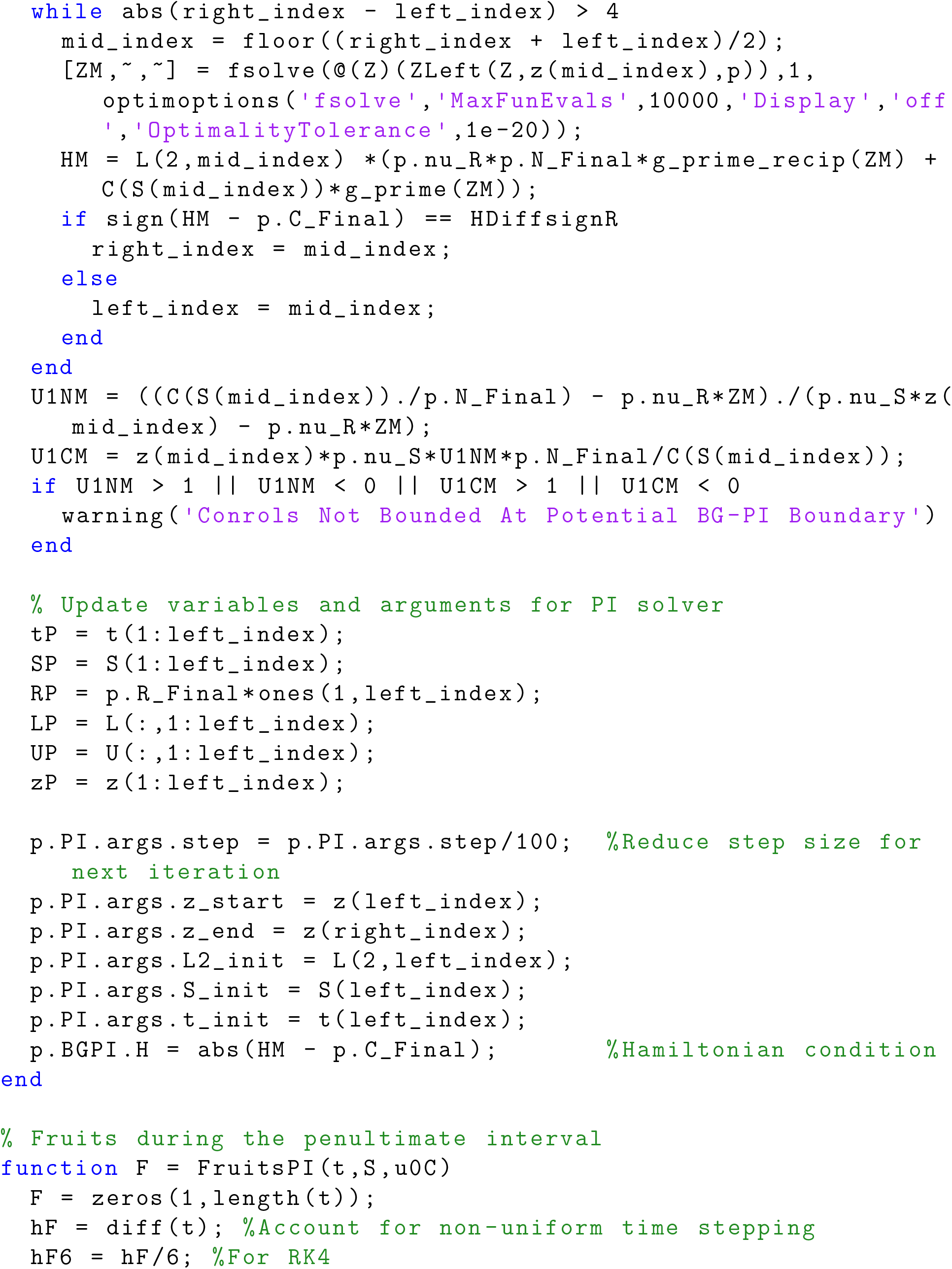
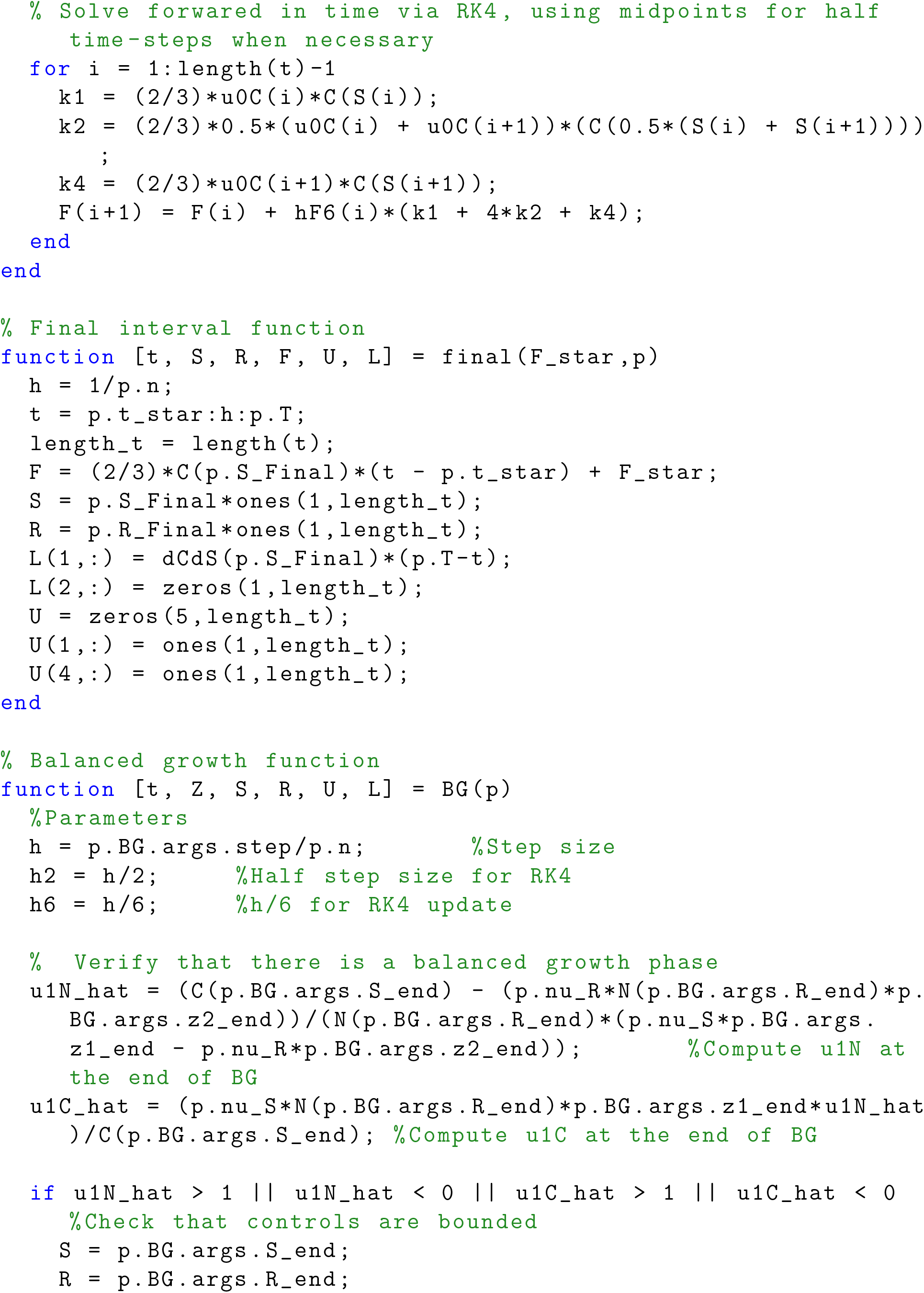
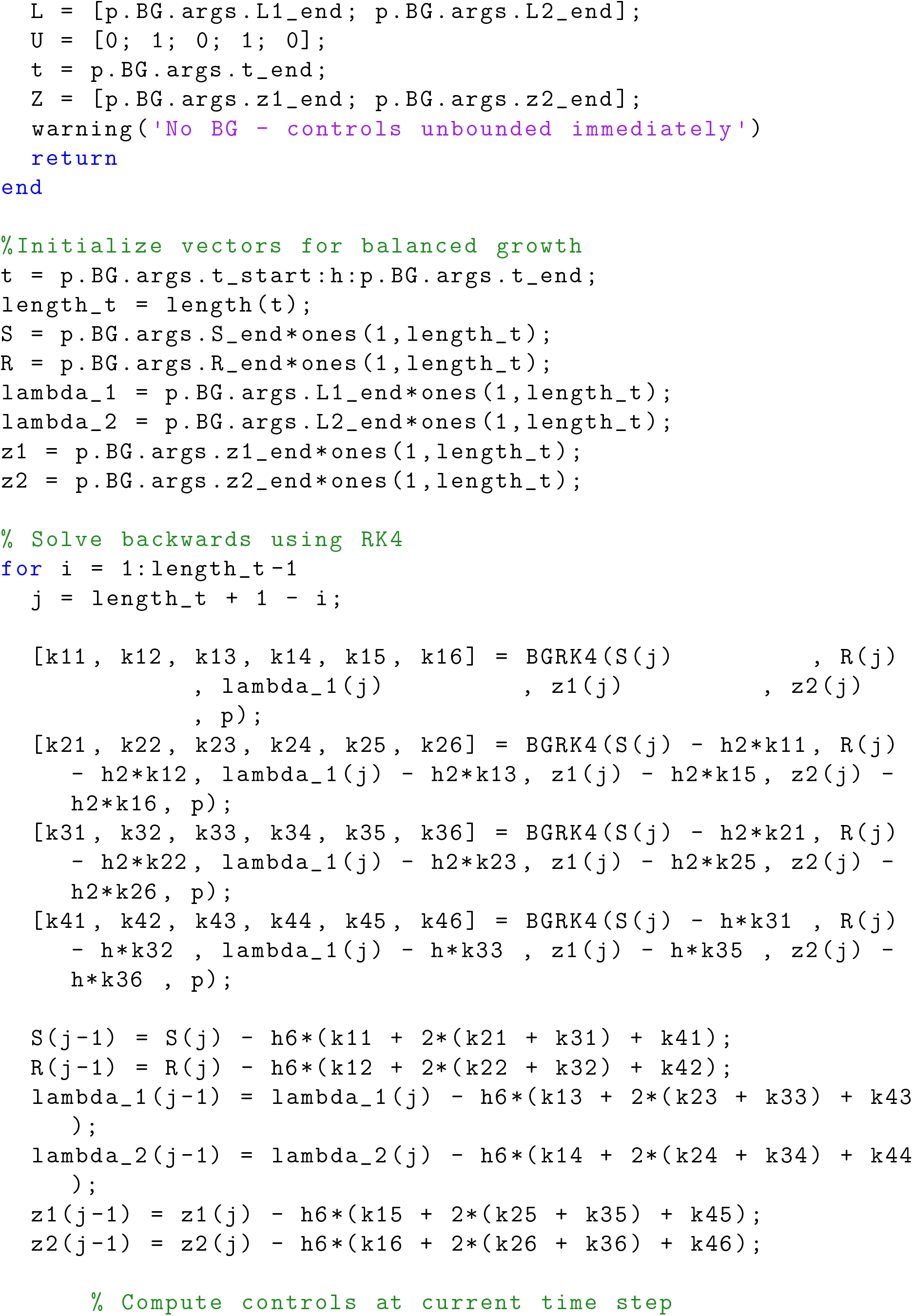
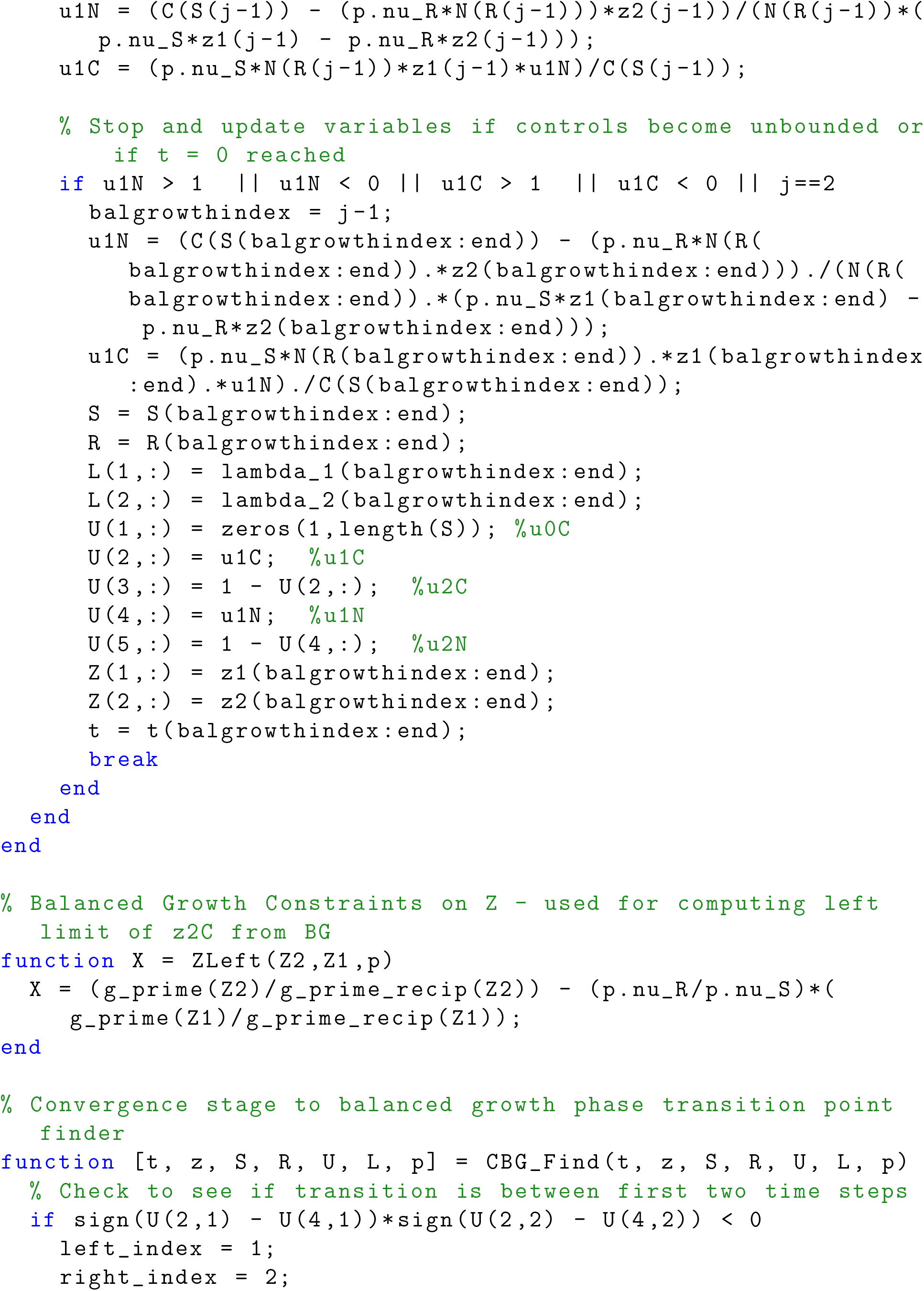
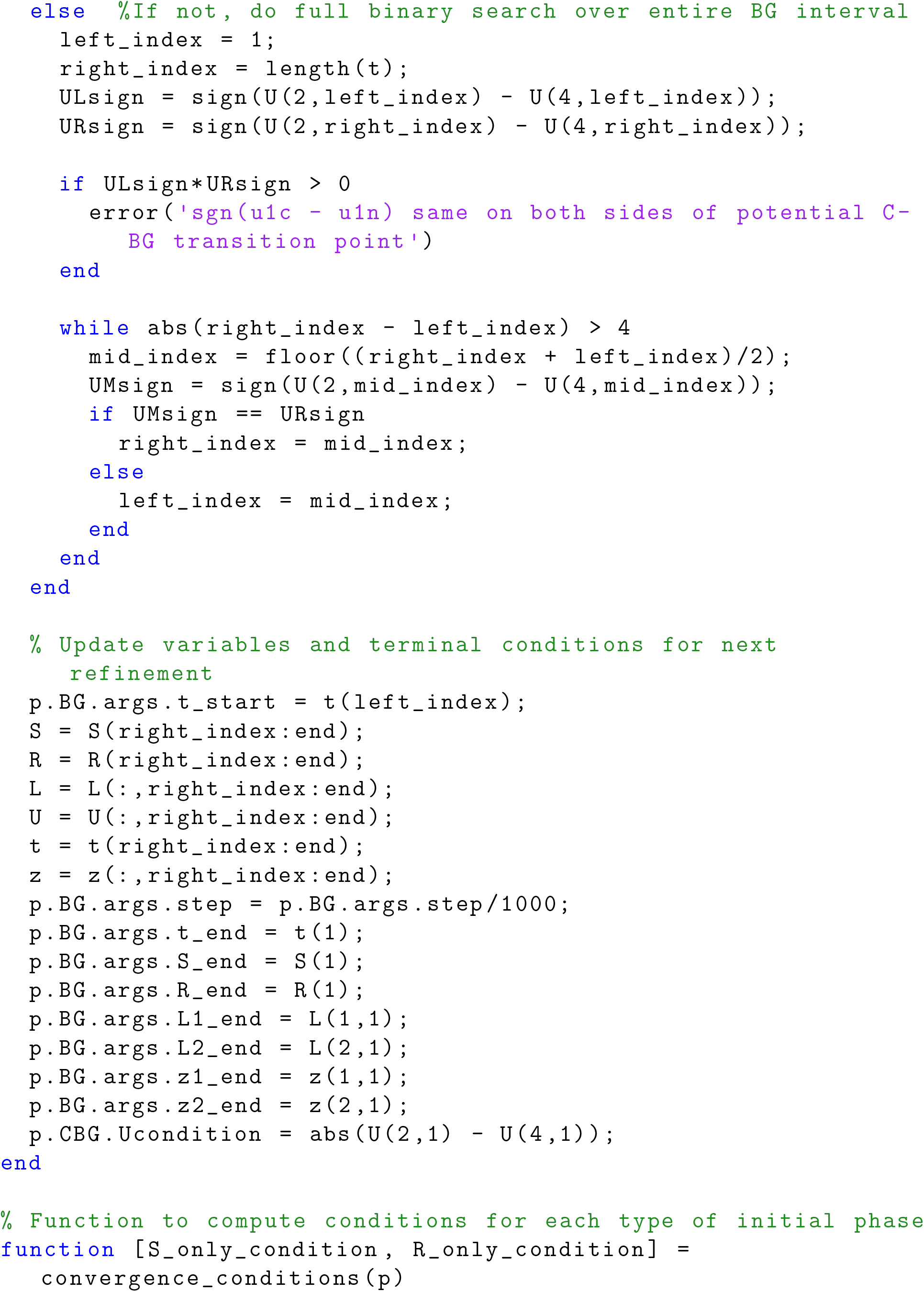
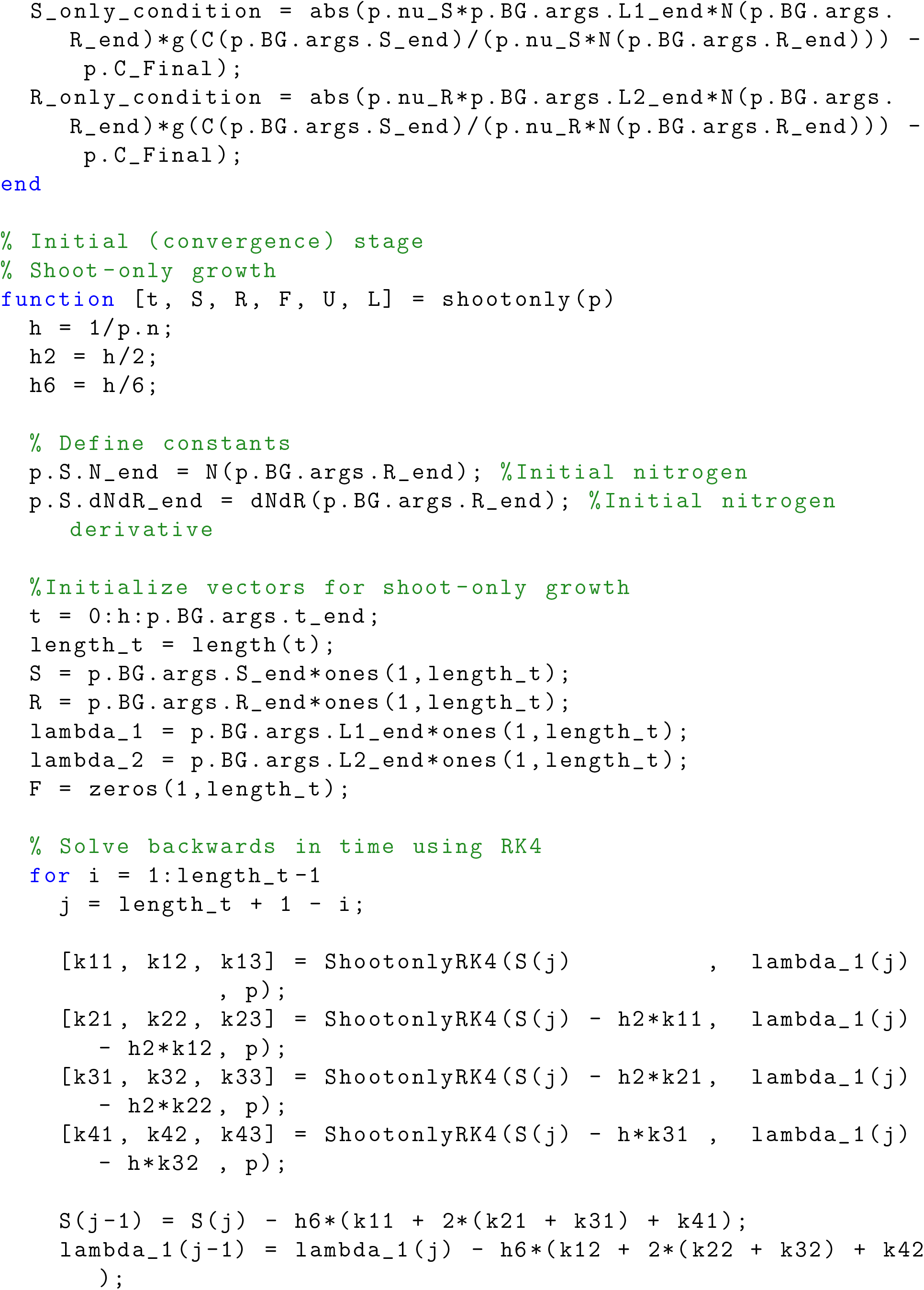
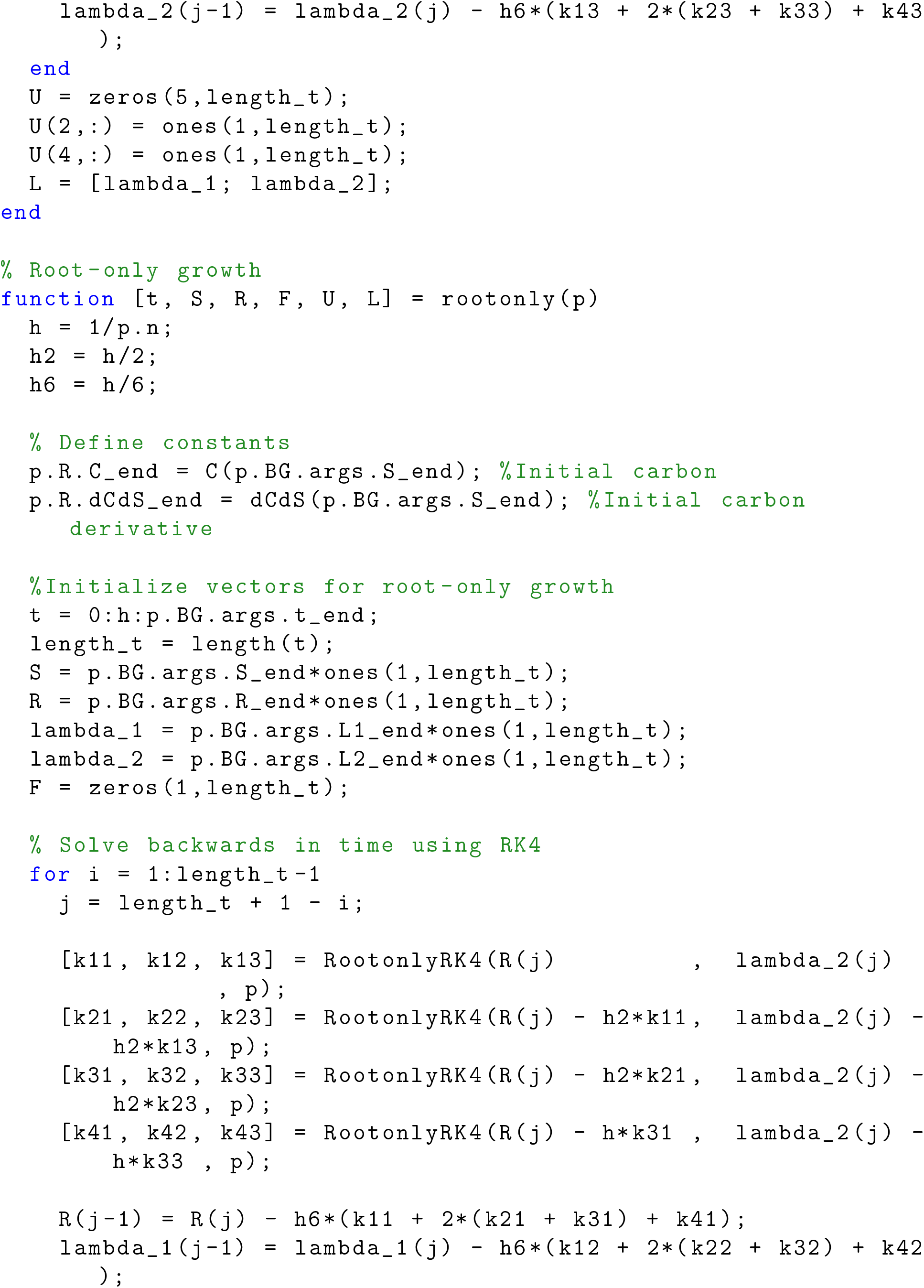
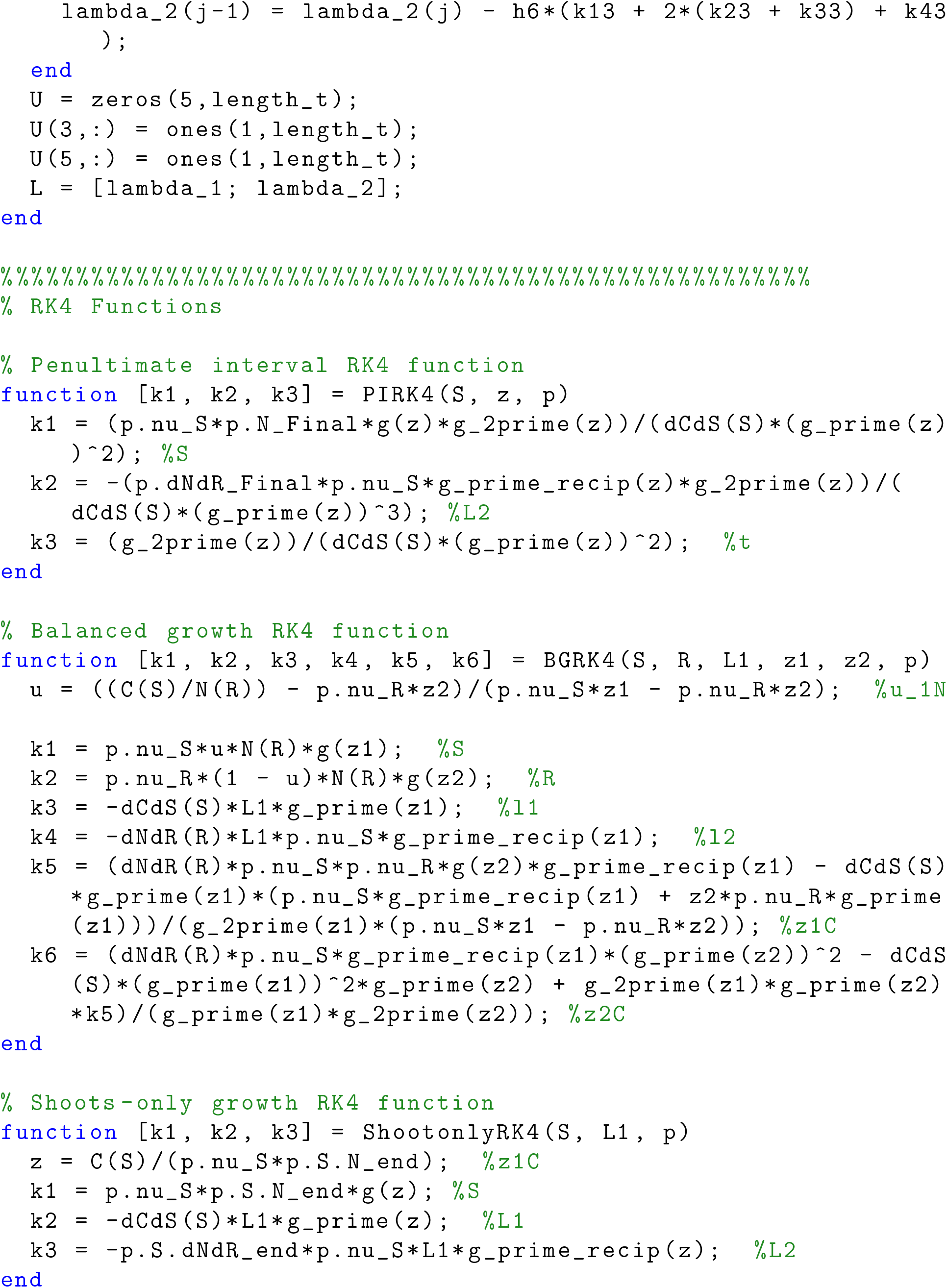
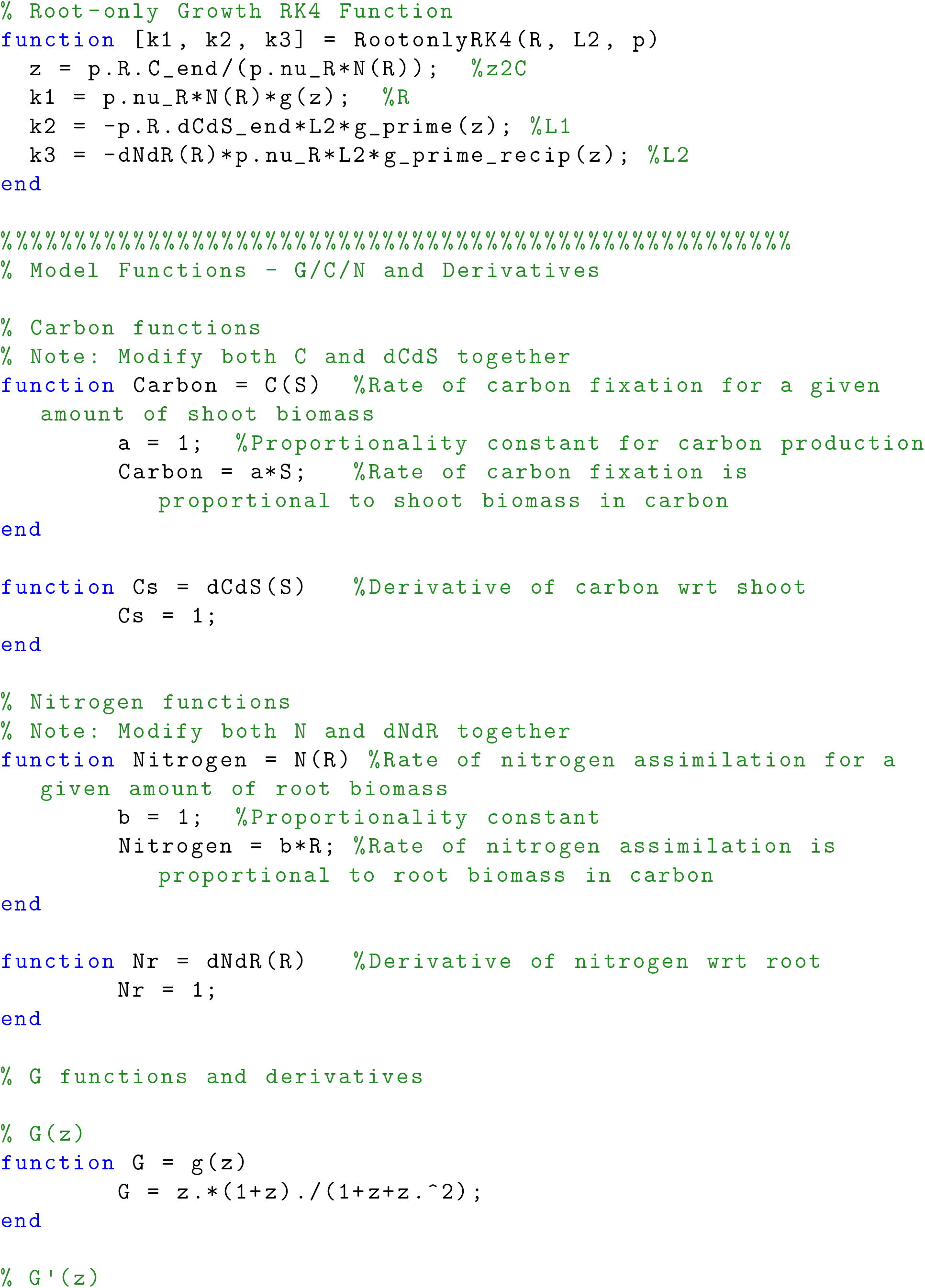
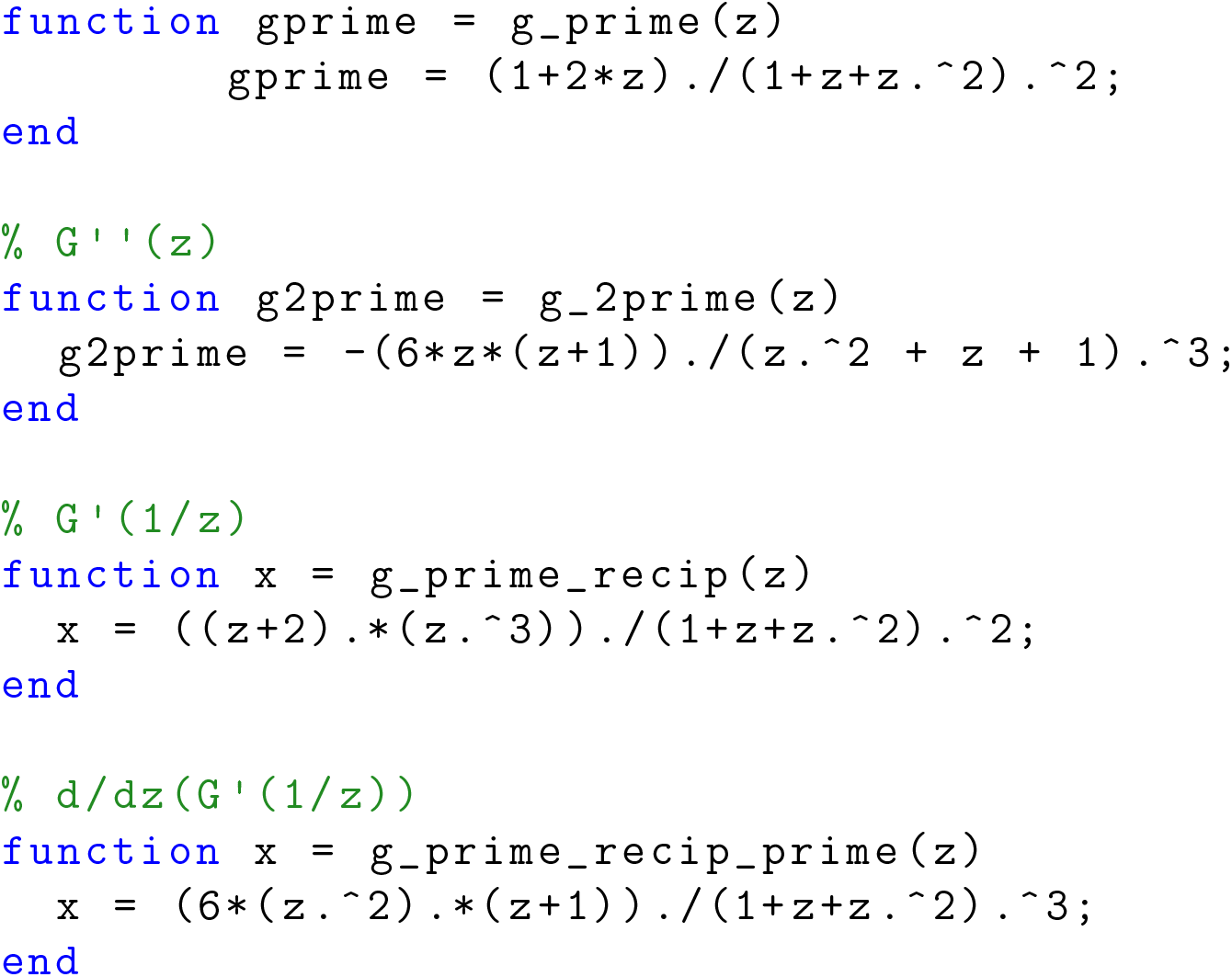

